# Initiator and Elongator tRNA Recognition Mechanism in *Mycobacterium tuberculosis* Methionyl-tRNA Synthetase

**DOI:** 10.1101/2025.05.27.656506

**Authors:** Shivani Thakur, Rukmankesh Mehra

**Affiliations:** Department of Chemistry, Indian Institute of Technology Bhilai, Durg 491002, Chhattisgarh, India; Department of Bioscience and Biomedical Engineering, Indian Institute of Technology Bhilai, Durg 491002, Chhattisgarh, India

**Keywords:** initiator tRNA, elongator tRNA, methionyl-tRNA synthetase, mechanism, *Mycobacterium tuberculosis*, molecular dynamics simulations

## Abstract

The protein synthesis is an essential target for anti-tubercular drug design. Methionyl-tRNA synthetase (MetRS) typically plays a role in elongation and the initiation of protein synthesis. Molecular recognition of the CAU anticodon of tRNA by MetRS in the two processes is a crucial step. Until now, no known experimental structures for *Mycobacterium tuberculosis* (*Mtb*) show this binding. We therefore modeled the *Mtb* MetRS complexes with initiator and elongator tRNAs to find their differential binding mechanism during molecular dynamics simulations of 6 µs. We found that the elongator tRNA binding was stable with the protein, while the initiator tRNA binds transiently, with major intra-tRNA interactions maintained in both. This could be due to fast initiator tRNA charging in contrast to the elongator tRNA, which could take more time to charge. tRNA interacts with the MetRS active site and anticodon domain. The electrostatic attractions between tRNA and the protein’s catalytic domain possibly caused its charging with methionine. The repulsive and attractive forces between tRNA and the protein’s connective peptide domain and KMSKS loop triggered the opening of the binding pocket, causing the reaction and the product release. At the same time, tRNA’s strong binding to the protein’s anticodon domain facilitated this reaction. These events show the possible pathway of tRNA charging. tRNA formed salt-bridges with the positively charged Arg and Lys, whereas the negatively charged Asp and Glu caused repulsive binding. In brief, this study provided a plausible mechanism for initiator versus elongator tRNA recognition by *Mtb* MetRS.

---

Protein synthesis is a crucial target for developing new anti-tubercular agents. Aminoacyl-tRNA synthetase charges tRNA with the amino acid, which is then transferred to the site of protein synthesis on ribosomes. Methionine embarks on the initiation of translation and is also added during the elongation of the amino acid chain. This makes aminoacyl-tRNA synthetase linked to methionine charging (MetRS) a crucial enzyme for targeting^1,2^ and central to research on the origin of life.^3–5^

Methionyl-tRNA synthetase (MetRS) charges two tRNAs, initiator and elongator, associated with the initiation and elongation of translation, respectively (**Figure 1a, b**). *Mycobacterium tuberculosis* (*Mtb*)^6^ comprised three methionyl-tRNA encoding genes: *metU* for initiator tRNA, and *metT* and *metV* for elongator tRNA (**Figure S1**). *metV* undergoes post-transcriptional modifications and charges isoleucine instead of methionine.^2,7^ The initiator and elongator tRNA (77 nucleotides in length) corresponding to *metU* and *metT* participate in binding to MetRS.^1,2^

**Figure 1.**
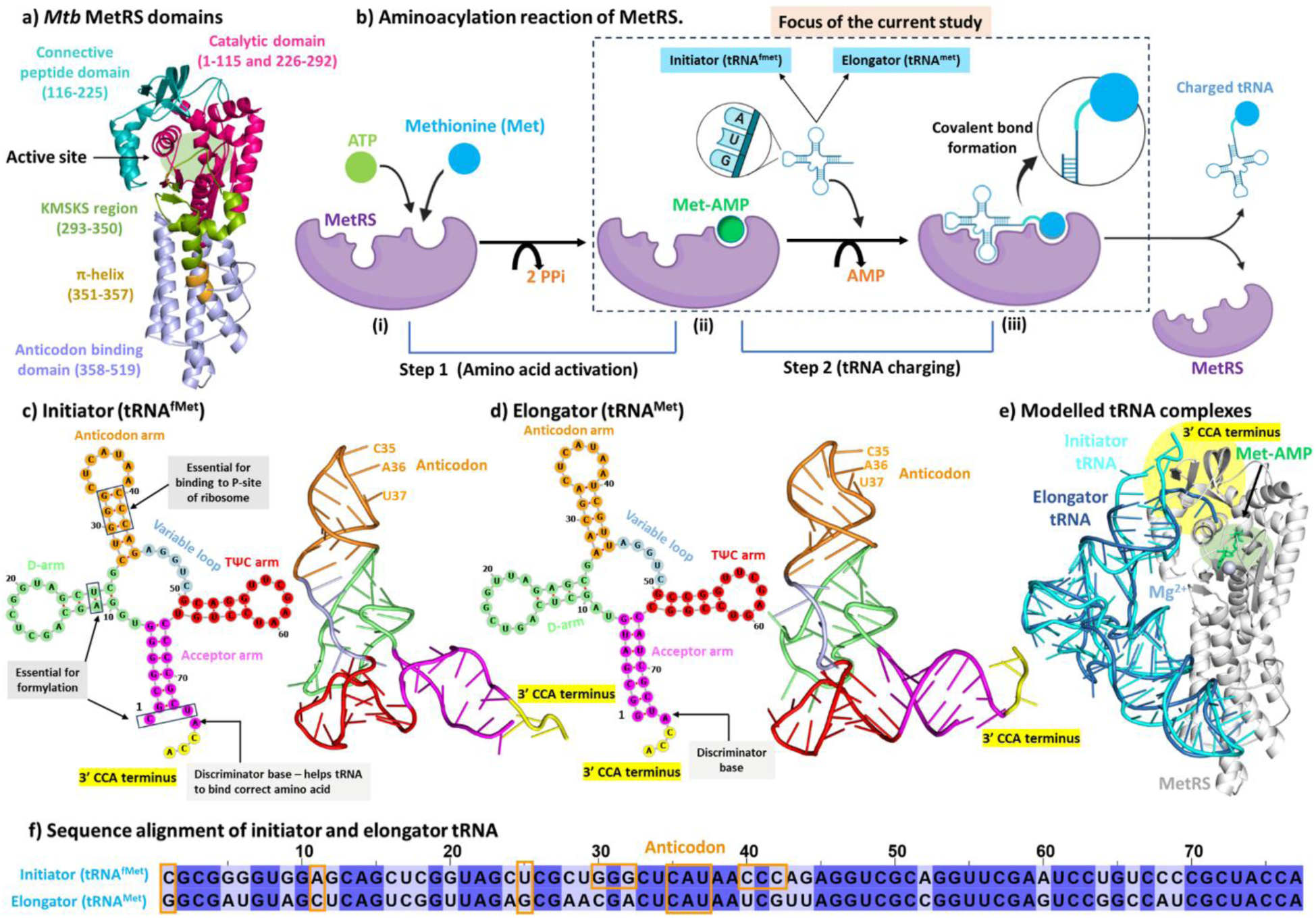
MetRS and tRNA in the aminoacylation reaction. **(a)** *Mtb* MetRS structure showing catalytic domain (pink, 1-115 and 226-292), connective peptide (CP) domain (dark cyan, 116-225), KMSKS domain (green, 293-350), π-helix region (brown, 351-357) and anticodon domain (light blue, 358-519). **(b)** Steps involved in aminoacylation reaction. The cloverleaf and L-shaped model of **(c)** the initiator and (d) the elongator tRNA of *Mtb* **(e)** Superimposed structures of modelled protein-tRNA complexes (initiator and elongator). **(f)** Nucleotide sequence alignment of both tRNAs, highlighting distinct nucleotides and conserved anticodon (orange). For consistency, nucleotides are numbered sequentially from 1 to 77 (may differ from the literature).

MetRS performs a two-step reaction (**Figure 1b**).^8,9^ In the first step, the substrates methionine and ATP bind to the protein and form the methionine-AMP (Met-AMP) complex. Subsequently, tRNA binds to protein via its anticodon and 3’ CCA (acceptor arm) end at two protein sites, leading to tRNA charging with methionine, which is released from the protein.

*Mtb* MetRS is 519 amino acids in length, divided into catalytic domain (1-115 and 226-292) connective peptide (CP) domain (116-225), KMSKS loop region (293-350), π-helix region (351-357) and anticodon binding domain (ABD, 358-519) (**Figure 1a**).^10^

*Mtb* initiator and elongator methionine tRNA form a typical clover-leaf structure (**Figure 1c, d**) of 77 nucleotides having acceptor arm (1-7 and 67-77), D-arm (8-27), anticodon arm (28-44), variable loop (45-49) and TΨC arm (50-66).^11^

The substrates (methionine and ATP) and the 3’ CCA terminal of tRNA bind to the protein’s catalytic domain, CP domain, and KMSKS region.^10,12,13^ The tRNA anticodon arm also interacts with the ABD site of MetRS for anticodon (CAU) recognition by protein. The tRNA interactions at two protein sites lead to tRNA charging. The two tRNAs (initiator and elongator) share a sequence identity of 67.9% (**Figure 1f**), suggesting the possibility of differential recognition mechanisms and functional consequences.

In the current work, we performed a detailed study on the molecular recognition of initiator versus elongator tRNA by *Mtb* MetRS using molecular dynamics (MD) simulations. We prepared the 3D states of these complexes and identified the structure-function relations of tRNA, protein, and Met-AMP. These relations helped elucidate the distinct structural and binding features between initiator and elongator tRNA complexes and the plausible tRNA recognition mechanisms.

## RESULTS AND DISCUSSION

### tRNA sequence analysis

*Mtb* contains three genes - *metU*, *metT,* and *metV* that encode tRNA with CAU anticodon to read AUG codon.^14^ The genes *metU* and *metT* encode for initiator (tRNA^fMet^) and elongator (tRNA^Met^) tRNA, respectively, to process methionine (**Figure 1c-f** and **S1a**). While the post-transcriptional modification of tRNA encoded by *metV* leads to an AUA anticodon that processes isoleucine (tRNA^Ile2^) instead of methionine.^2,7^

These three sequences (tRNA^fMet^, tRNA^Met^, and tRNA^Ile2^) were compared using alignment. Since both tRNA^Met^, and tRNA^Ile2^ are elongator sequences, they share a high similarity. BLASTN^15^ showed 81.97% identity (with 79% query coverage) and EMBOSS^16^ revealed 77.5% identity (with full query coverage) (**Table S1, Figure S1d**). These results suggest a high nucleotide conservation between tRNA^Met^ and tRNA^Ile2^.

In contrast, tRNA^fMet^ (initiator) showed no significant local similarity with either tRNA^Met^ or tRNA^Ile2^ (elongators) using BLASTN (**Table S1**), indicating functional differences. However, EMBOSS revealed moderate identity between tRNA^fMet^ and tRNA^Met^ (67.9%), and between tRNA^fMet^ and tRNA^Ile2^ (64.3%), suggesting partial conservation across the complete sequences (**Table S1**). We observed the conserved nature of the anticodon loop (_33_CUCAUAA_39_; **Figure S1b-d**) in the three tRNAs. This means that the anticodon loop is essential for tRNA recognition.

### The protein in the elongator state exhibited higher conformational dynamics

We prepared two *Mtb* tRNA-MetRS complexes: one with initiator (tRNA^fMet^) and the other with elongator (tRNA^Met^) tRNA (**Figure 1e, S2,** details in Methods). In both complexes, intermediate (Met-AMP) and Mg^2+^ were present.

MD simulations of both complexes were performed in triplicate for 1000 nanoseconds (ns) each, for a total of 6 microseconds (µs; 2 systems × 3 µs each). The primary analysis was performed on the full trajectory involving root mean square deviation (RMSD), radius of gyration (R_g_), and total solvent-accessible surface area (SASA). The protein backbone RMSD showed greater conformational flexibility of the elongator protein than the initiator (**Figure S3a**). Their corresponding average values (of three MD runs) were 0.40 and 0.36 nm (**Figure 2a**).

**Figure 2.**
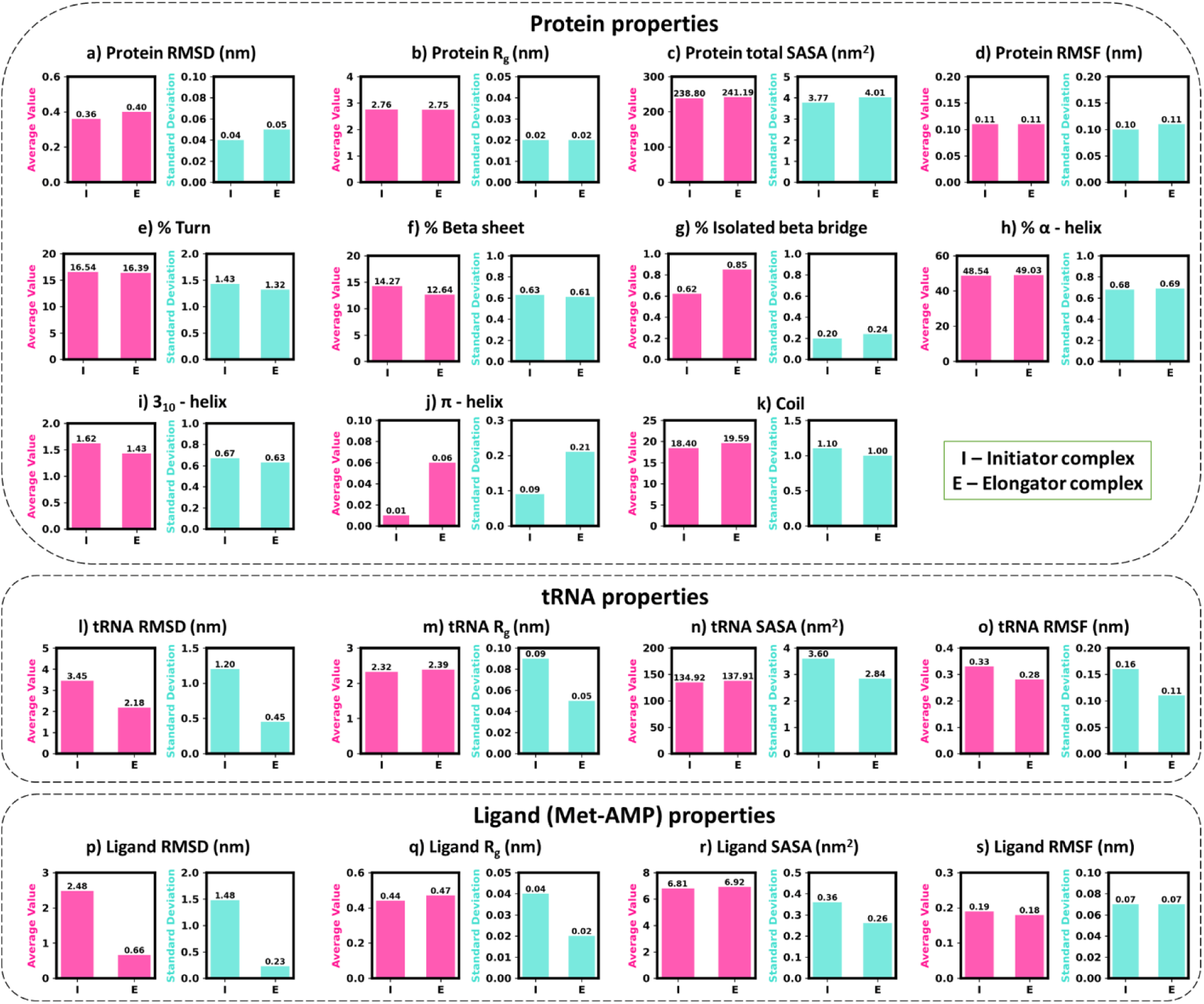
Comparisons of the simulated average properties (in magenta) of the three independent MD runs and their standard deviation (in cyan) in initiator (I) versus elongator (E) complexes. Protein properties: **(a)** RMSD, **(b)** R_g_, **(c)** total SASA, **(d)** RMSF, and **(e-k)** secondary structures. tRNA properties: **(l)** RMSD, **(m)** R_g_, **(n)** SASA, **(o)** RMSF. Ligand (Met-AMP) properties: **(p)** RMSD, **(q)** R_g_, **(r)** total SASA, **(s)** RMSF.

Despite some structural fluctuations, the protein’s R_g_ stayed consistent in both complexes (**Figure S3b**). This suggests that the protein’s overall size and compactness remained unchanged throughout the simulations (**Figure 2b**).

However, a modest increase in solvent accessibility was observed for the protein in the elongator complex, with a higher average SASA of 241.19 nm², compared to the initiator complex (238.80 nm², **Figures S3c, 2c**). Both hydrophobic (195.40 nm²) and hydrophilic (288.25 nm²) SASA of protein were greater in the elongator complex than in the initiator complex (193.80 nm² and 286.66 nm², respectively), indicating that the conformational changes led to increased exposure of protein surface residues to the solvent (**Figure S4**).

### Dominant motions observed in the N-terminal, CP domain, KMSKS loop, and C-terminal

Free energy landscape principal component analysis (FEL-PCA) was performed to investigate the dominant motions of the protein in the initiator and elongator tRNA complexes. The first two eigenvectors (EV1 and EV2) accounted for a significant portion of the total motion (details in **Supplementary Section S1** and **Figure S5**).

We identified the conformationally active protein regions using porcupine plots of EV1 and EV2 for six simulations (**Figures S6 and S7**). The dominant motions were observed in key functional regions, including the catalytic domain (residues 9-21), CP domain (116-155), KMSKS loop (296-306), and C-terminal, suggesting enhanced mobility around the active site.

The residue-wise root-mean-square-fluctuation (RMSF) (**Figures 2d** and **S8**) and the ratio of intra-molecular contacts (**Figure S9**) of protein support the observations from the porcupine plots. The N- and C-terminals, CP domain, and KMSKS loop consistently showed higher flexibility and intra-molecular contacts. The CP domain displayed greater RMSF and increased intra-molecular contacts in the elongator than in the initiator complex. This might be due to the conformational changes driven by prominent interactions of this region with the elongator tRNA.

In contrast, the KMSKS loop (299-303) exhibited comparatively higher flexibility and contacts in the initiator complex. This indicates that tRNA interactions at this region might cause the opening of the KMSKS loop surrounding the active site.

### Decrease in β-sheet content found in elongator protein

The analysis of secondary structural elements of proteins over the last 500 ns of the simulations showed differential patterns between the initiator and elongator states (**Figures 2e-k** and trajectories in **Figure S10**). Specifically, β-sheet content was higher in the initiator (14.27%) compared to the elongator (12.64%), while the isolated β-bridge content was lower (0.62% in the initiator versus 0.85% in the elongator). This indicates the formation of comparatively stable structural elements in the initiator protein. Minor differences were observed for other secondary structures. The percentage of 3_10_-helix was higher in the initiator protein, whereas the π-helix and coil content were higher in the elongator.

The CP domain residues between 120-125 showed a predominant regular β-sheet in the initiator protein, while this region majorly converted to a flexible coil structure in the elongator (indicated by dashed line, **Figure S11**).

Thus, the initiator complex encountered a higher proportion of β-sheets contributing to the comparatively greater stability of the protein, with major differential content observed in the CP domain (120-125). The CP domain directly binds with tRNA via its CP1 knuckle region (residue 120-140).^2,12,17^ We observed above that this domain fluctuated highly and formed more intra-molecular contacts in both initiator and elongator states, with higher fluctuations and contacts in the elongator protein. This means that the longer binding of the CP domain of elongator protein with tRNA led to the conversion of a stable β-sheet to a coil structure, possibly due to the continuous repulsive tRNA-protein interactions at this region (discussed later). While in the initiator protein, such repulsive tRNA-protein interactions lasted for a shorter duration and thus formed a more stable β-sheet structure.

### Stable binding of elongator tRNA and transient binding of initiator tRNA

The primary analysis of tRNA (over full trajectory) indicated higher conformational flexibility of the initiator tRNA than the elongator (**Figures 2l** and **3a**). The initiator tRNA showed the average RMSD (for the three runs) of 3.45 nm compared to 2.18 nm for elongator. This suggests that elongator tRNA maintained a stable conformation while initiator tRNA formed transient interactions with protein.

**Figure 3.**
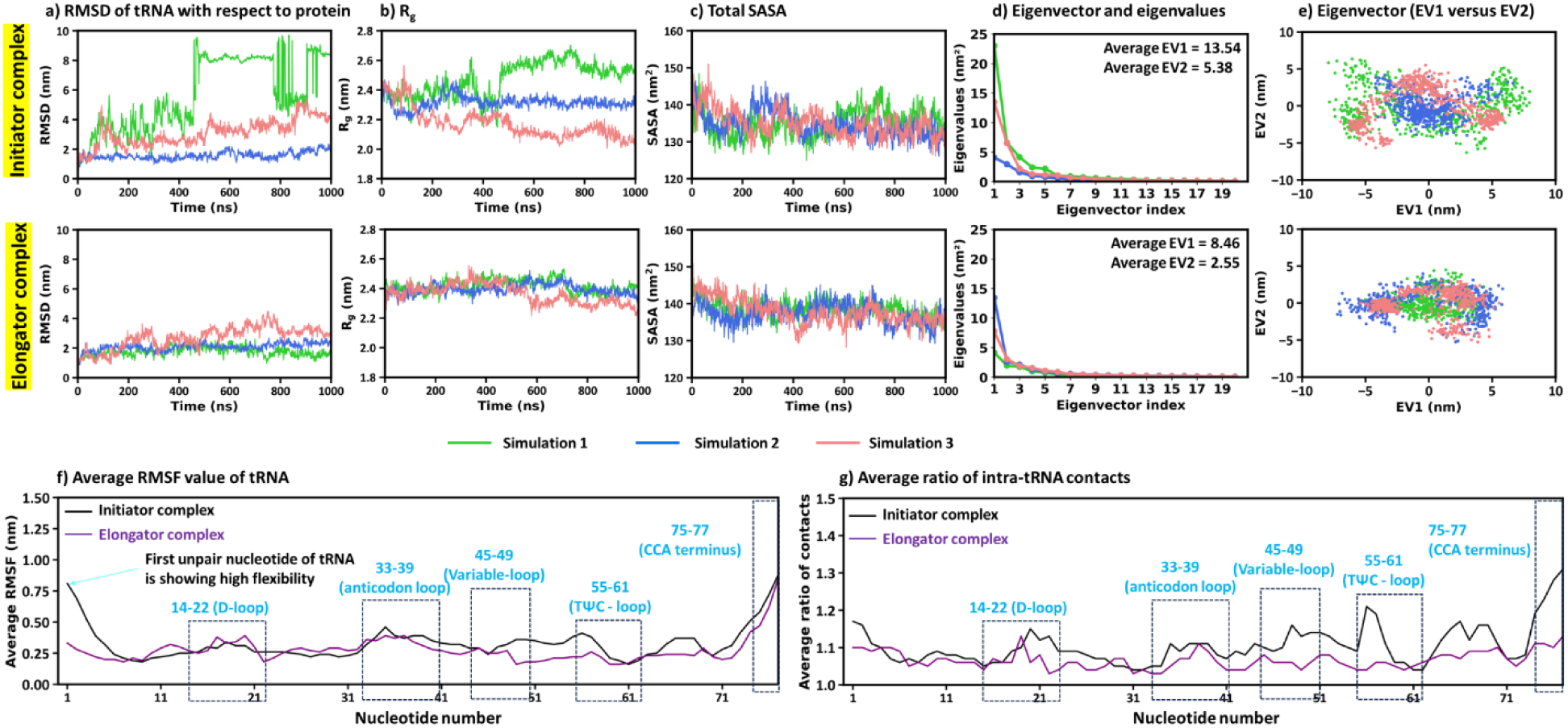
Comparisons of the simulated properties of the initiator and elongator tRNAs. **(a)** RMSD. **(b)** R_g_. **(c)** Total SASA. **(d)** Eigenvectors versus eigenvalues plots. **(e)** 2D projections along EV1 and EV2. **(f)** Average RMSF values of three runs, showing flexibility of nucleotides. **(g)** Average ratio of intra-tRNA contacts of each nucleotide. The contact ratio is the total number of contacts divided by their mean value during simulation.

Though the R_g_ of the initiator (2.32 nm) was slightly lower than that of the elongator (2.39 nm, **Figure 2m**), the initiator tRNA exhibited a high variability (**Figure 3b**). SASA also revealed a similar trend (135 nm^2^ of initiator versus 138 nm^2^ for elongator, **Figure 2n and 3c**), with a higher variation for initiator tRNA.

These findings indicate that the initiator tRNA binds transiently, with a high variability in size and solvent exposure. The elongator tRNA maintained stable conformations in a slightly extended form during simulations.

### Confirmation of comparatively higher motion of initiator tRNA

To confirm the differential motion of initiator versus elongator tRNA, FEL-PCA of tRNA dynamics was performed. This analysis showed higher motion and a broader conformational landscape of the initiator tRNA. The average EV1 and EV2 values (of three runs) were 13.54 and 5.38 nm^2^ for initiator tRNA, while the corresponding values for elongator tRNA were 8.46 and 2.55 nm^2^ (**Figure 3d**). The EV1 versus EV2 scatter plots showed a wider conformational space of the initiator tRNA (**Figure 3e**). The elongator tRNA, on the other hand, spanned a comparatively confined space, suggesting its lesser motion.

### Initiator tRNA exhibited higher flexibility and intra-molecular contacts

RMSF analysis of nucleotides showed that the initiator tRNA typically displayed higher fluctuations (**Figures 2o**, **3f, and S12**), which could relate to its higher ratio of intra-molecular contacts (**Figure 3g and S13**) compared to elongator tRNA. The CCA end (75-77) showed higher dynamics in both initiator and elongator tRNA. The first nucleotide (C1) of the initiator tRNA displayed increased flexibility. In addition, a higher contact ratio was observed for nucleotides in the TΨC (55-61) and variable loops (45-49) of the initiator tRNA (**Figure 3g**).

These results lead to the concept that initiator tRNA in *Mtb* might bind for a shorter time and perform reaction quickly, which could be due to its necessity to initiate protein synthesis. However, the elongator tRNA remained stably bound to MetRS and could require more time to charge tRNA, which is involved in protein elongation. The respective short and long lifetimes of the initiator and elongator tRNA complexes perfectly correlated with their experimental association constant (K_a_) values of 16±2 and 57±10 µM^-1^ in *Escherichia coli*.^18^ A low K_a_ indicates short lifetime. The quick reaction also possibly relates to the rapid formylation of the initiator tRNAs, which is critical for its preferential binding to the ribosome’s P-site and prevents other tRNAs.^19^

### Ligand spanned a shorter period in the active site of the initiator complex

The primary analysis of the ligand over 1000 ns of the three runs revealed a highly unstable nature of Met-AMP in the initiator complex relative to the elongator (**Figure 2p** and **S14a**). Met-AMP remained in the protein’s binding pocket for a shorter time in the initiator complex. In contrast, the ligand occupied the active site throughout the simulations in the elongator (**Figure S14a**). Met-AMP’s average RMSD (of three runs) was 2.48 and 0.66 nm in initiator and elongator complexes, respectively (**Figure 2p**).

The higher ligand motion was supported by a higher variability of its R_g_ in initiator complexes. In contrast, the ligand R_g_ was consistently maintained in the elongator state during simulations (**Figures 2q** and **S14b**). Similarly, the ligand SASA showed more variability in the initiator complex (**Figures 2r** and **S14c**).

These observations support the early release of the ligand from the protein’s active site in the initiator complex, leading to its size and solvent exposure variability. In contrast, the elongator complex maintained a comparatively stable ligand state in the protein’s binding pocket.

The observed Met-AMP instability in the initiator was strongly related to the early release of the initiator tRNA from the protein compared to the elongator tRNA. The tRNA-protein interactions appear to trigger the opening of the protein’s active site, causing the ligand release. This complete event was observed on a shorter timescale in the initiator complex. The elongator complex also revealed a similar tendency; however, the stable tRNA binding to protein prevented the complete release of Met-AMP from the active site. This shows that tRNA plays a crucial role in stabilizing Met-AMP binding.

This means that the reaction of the initiator tRNA could be fast on a timescale supported by its transient binding, which could also cause an early release of the ligand. However, the elongator tRNA reaction with MetRS could be comparatively slower, supported by its binding for a substantial time and comparatively more ligand stability at the active site. These are supported by the higher experimental K_m_ values for initiator tRNA (3.7 µM) compared to elongator (2.6 µM) in *E. coli*.^18^

### Ligand PCA and RMSF confirmed the dynamical correlation with tRNA

FEL-PCA of the ligand showed higher average eigenvalues of EV1 and EV2 (of three runs) in the initiator ligand (1.40 and 0.73 nm) than in the elongator (0.78 and 0.32 nm) (**Figure S15a**). Consistent with this, the EV1 versus EV2 plot showed a wider distribution of the conformational landscape for the initiator ligand than for the elongator (**Figure S15b**). Moreover, the RMSF analysis of ligand atoms clearly showed more movements of the initiator ligand than the elongator (**Figures S16** and **2s**).

These observations correlated excellently with the dynamic landscape of tRNA, i.e., EV1 and EV2 were higher for both the initiator tRNA and ligand compared to the elongator counterparts. Similarly, the initiator tRNA and ligand showed higher RMSF of atoms than the elongator ones.

In addition, we performed FEL-PCA of the complete initiator versus elongator complexes (i.e., with protein, tRNA, and Met-AMP together). The major contribution to complex mobility was from tRNA and ligand molecules (details in **Supplementary Section S3** and **Figure S17**).

### Intra-molecular hydrogen bonds increased in protein while reduced in tRNA of initiator complex

The numbers of intra-molecular hydrogen bonds within protein and tRNA were analyzed for the final 500 ns trajectories (**Figure S18**). Initiator complex formed a higher number of intra-protein and a reduced number of intra-tRNA hydrogen bonds. The average (of three runs) intra-protein hydrogen bonds were 337.91 in the initiator complex and 307.03 in the elongator. The corresponding average intra-tRNA hydrogen bonds were 103.85 and 115.51, respectively.

The initiator protein achieved a more stable conformation than the elongator due to the higher number of hydrogen bonds. This relates well to the protein RMSD (discussed earlier). In contrast, tRNA achieved a more open conformation in the initiator due to the decreased hydrogen bonds, which correlated with the increased RMSD of tRNA in this complex.

These findings suggest that due to the higher motion of tRNA and its open conformation in the initiator, the tRNA was not bound for longer to the protein, causing a comparatively stable protein state due to fewer tRNA-protein interactions. In comparison, longer tRNA-protein contacts in the elongator complex influenced more conformational changes to the protein.

### Solvent interactions showed a distinguishing effect for tRNA and ligand

The number of hydrogen bonds of water with protein, tRNA, and ligand (Met-AMP) was analyzed for the last 500 ns (**Figure S19**). The protein-water hydrogen bonds were slightly higher in the initiator complex (**Figure S19a**). The average numbers (of three runs) were 943.48 in initiator versus 934.71 in elongator complexes. This might be due to the transient binding of the initiator tRNA and its early release from the protein’s primary binding site.

The tRNA-water hydrogen bond trajectories showed higher variations and lower average numbers in the initiator complex (**Figure S19b**). The average numbers were 519.62 and 599.03 for initiator and elongator complexes, respectively. This suggests the possibility of initiator tRNA binding to distinct parts of the protein and its comparatively compact conformation (indicated by R_g_) that reduced the tRNA-water contacts.

Met-AMP formed more hydrogen bonds with water in the initiator complex (average = 11.70) than the elongator (average = 10.48) (**Figure S19c**). This was consistent with the higher ligand dynamics in the initiator complex that caused the early release of the ligand from the active site to the solvent.

### tRNA-protein interactions and plausible tRNA recognition mechanism

The tRNA-protein interactions were analyzed for the equilibrated trajectories (final 500 ns, **Figures S20** and **4a-b**). The initiator complex showed fewer hydrogen bonds (average = 8.5) than the elongator state (10.66). Similarly, the salt-bridges were less in the initiator (4.80) than in the elongator (5.22) state. In addition, minor changes in other interactions were observed; π-π and π-cation contacts were more for the initiator complex (**Figure S20c-d**), whereas π-anion contacts were more for the elongator (**Figure S20e**).

The binding affinity (ΔG_bind_) between tRNA and protein was analyzed for the last 500 ns simulations (**Figures 4c** and **5a**). Notably, the elongator tRNA showed a stronger binding to the protein (ΔG_bind_ = -53.89 kcal/mol) compared to the initiator (ΔG_bind_ = -13.16 kcal/mol).

**Figure 4.**
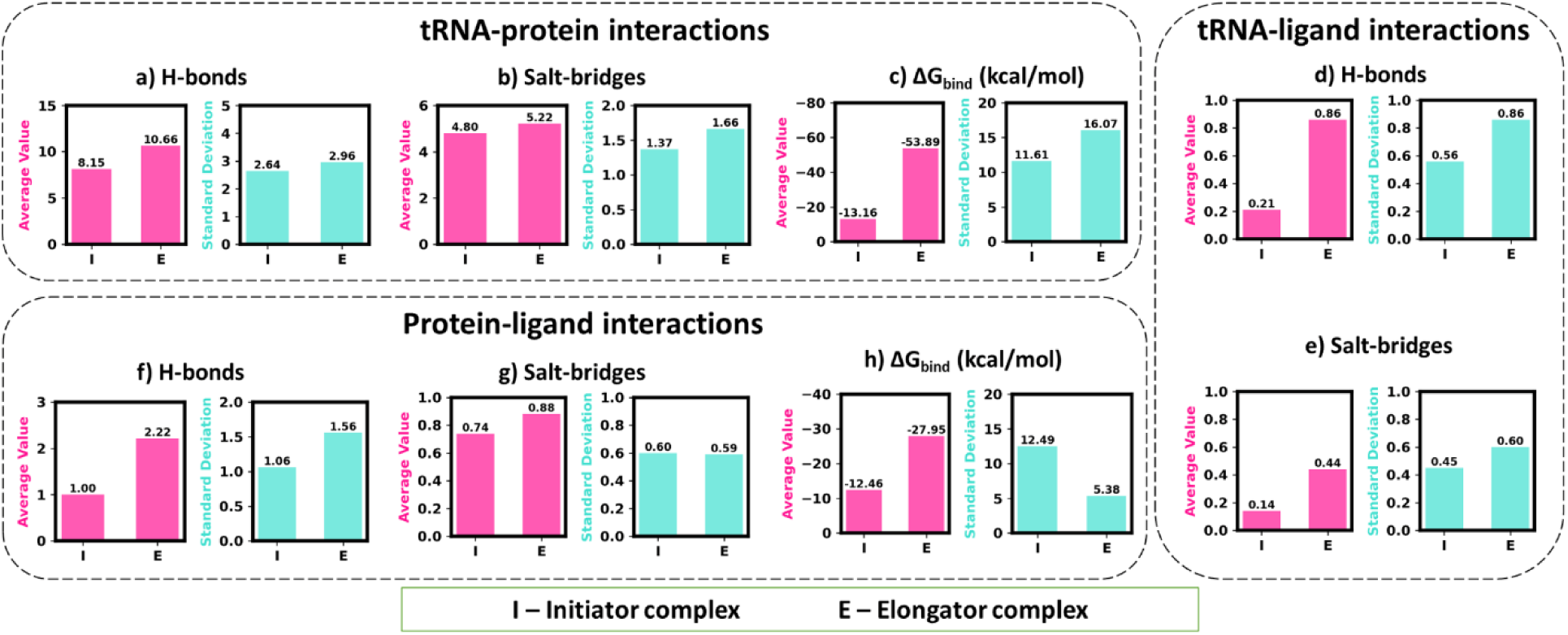
Average binary interactions for three MD simulations in the initiator (I) and elongator (E) complexes and their standard deviation values. Comparison of hydrogen bond counts, salt-bridge counts, and binding free energy (ΔG_bind_) values observed due to interactions between **(a-c)** tRNA and protein, **(d-e)** tRNA and ligand, and **(f-h)** protein and ligand.

**Figure 5.**
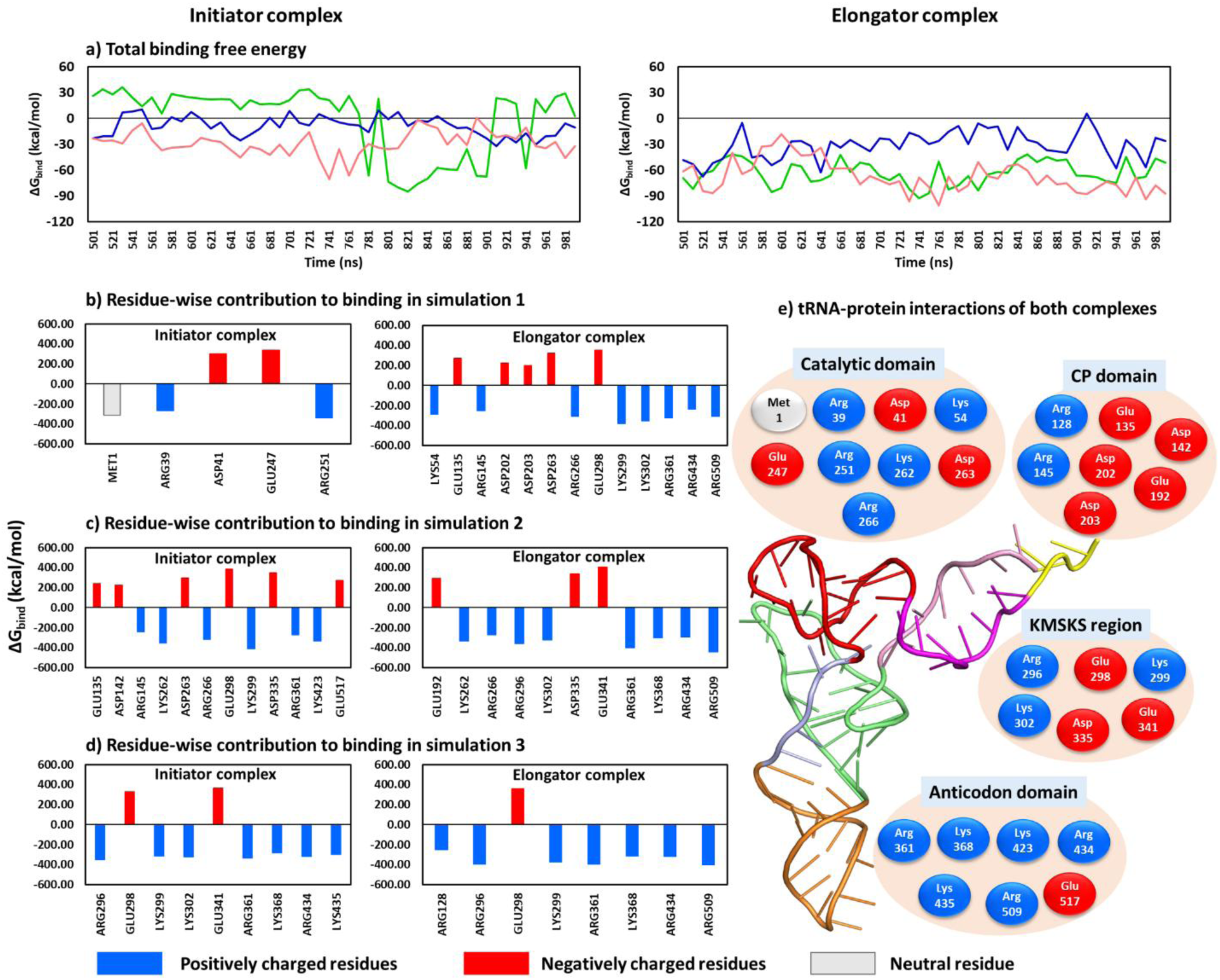
ΔG_bind_ of tRNA-protein interactions in initiator and elongator complexes. **(a)** ΔG_bind_ over three simulations. **(b-d)** Protein residue-wise impact on ΔG_bind_ in each run. A cutoff of ± 50 kcal/mol was used to define the prominent contributions. **(e)** tRNA-interacting residues divided into protein regions. Blue, red, and gray represent positive, negative, and neutral residues.

These findings indicate that the hydrogen bonding and electrostatic interactions were prominent in both complexes, with decreased numbers for the initiator state due to the transient tRNA binding to the protein. These contacts led to the stronger binding of elongator tRNA to protein, increasing the elongator tRNA’s lifetime in the protein’s active site. In comparison, the weaker binding caused early release of the initiator tRNA. These results correlated well with the experimental study on aminoacylation of the initiator and elongator tRNA by *E. coli* MetRS,^18^ which showed decreased binding affinity of the initiator tRNA for MetRS compared to the elongator.

The analysis of protein residues contributing to the binding showed that the favorable contacts were formed by arginine and lysine, while the unfavorable contacts occurred with aspartate and glutamate residues (**Figure 5b-e**). The main interactions occurred with the catalytic domain (1-115 and 226-292), CP domain (116-225), KMSKS loop region (293-350), and anticodon domain (358-519) (**Figure 5e**). The catalytic domain primarily formed electrostatic attractions (blue colored residues in **Figure 5e**), contributing to tRNA binding to the active site. This event might help in the tRNA charging with methionine.

The CP domain contributed mainly electrostatic repulsive forces with tRNA via its negatively charged residues (five residues, red color in **Figure 5e**). Two residues, Arg128 and Arg145, formed favorable contacts. The KMSKS loop (299-303) is conserved in MetRS^10^ and is present at the entry of the active site (**Figure 1a**), forming an anchor that might stabilize the ligand Met-AMP at this site. This loop primarily increased tRNA binding via Lys299 and Lys302. The region surrounding this loop also showed repulsive binding to tRNA via Glu298, Asp335, and Glu341. The attractive and repulsive forces between tRNA and the protein’s active site region led to the higher fluctuation of this CP domain and KMSKS protein region (shown in RMSF analysis) that might open the active site. This may be required to accommodate the tRNA’s CCA end (acceptor stem) for productive interaction with Met-AMP and eventually the product release.

During these events, tRNA remained strongly bound (electrostatic attractions) via its anticodon region to the anticodon domain of the protein (**Figure 5e**). tRNA anticodon interacted favorably with arginine (Arg361, Arg434 and Arg509) and lysine (Lys368, Lys423 and Lys435). This strong anticodon binding could facilitate the tRNA charging at the active site by maintaining the productive tRNA conformation.

Literature evidence shows that KMSKS loop flips upon amino acid activation and reaches the closed conformation after Met-AMP formation.^20^ It was speculated that the binding of tRNA to the anticodon domain facilitates the access of 3’ CCA to the active site by opening the KMSKS loop.^21^ A mutation in *E. coli* MetRS at R533 of the anticodon domain affects the aminoacylation rate.^22^ This residue was presumed to cause productive 3’ CCA binding to the KMSKS and CP domain.^22^ We observed the equivalent *Mtb* MetRS residue R509 interacts with tRNA. The mutagenesis in *E. coli* MetRS shows a main role of the CP domain in methionine activation and amino acid transfer to the 3’ CCA of tRNA.^23,24^ In *E. coli* MetRS, CP domain mobility possibly relates to these catalytic steps.^18,25,26^ This domain was assumed to force the tRNA’s acceptor arm to the active site.^21^ Therefore, our observations were well supported by the experimental data in literature.

### tRNA-ligand contacts were more for the elongator complex

Initiator tRNA and Met-AMP interacted for a shorter time while the elongator tRNA maintained hydrogen bond and salt-bridge contacts with the ligand for longer (**Figure S21**). The average hydrogen bonds and salt-bridges in the initiator were 0.21 and 0.14, and in the elongator were 0.86 and 0.44, respectively (**Figure 4d-e**). In addition, minor changes in π-π, π-cation, and π-anion interactions were observed (**Figure S21c-e**).

These results suggest that while maintaining a stable state in the protein’s active site, Met-AMP interacts with tRNA via favorable contacts. These attractive tRNA-ligand contacts worked simultaneously with the repulsive forces between tRNA and the protein’s active site region. These opposite forces, acting in parallel, influenced the ligand binding to tRNA and active site opening, possibly for the reaction and the release of products.

### Protein-ligand contacts and binding were higher for the elongator complex

The protein-Met-AMP complex exhibited more hydrogen bonds and salt-bridges in the elongator complex (2.22 and 0.88, respectively) than in the initiator (1.00 and 0.74) (**Figures S22** and **4f-g**). A minor variation in other interactions was observed (**Figure S22c-e**).

Importantly, stronger protein-ligand binding occurred in the elongator state (ΔG_bind_ = -27.95 kcal/mol) than in the initiator (-12.46 kcal/mol) (**Figure S23a and 4h**). In elongator complex, Tyr12, Asn14, His21, Glu24, Lys54, Phe292, Lys302, Ser 303 and Val304 mainly contributed to ligand binding (**Figure S23b-e**). In contrast, the initiator complex showed a few contributing residues, reflecting dissociation of the ligand (**Figure S23b-d**). We also analyzed the interactions of Mg^2+^, as detailed in **Figure S24**. Mg^2+^ formed ionic contacts mainly with tRNA.

These indicate that more intermolecular interactions in the elongator led to the stronger binding of the ligand, whereas the weaker binding in the initiator was attributed to fewer ligand interactions.

### Initiator versus elongator tRNA features

We analyzed the characteristic features of tRNA reported in the literature and checked their presence in our simulations. Two typical features of initiator tRNA include the absence of a hydrogen bond between the nucleotides C1 and U73, and its presence between A11 and U25.^27^ We found a similar tendency with an average bond of 0.02 and 1.76, respectively (**Figure S25a-b and S26a-b**).

Similarly, the elongator tRNA is known to form hydrogen bonds between the nucleotides G1 and U73.^28,29^ We found a bonding pattern of mainly two hydrogen bonds (**Figure S25c**), with an average of 1.07 (**Figure S26c**).

The absence of a hydrogen bond between C1 and U73 in the initiator tRNA might provide conformational freedom to charge the 3’ CCA at the protein’s active site on a shorter timescale. In contrast, the hydrogen bonds between the same positions G1 and U73 in elongator tRNA possibly restrict its conformational unwinding, which results in a longer time for tRNA charging. This validation was also strengthened by another experimental fact: a short lifetime occurred for the initiator complex and a long one for the elongator in *E. coli*.^18^

### Anticodon recognition for initiator and elongator tRNA

Methionine tRNA anticodon CAU (35-37 position) is conserved and forms a main recognition element for interacting with MetRS (anticodon domain: 358-519) during the aminoacylation reaction.^2,30^ Therefore, we analyzed the hydrogen bonds between CAU bases and protein (**Figure S27** and **S26d-f**). All three bases formed hydrogen bonds, with C35 and A36 showing more bonds in the initiator tRNA, while U37 exhibited more contacts in the elongator tRNA. The initiator and elongator protein formed average hydrogen bonds (of three MD runs) of 1.57 and 0.14 with C35, 0.59 and 0.41 with A36, and 0.25 and 0.56 with U37 bases, respectively.

These differences suggest that initiator tRNA recognition relies on C35 and A36 bases, whereas elongator tRNA prefers U37 along with A36 base.

### Protein anticodon recognition site interaction with tRNA

The anticodon domain of MetRS performs an essential function of tRNA anticodon binding, primarily involving three conserved residues, Asn357, Arg361, and Trp431 (**Figure S28**).^2,31,32^ The hydrogen bonding between these residues and tRNA was analyzed (**Figures S29** and **S26g-i**). The average hydrogen bonds of the initiator and elongator tRNA with Asn357 were 0.94 and 0.43, Arg361 were 0.35 and 0.66, and Trp431 were 0.04 and 0.08, respectively.

Thus, while the interactions of tRNA with the conserved residues were meaningful, prominent interactions occurred with Asn357 and Arg361. Asn357 interacted more with the initiator tRNA, whereas Arg361 displayed more contact with the elongator.

### Protein and ligand interaction with the CCA end of tRNA

The CCA terminal (75-77 position) of tRNA is conserved and plays a key functional role in the acylation of the correct amino acid.^33,34^ Its interactions with both protein and ligand (Met-AMP) revealed that the CCA region of elongator tRNA typically formed more hydrogen bonds than the initiator (**Figures S30-S31** and **S26j-o**). The protein formed average bonds of 0.07 and 1.36 with C75, 0.32 and 0.90 with C76, and 1.08 and 1.10 with A77, of initiator and elongator tRNA, respectively.

Interestingly, the ligand did not form any hydrogen bonds with CCA bases in the initiator state, possibly due to its release from the binding site. The ligand in the elongator complex showed 0.32, 0.09, and 0.07 average bonds with C75, C76, and C77 bases, respectively (**Figure S31** and **S26m-o**).

This highlights that the tRNA’s CCA terminal forms hydrogen bonds with both ligand and protein that might enable the reaction progression, while the repulsive electrostatic interactions of tRNA with the protein’s active site possibly help accommodate tRNA and product release by attaining an open conformation of this site. The overall plausible mechanism of tRNA recognition by *Mtb* MetRS is summarized in **Figure 6 (**details in **Supplementary sections S3**-**S4** and **Figures S32-S40**).

**Figure 6.**
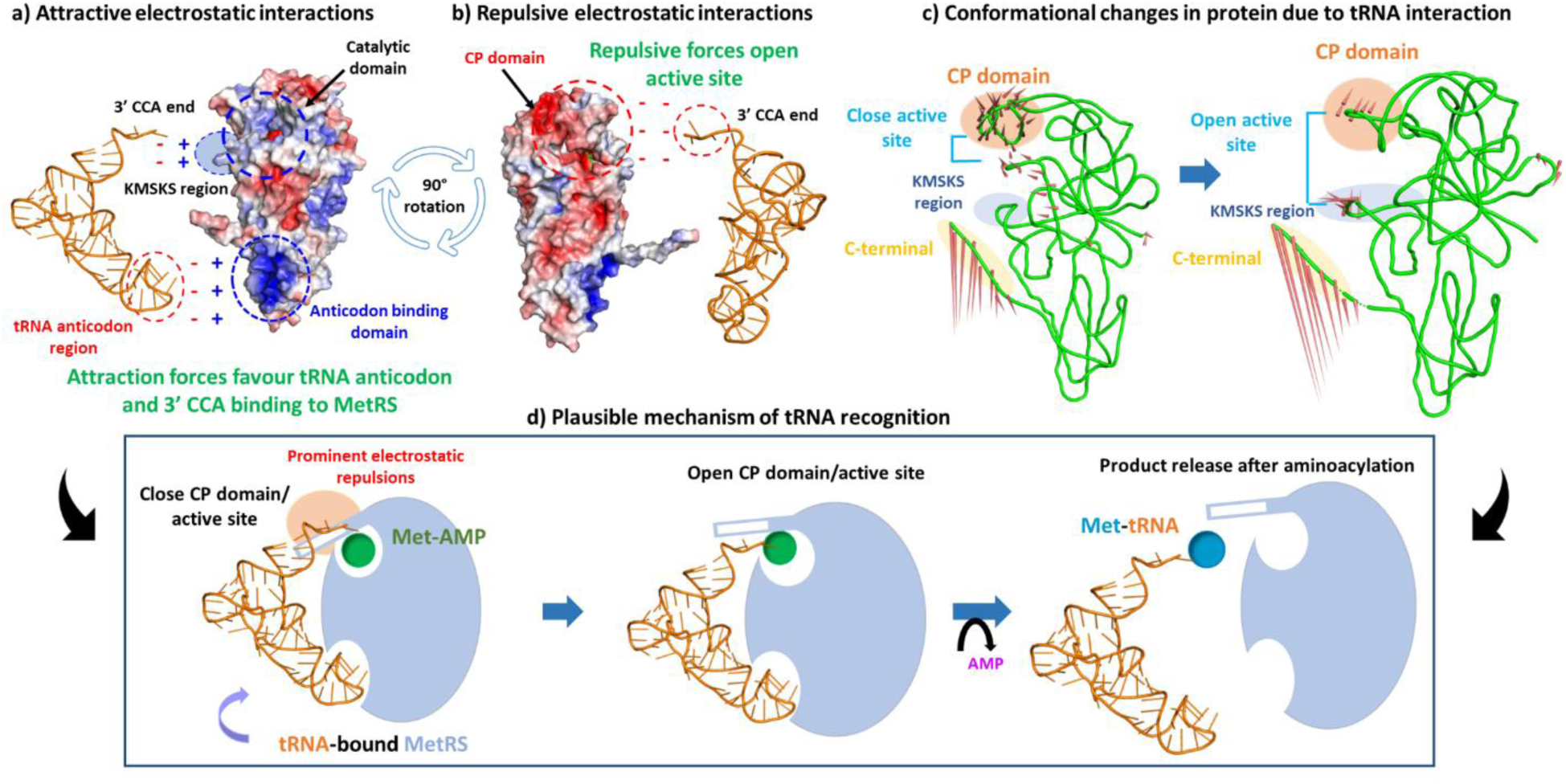
Plausible mechanism of tRNA recognition by *Mtb MetRS*. **(a)** Attractive interactions favor tRNA binding via its anticodon and acceptor arm to the MetRS anticodon domain and active site, respectively. **(b**) Negatively charged CP domain forms repulsive contacts with the acceptor arm, inducing active site opening. **(c)** MetRS achieves close to open conformation during tRNA interactions, an event favorable for the tRNA entry into the site and product release. **(d**) The CP knuckle opens, and tRNA enters the active site and interacts with Met-AMP, forming the product.

## CONCLUSIONS

The charging of tRNA by MetRS is a crucial step in protein synthesis. MetRS, which is involved in initiating and elongating protein synthesis, is an essential target for anti-tubercular drug design. The two tRNAs (each 77 bases in length) corresponding to the initiation and elongation processes shared a sequence identity of 67.9%. Due to their sequential and functional differences, they are expected to bind differentially to MetRS. Here, we found distinct conformational and binding features of tRNA-protein complexes (in the presence of Met-AMP) in the two states. Initiator tRNA showed higher conformational changes and transient binding to protein, while the elongator tRNA showed stable binding to MetRS. However, both tRNAs maintained the intra-tRNA interactions required for the typical cloverleaf structure. This concludes that the initiator tRNA charging with methionine could occur on a shorter timescale, while the elongator tRNA might require more charging time. This shorter timescale of the initiator tRNA could be linked to its necessity for initiating protein synthesis. tRNA interacts at two protein pockets: tRNA CCA terminal interacts with the protein’s active site (aminoacylation site), while tRNA anticodon CAU (35-37 position) binds to the protein’s anticodon domain. The tRNA charging at the two protein sites could be divided into three parts: (a) The electrostatic attractions between the tRNA and the protein’s catalytic domain could help tRNA charging with methionine, (b) the repulsive forces of tRNA with the protein’s connective peptide and both repulsive and attractive contacts with KMSKS region could open the active site to accommodate tRNA and cause the product release, (c) and the strong tRNA electrostatic binding via its anticodon to the protein’s anticodon domain could facilitate the reaction progression by maintaining the productive tRNA conformation. These events indicate a structural coupling between the two protein sites. Our study provides a mechanistic framework of how *Mtb* MetRS distinguishes and processes two sequentially divergent tRNAs and the plausible tRNA charging mechanism through conformational flexibility and electrostatic complementarity.

## METHODS

### Nucleotide sequence alignment of tRNAs

The three tRNA sequences, tRNA^fMet^, tRNA^Met^, and tRNA^Ile2^, were retrieved from RNAcentral^35^ (**Figure S1**). The length of each tRNA is 77 nucleotides. The pairwise local and global sequence alignments were performed using BLASTN^15^ and EMBOSS Needle.^16^

### tRNA-protein complex building

The tRNA-bound structure for *Mtb* MetRS was absent from the Protein Data Bank (PDB).^36^ However, a 3D structure of the initiator tRNA complexed with ribosome was available for *Mycobacterium smegmatis* (PDB structure: 8FR8), a close homolog of *Mtb*.^37^ Both shares a 100% sequence identity for the initiator tRNA. Therefore, we used the 3D coordinates of *M. smegmatis* initiator tRNA to build the tRNA of *Mtb* counterpart (tRNA^fMet^).

For elongator tRNA, no structure for *Mtb* MetRS or a homolog was available in PDB, so we modeled its structure. Its secondary structure was predicted using RNAstructure^38^ and 3D coordinates were modeled using RNAComposer (**Figure 1e**).^39^

For protein 3D coordinates, the structure of *Mtb* MetRS was retrieved from PDB with code 5XET^10^ (**Figure 1a**). This protein bound to intermediate (Met-AMP) and one magnesium ion at its active site was used. Missing residues of protein were modeled using CHARMM-GUI.^40^

To build the starting geometry of protein-tRNA complexes for both initiator and elongator states of *Mtb* MetRS, we used the MetRS-tRNA structure from *Aquifex aeolicus* in the intermediate state (containing Met-AMP, PDB code 2CT8) (**Figure S2**).^2^ The protein sequence identity between *Mtb* and *A. aeolicus* was 41.3%. The *A. aeolicus* tRNA (in PDB 2CT8) shared an identity of 72.5% with *Mtb* elongator tRNA and 66.7% with *Mtb* initiator tRNA (using EMBOSS Needle alignment). Therefore, we structurally aligned the protein and tRNA linked to *Mtb* MetRS with the *A. aeolicus* protein-tRNA complex to build the starting geometries.

### MD simulations of tRNA-protein complexes

The two modelled *Mtb* tRNA-MetRS complexes were prepared at pH 7 in a rectangular box using CHARMM-GUI.^40^ The total charge of each system was adjusted to zero, and a salt concentration of 0.15 M was maintained by adding the required Na^+^ and Cl^-^. The CHARMM36 force field was applied^41^ and MD simulations were performed using GROMACS^42^ at 300 K and 1 bar. The systems were subjected to two-step equilibrations (NPT and NVT) for 100 picoseconds each. We used C-rescale barostat and V-rescale thermostat.^43^ Three replica simulations of each system were performed using NPT for 1000 ns with a time-step of 2 femtoseconds.

### Statistical analysis and molecular interactions

Primary analysis was performed over the whole MD trajectory (1000 ns). This involved computing the protein backbone RMSD, R_g_, and SASA, as well as the hydrophilic and hydrophobic components of SASA. Similarly, RMSD, R_g_, and SASA were also computed for the tRNA and ligand (Met-AMP).

The secondary analysis was performed on the last 500 ns of the MD trajectory. FEL-PCA was performed on individual molecules, i.e., protein, tRNA, and ligand, and additionally on their complex form. We calculated RMSF and the number of inter- and intra-molecular interactions. The ratio of intra-molecular contact for protein and tRNA residues was determined as the total number of contacts divided by their mean value. The secondary structures of protein were analyzed using the VMD-SS plugin.^44^ The binding free energy (ΔG_bind_) was calculated for protein-tRNA and protein-ligand binding using gmx_MMPBSA^45^ at 10 ns intervals.

In addition, representative structures for the six simulated trajectories were extracted from the final 500 ns runs. Clustering was performed using GROMOS,^46^ with an RMSD cutoff of 0.4 nm. From the most populated cluster, the central structure was selected as representative.

## ASSOCIATED CONTENT

### Supporting Information

Detailed analysis of simulations is provided in the supporting file.

## AUTHOR INFORMATION

### Author Contributions

RM conceptualized and supervised the work. ST performed the work. RM and ST analyzed the work. RM and ST wrote and refined the manuscript.

### Notes

The authors declare no conflict of interest.

## ACKNOWLEDGEMENTS

RM acknowledges Science and Engineering Research Board, Government of India, for Start-up Research Grant (SRG) SRG/2022/000304 and IIT Bhilai for Research Initiation Grant (RIG), 2005900. ST acknowledges the Ministry of Education, Government of India, for the PMRF fellowship grant 5001800.

## ABBREVIATIONS USED

Mtb: Mycobacterium tuberculosis
MetRS: methionyl-tRNA synthetase
PDB: Protein Data Bank
MD: molecular dynamics
RMSD: root mean square deviation
R_g_: radius of gyration
SASA: solvent-accessible surface area
FEL-PCA: free energy landscape principal component analysis
EV: eigenvector
RMSF: root-mean-square-fluctuation
CP: connective peptide.

