## Supporting Information for "Initiator and Elongator tRNA Recognition Mechanism in *Mycobacterium tuberculosis* Methionyl-tRNA Synthetase"

*Rukmankesh Mehra: 0000-0001-6010-1514*

*Shivani Thakur: 0000-0002-7869-6648*

**Section S1. Free energy landscape principal component analysis of protein showed similar conformational landscapes**

Free energy landscape principal component analysis (FEL-PCA) was performed to investigate the dominant motions of the protein in the initiator and elongator tRNA complexes (**Figure S5**). The first two eigenvectors (EV1 and EV2) accounted for a significant portion of the total motion. The plots of eigenvalues versus eigenvectors and EV1 versus EV2 showed that both initiator and elongator complexes exhibited a similar behavior. The initiator complex displayed the average of three runs for EV1 and EV2 as 12.02 $nm^2$ and 5.92 $nm^2$, and the elongator complex showed 11.57 $nm^2$ and 5.36 $nm^2$, respectively. This indicates relatively similar conformational landscapes of protein in both systems. Additionally, the rapid decay of eigenvalues after the first few components in both systems suggests that a limited number of principal components capture the majority of essential motions.

**Section S2. Differential motion of the complete initiator versus elongator complexes**

We analyzed the FEL-PCA analysis of the complete initiator versus elongator complexes, i.e., collectively for protein, ligand, and tRNA (**Figure S17**). We observed a higher motion for the initiator complex. The average EV1 and EV2 values for three runs were 4604.98 and 2662 nm2 for the initiator complex, whereas the corresponding values were 148.56 and 62.54 nm2 for the elongator. These values indicate the higher motion of the initiator complex, which was consistent with the above observations of higher tRNA and ligand mobility in the initiator complex. This means that the major contribution to complex mobility was from tRNA and ligand molecules, while protein contributed to it to a smaller extent.

**Section S3. Electrostatic surface charge analysis of protein**

Electrostatic potential surface analysis of representative protein structures (**Figure S32-S33**) revealed key differences in electrostatic patterns across three major tRNA interaction regions: (i) the CP1 domain, (ii) the anticodon-binding domain (ABD), and (iii) the KMSKS loop. These differences are closely associated with the conformational and energetic adaptations necessary for tRNA binding and ligand coordination.

In the CP1 domain, the negatively charged phosphate backbone of the tRNA interacts with the acidic residues in the CP knuckle region, generating repulsive forces. These repulsions appear to trigger conformational rearrangements in the CP knuckle loop, eventually opening the pocket to accommodate the CCA end of the tRNA for productive interaction with the Met-AMP ligand. This structural reorganization is energetically favorable, as it reduces unfavorable electrostatic interactions and enables precise positioning of the tRNA's 3' end in the active site, facilitating efficient methionine charging.

In the ABD, a cluster of basic residues particularly Arg361 (conserved), Lys368, Arg373, Arg434, Lys435, and Arg444 forms a positively charged electrostatic surface. These residues, located within positions 357-373 and 430-445, enable direct and stable recognition of the anticodon loop through long-range electrostatic attractions, without inducing major conformational changes in the domain, as confirmed by the low RMSF values observed in this region (**Figure S8**). These interactions stabilize tRNA binding and contribute favorably to the overall binding free energy through persistent hydrogen bonding and electrostatic complementarity.

In the KMSKS loop region, interactions between the negatively charged tRNA backbone and positively charged residues (e.g., lysine's and arginine's) create an attractive electrostatic field that brings the loop closer to the tRNA (**Figure S33**). This movement is supported by increased flexibility of the KMSKS loop, as indicated by high RMSF values (**Figure S8**). This conformational adjustment partially opens the ligand-binding pocket, weakening ligand-protein interactions as reflected in the increased ligand flexibility and exposure observed in the initiator complex. This shift may enhance the possibility of ligand-tRNA interactions, thereby altering the structural dynamics and binding energetics within the active site.

In summary, these findings reinforce the pivotal role of electrostatic complementarity in governing tRNA recognition, ligand coordination, and dynamic rearrangements within *Mtb* MetRS complexes, contributing to the functional specificity and stability of initiator and elongator tRNA interactions.

**Section S4. Plausible mechanism of tRNA interaction and recognition**

The molecular recognition of tRNAs by *Mtb* MetRS involves a coordinated interplay of electrostatic complementarity and structural adaptation. Electrostatic surface analysis of representative structures revealed three primary tRNA-binding regions on the protein: the CP1 domain, ABD, and the KMSKS loop. The phosphate backbone of the tRNA, being negatively charged, is electrostatically attracted to positively charged lysine and arginine residues in the ABD and KMSKS regions. Importantly, positively charged residues such as Arg361, Lys368, Arg373, and Arg444 within the ABD stabilize tRNA binding by favoring long-range electrostatic interactions without requiring significant conformational rearrangements. In contrast, the interaction with the CP1 domain (acidic residues) triggers repulsion, prompting a shift from a closed to an open conformation in the CP knuckle loop (residues 127-154; **Figure S8**)[1]. tRNA binding in elongator complex induced secondary structural rearrangements in the CP domain of *Mtb* MetRS, including β-sheet-to-coil transition and loss of a $3_{10}$-helix. This conformational and secondary structure transitions suggest remodeling of active site which might be essential for accommodating the 3' CCA end of the tRNA in the catalytic pocket. These initial recognition events may contribute to specificity and secure the tRNA in a favorable orientation for subsequent steps of aminoacylation.

Upon binding, elongator and initiator tRNAs demonstrate distinct interaction dynamics. The elongator tRNA adopts a more stable conformation, consistently engaging both the CP domain and the anticodon-binding region, which is reflected in its strong binding affinity and lower structural fluctuations. In contrast, the initiator tRNA shows flexible binding modes, including back-face interaction via the TΨC loop - a feature reminiscent of class II aaRS recognition.[2] This divergence in binding behavior highlights the idiosyncratic recognition strategy of the initiator tRNA and emphasizes its specialized function in initiating the translation process.

Simulation data further suggest that tRNA binding directly instigates conformational changes in the catalytic domain that weaken ligand-protein interactions, particularly in the initiator tRNA-bound complex (**Figure S23 and S34**). In multiple simulations, initiator tRNA engagement led to repositioning of the CP1 loop, thereby opening the ligand-binding site and destabilizing the Met-AMP ligand. Interestingly, this destabilization was not uniform across all simulations. The tRNA deviated from the active site (green trajectory) in one trajectory, which stabilized ligand positioning. However, in two other simulations (blue and light brown), the 3' CCA end persistently interacted with the CP1 domain, reshaping the pocket and weakening ligand retention. This ligand displacement was unique to the initiator tRNA complex and did not occur in the elongator complex, where the ligand remained stably associated. These findings suggest that the structural flexibility and distinct base-pairing at the acceptor stem of the initiator tRNA, including the unpaired C1:U73 base pair, may facilitate an interaction with Met-AMP on a shorter timescale, enhancing aminoacylation efficiency.

The interaction of tRNA with KMSKS loop region of MetRS also modulates ligand stability within the catalytic pocket. Particularly in the initiator complex, electrostatic pulling of the KMSKS loop induced by phosphate-backbone attraction leads to partial opening of the ligand-binding site, thereby weakening ligand-protein interactions and increasing the likelihood of ligand-tRNA contact. In contrast, the elongator complex maintains tight ligand coordination due to structural rigidity and reduced loop mobility.

**Table S1.** Nucleotide sequence alignment of tRNA sequences of *Mtb.*

| **Sequence name 1** | **Sequence name 2** | **BLASTN local alignment** | | **EMBOSS global alignment** | |
|---|---|---|---|---|---|
| | | **% identity** | **Query coverage** | **% identity** | **% similarity** |
| $tRNA^{fMet}$ | $tRNA^{Met}$ | NA | NA | 67.9 | 67.9 |
| $tRNA^{fMet}$ | $tRNA^{Ile2}$ | NA | NA | 64.3 | 64.3 |
| $tRNA^{Met}$ | $tRNA^{Ile2}$ | 81.97 | 79 | 77.5 | 77.5 |

*NA – No significant similarity found.

### a) Sequence of tRNAs found in *Mtb*

**tRNA$^{fMet}$ (encoded by *metU* gene) – 77 nucleotides**

CGCGGGGUGGAGCAGCUCGGUAGCUCGCUGGGCUCAUAACCCAGAGGUCGCAGGUUCGAAUCCUGUCCCCGCUACCA

**tRNA$^{Met}$ (encoded by *metT* gene) - 77 nucleotides**

GGCGAUGUAGCUCAGUCGGUUAGAGCGAACGACUCAUAAUCGUUAGGUCGCCGGUUCGAGUCCGGCCAUCGCUACCA

**tRNA$^{Ile2}$ (encoded by *metV* gene) - 77 nucleotides**

GGGGCGGUAGCUCAGUUGGUUAGAGCCGCGGACUCAUAAUCCGUUGGUCGCGGGUUCGAGCCCCGCCCGCCCCACCA

**Note:** The tRNA$^{fMet}$ contains the CCA end, but in tRNA$^{Met}$ and tRNA$^{Ile2}$, the missing CCA end is modified and added by CCA-adding enzyme during post-transcriptional modifications.

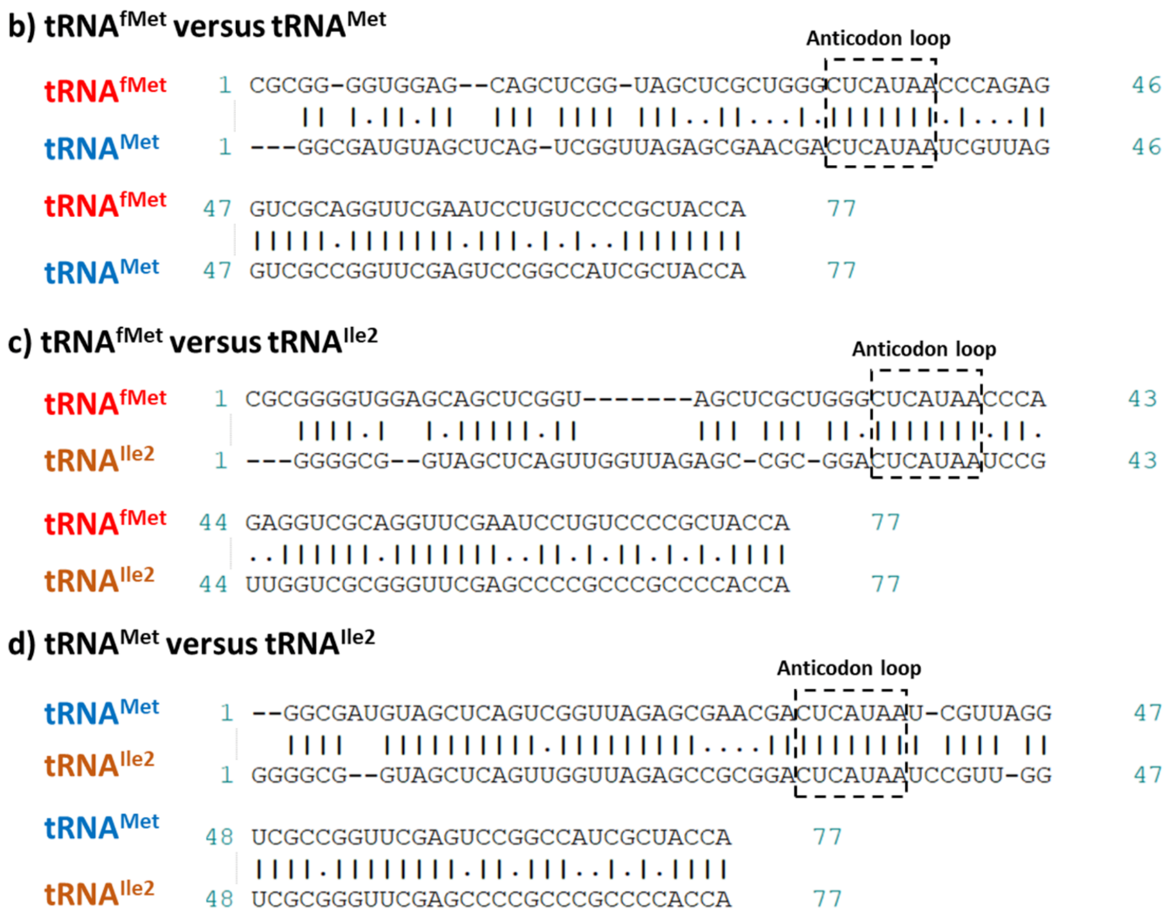


**Figure S1. Nucleotide sequence alignment of tRNAs. (a)** Sequences of tRNAs (tRNA$^{fMet}$, tRNA$^{Met}$, tRNA$^{Ile2}$) containing CAU anticodon found in *Mtb* Sequence alignment of **(b)** tRNA$^{fMet}$ versus tRNA$^{Met}$, **(c)** tRNA$^{fMet}$ versus tRNA$^{Ile2}$ and **(d)** tRNA$^{Met}$ versus tRNA$^{Ile2}$ highlighting conserved anticodon loop region.

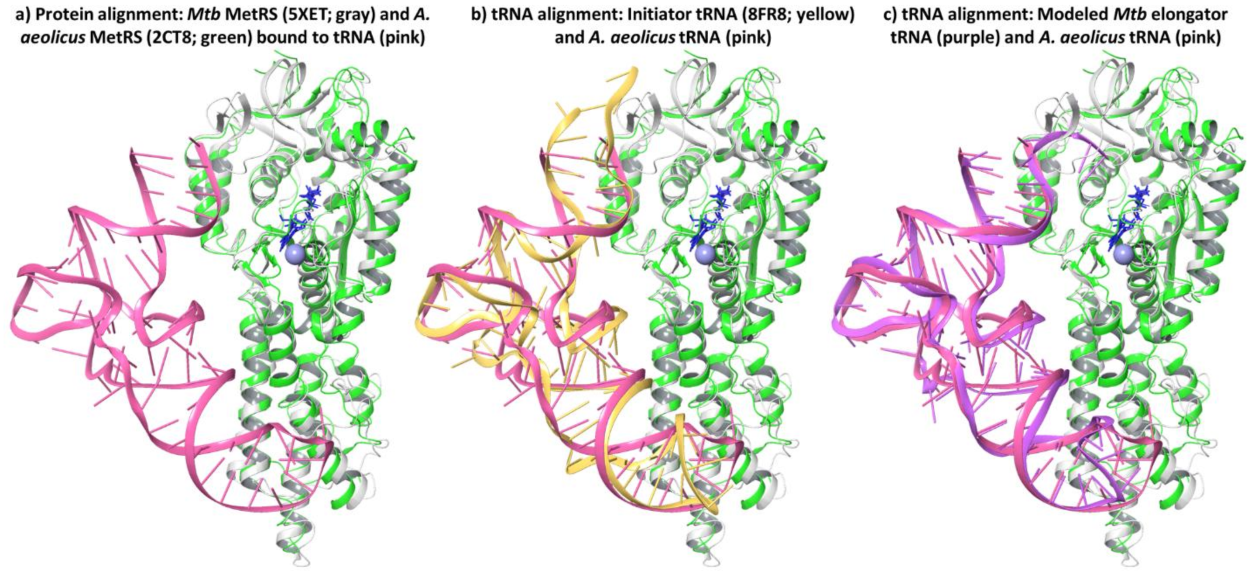


**Figure S2. Steps involved in tRNA-complex building. (a)** In first step, we aligned MetRS protein of *Mtb* (5XET; gray) with *A. aeolicus* (2CT8; green) bound to tRNA (pink). RMSD of MetRS protein (backbone) between *Mtb* (5XET) and *A. aeolicus* (2CT8) structures was low (i.e., 1.65 Å), suggesting high structural similarity between the two. Next step is alignment of tRNAs where we aligned **(b)** *M. smegmatis* initiator tRNA (8FR8; yellow) and *A. aeolicus* tRNA (2CT8; pink) to build initiator-tRNA complex and **(c)** modeled *Mtb* elongator tRNA (purple) with *A. aeolicus* tRNA (2CT8; pink) to build elongator-tRNA complex.

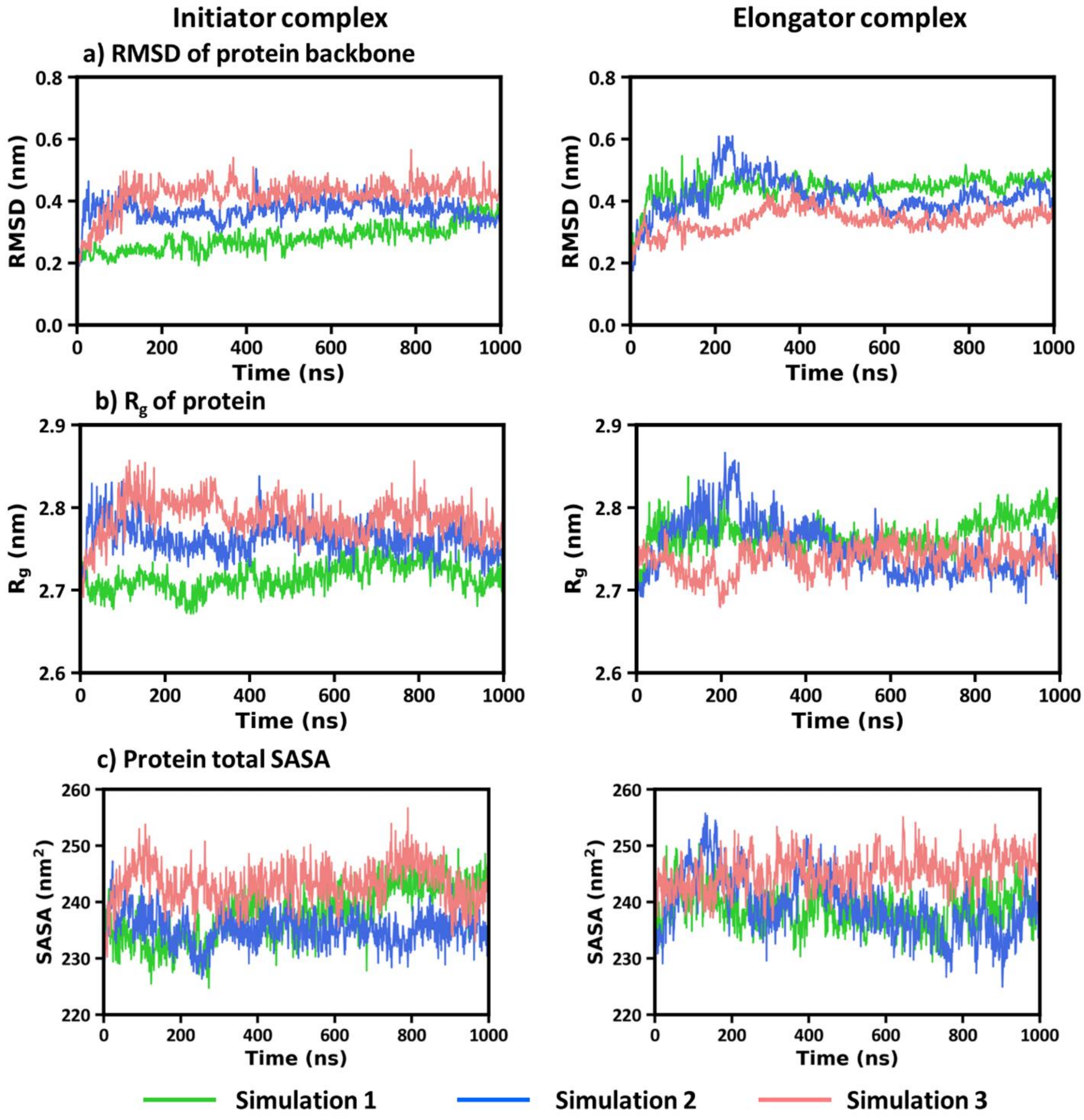


**Figure S3. The simulated properties of MetRS protein. (a)** The RMSD, **(b)** $R_g$, **(c)** total SASA of protein over three simulations (green, blue, peach). The representative colors for three simulations used in this figure are consistently followed in all subsequent plots.

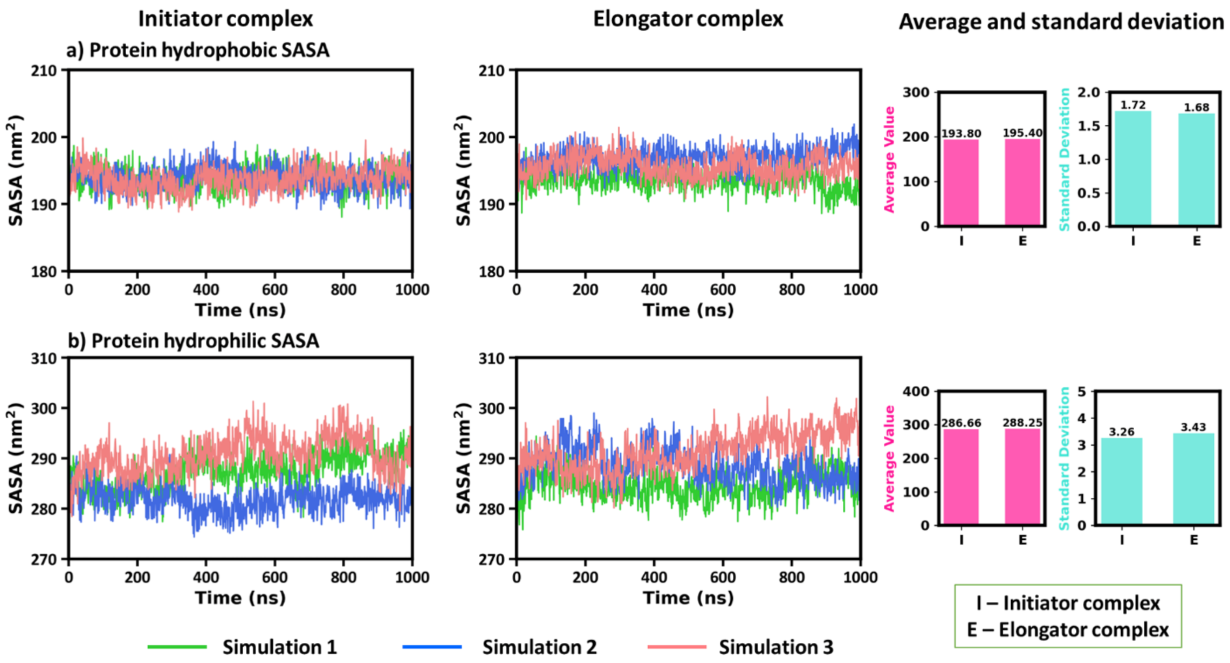


**Figure S4. Hydrophobic and hydrophilic solvent-accessible surface area (SASA) analysis of *Mtb* MetRS protein. (a)** Hydrophobic SASA and **(b)** hydrophilic SASA of protein over time for three simulations. Bar plots show the average and standard deviation of hydrophobic and hydrophilic SASA across the three simulations.

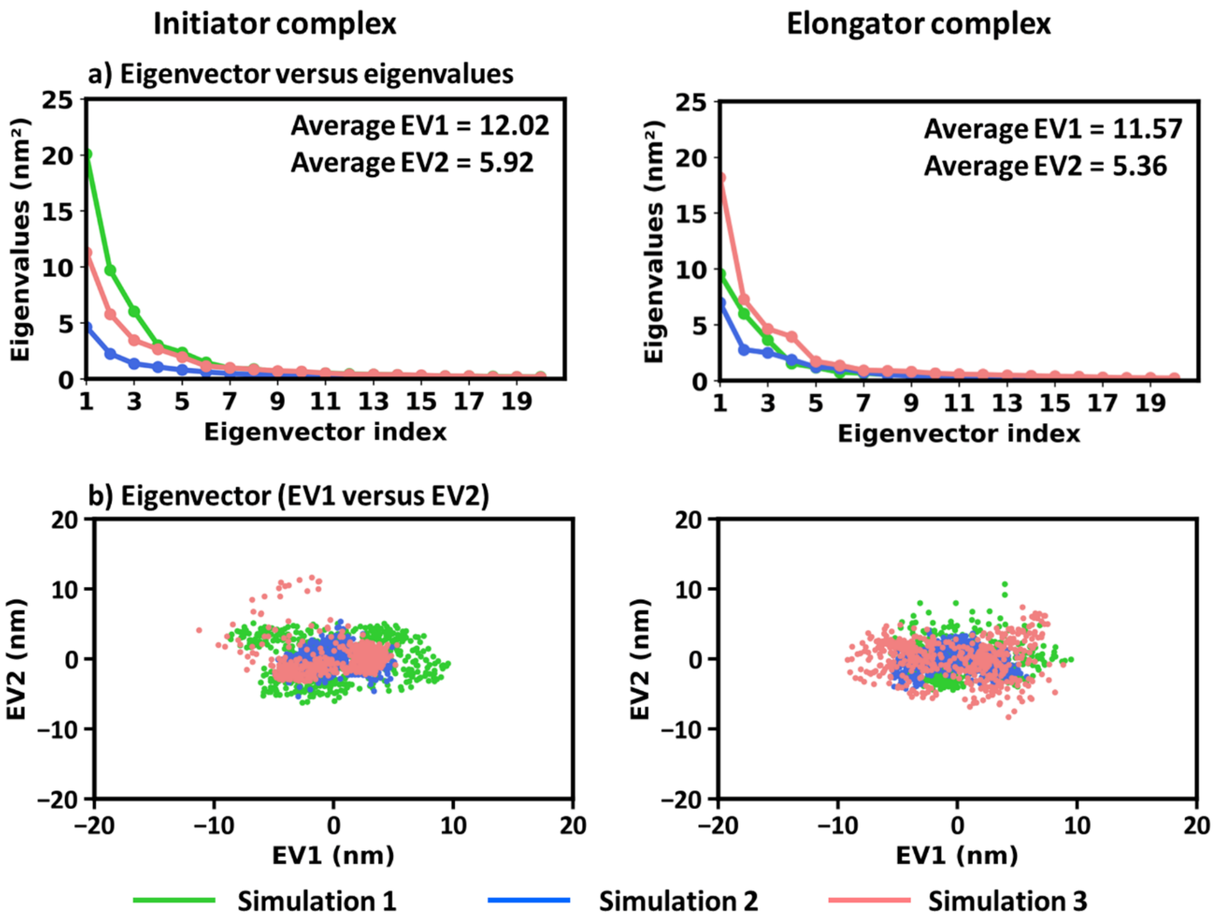


**Figure S5. Free energy landscape principal component analysis (FEL-PCA) of protein. (a)** First 20 eigenvectors (EV) with their eigenvalues and **(b)** EV1 versus EV2 projection plot for protein in initiator and elongator complex.

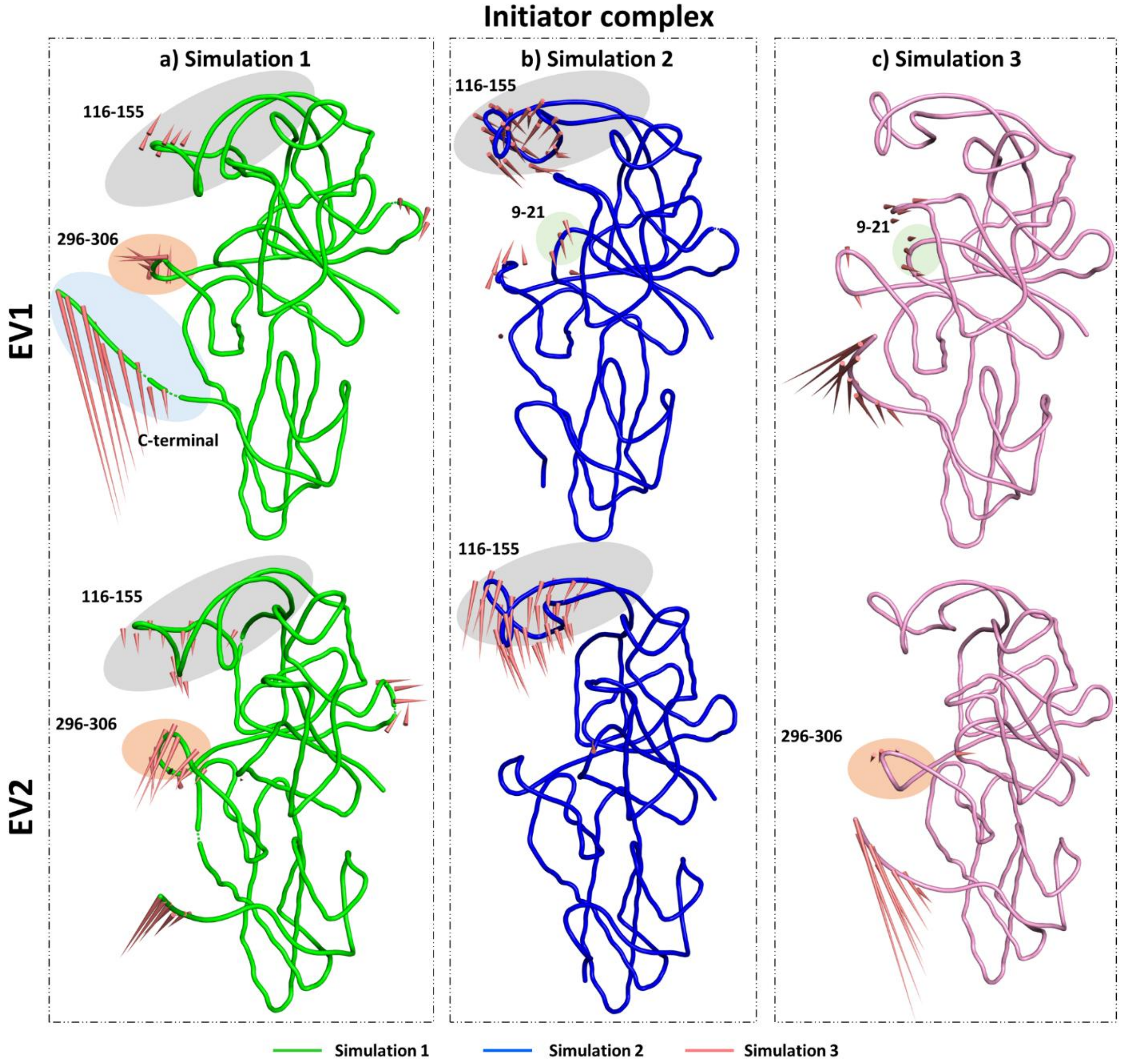


**Figure S6. Mobility of protein in initiator complex. (a) – (c)** Porcupine plots for first principal component (EV1) and second principal component (EV2) obtained from three simulations (represented in green, blue, peach color) of protein. The porcupine cones attached to average position of each backbone atom points towards direction of motion described by EV1 and EV2.

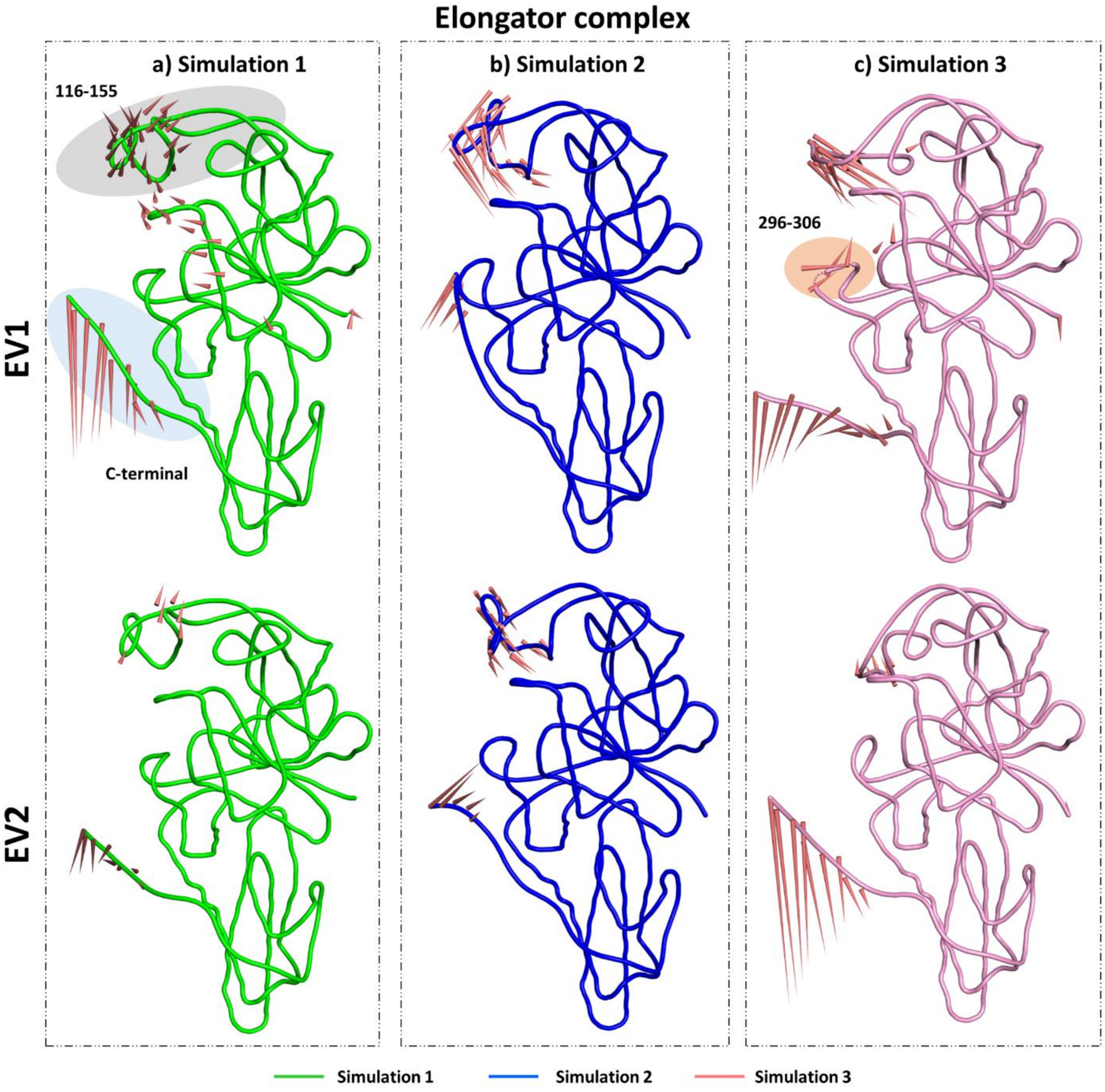


**Figure S7. Mobility of protein in elongator complex. (a) – (c)** Porcupine plots for first principal component (EV1) and second principal component (EV2) obtained from three simulations (represented in green, blue, peach color) of protein. The porcupine cones attached to average position of each backbone atom points towards direction of motion described by EV1 and EV2.

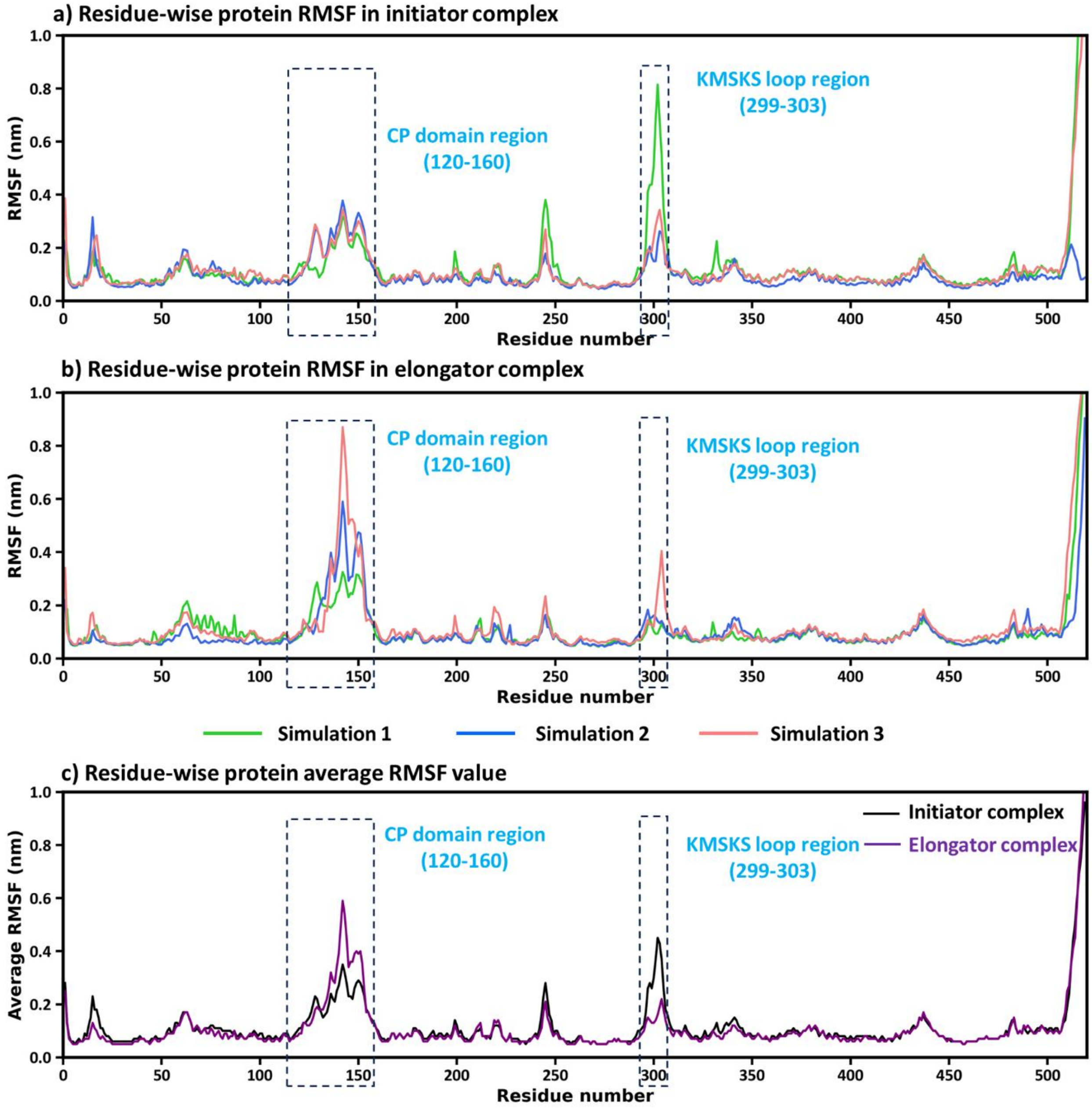


**Figure S8. Residue-wise RMSF of MetRS protein in (a)** initiator and **(b)** elongator tRNA-bound complexes across three simulations. **(c)** Average RMSF comparison highlights flexible regions, notably the CP domain (residues 120-160) and the KMSKS loop (299-303), showing differential mobility between complexes.

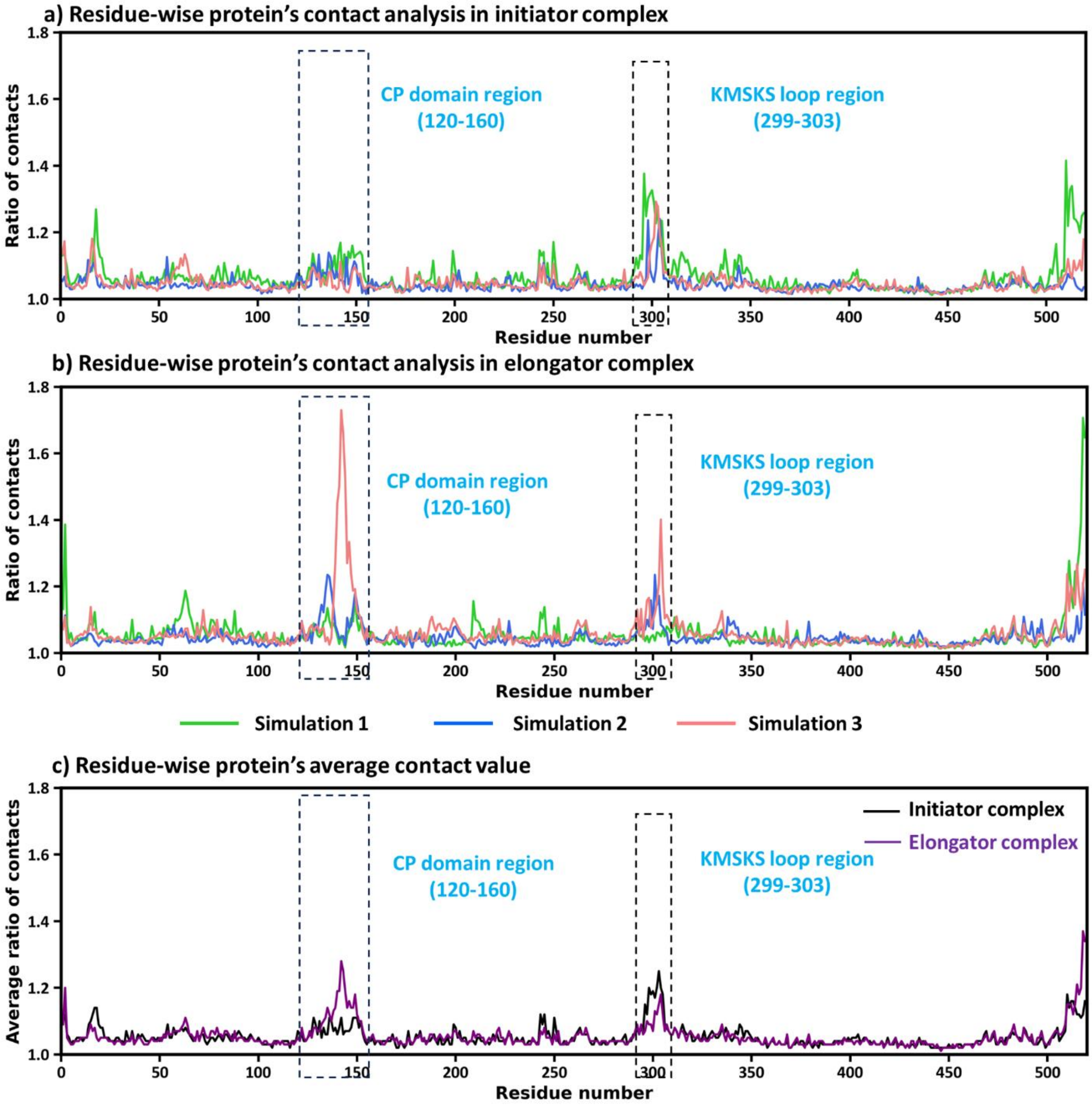


**Figure S9. Residue-wise analysis of contacts in MetRS upon tRNA binding for (a)** initiator and **(b)** elongator complexes over three simulations. Here "contacts" means count of number of different atomic contacts formed by each protein residue (x-axis) with atoms of other protein residues during simulations. "Ratio of contacts" is the total number of contacts divided by their mean value. **(c)** Average contact ratio comparison reveals enhanced contact formation in the CP domain (residues 120-160) and KMSKS loop (299-303).

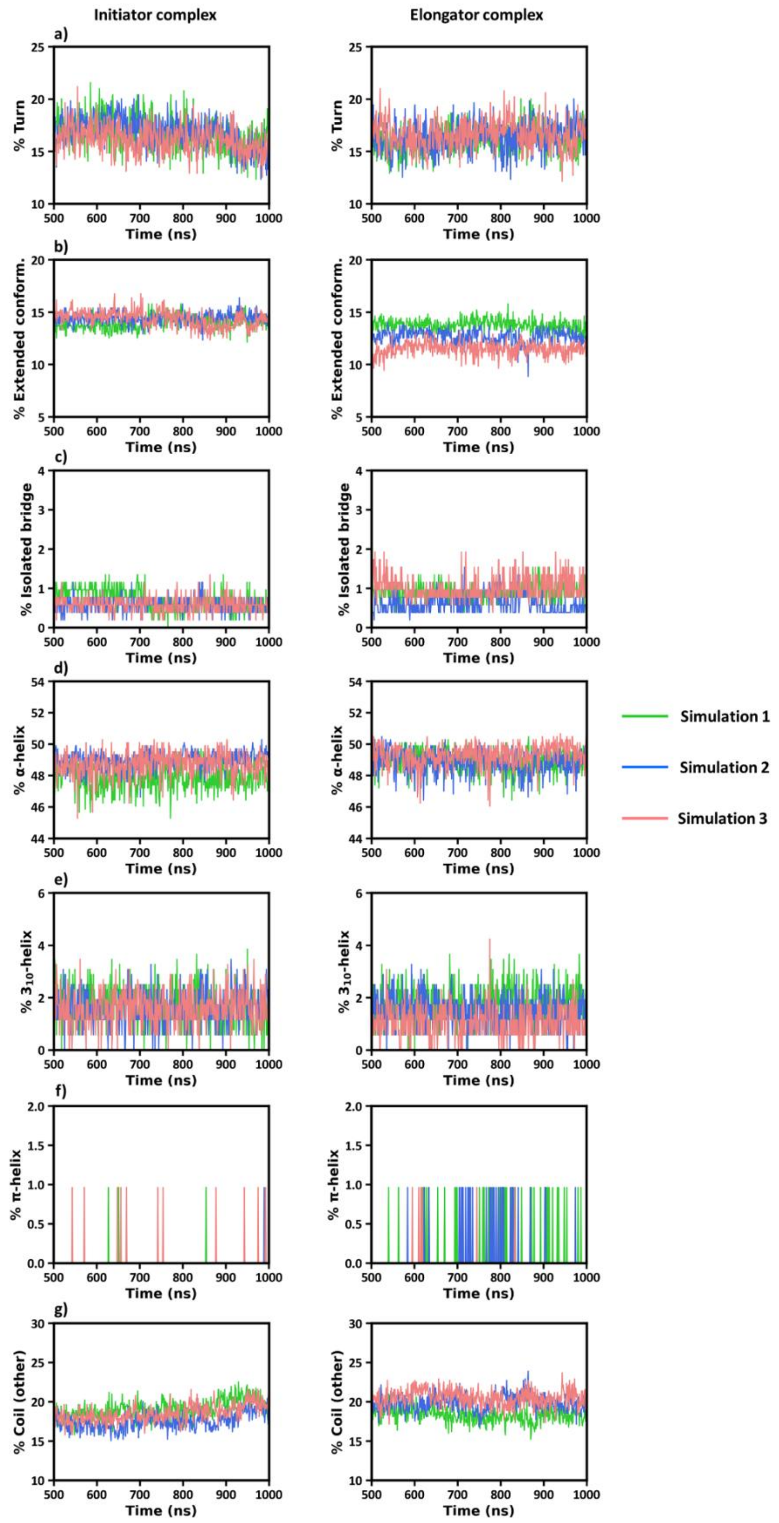


**Figure S10. Protein secondary structure elements. (a)** % Turn, **(b)** % Extended conformation or β-sheet, **(c)** % isolated bridge, **(d)** % α-helix, **(e)** % $3_{10}$-helix, **(f)** % π-helix and **(g)** % coil (other) in initiator (left) and elongator (right) complexes over 1 μs simulations.

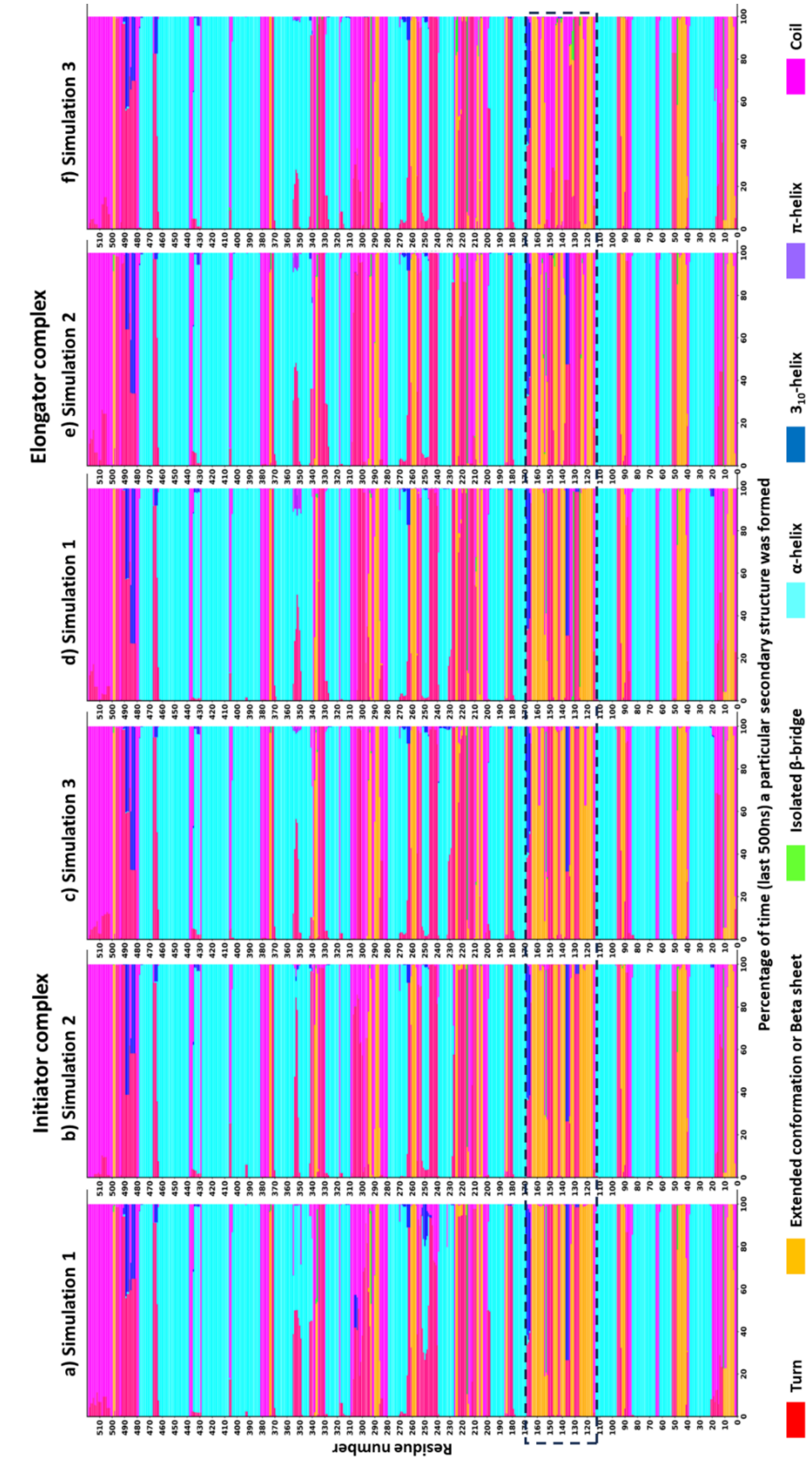


**Figure S11. Residue-wise secondary structure assignment of MetRS**. X-axis shows percentage of secondary structure formed by each residue of protein (Y-axis) across three simulations each for **(a-c)** initiator and **(d-f)** elongator complexes.

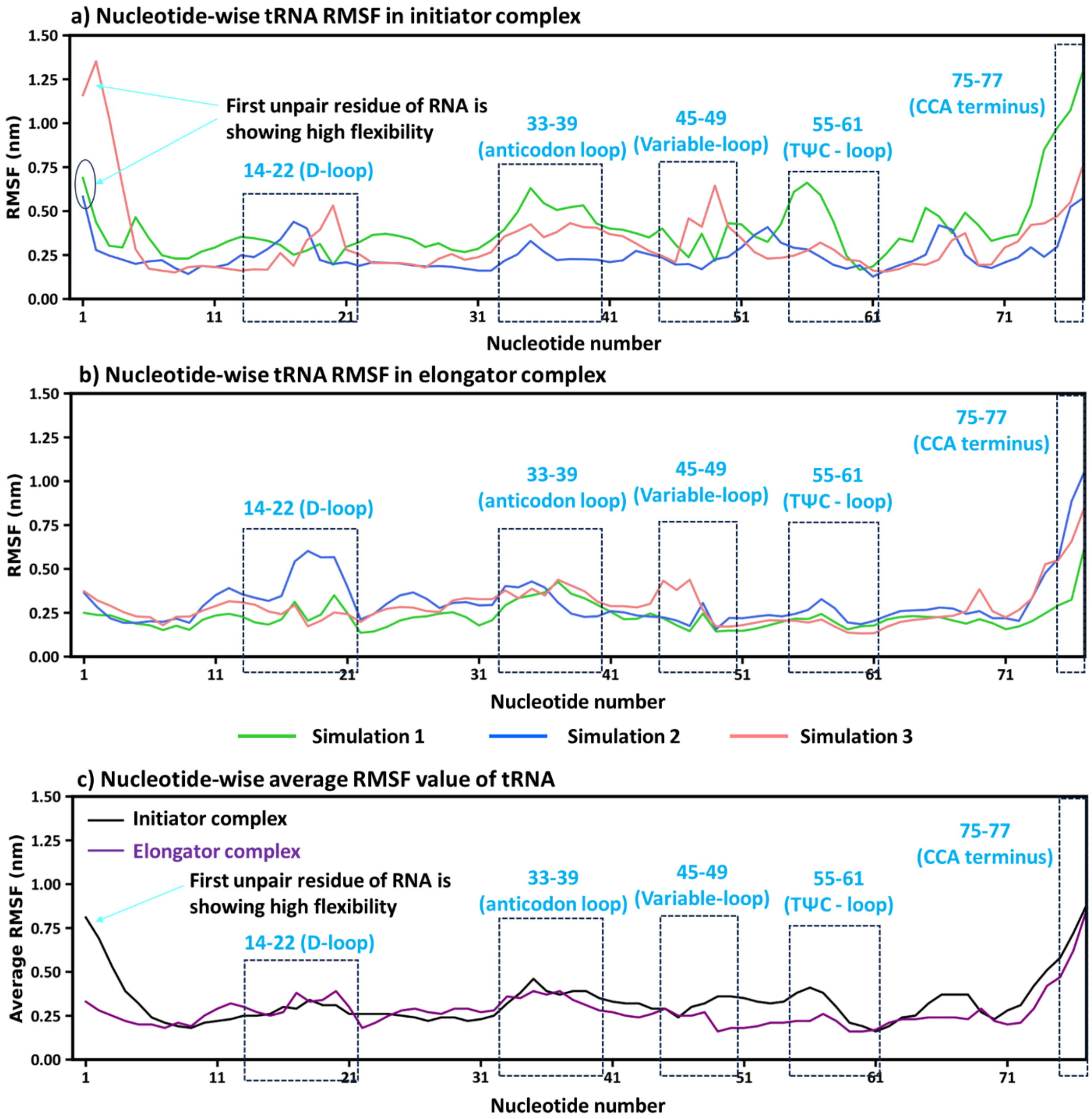


**Figure S12. Nucleotide-wise RMSF analysis of tRNA in (a)** initiator complex and **(b)** elongator complex. **(c)** Comparison of average RMSF values to show flexibility of nucleotides at the 5' end, D-loop (14-22), anticodon loop (33-39), variable loop (45-49), TΨC loop (55-61) and CCA terminus.

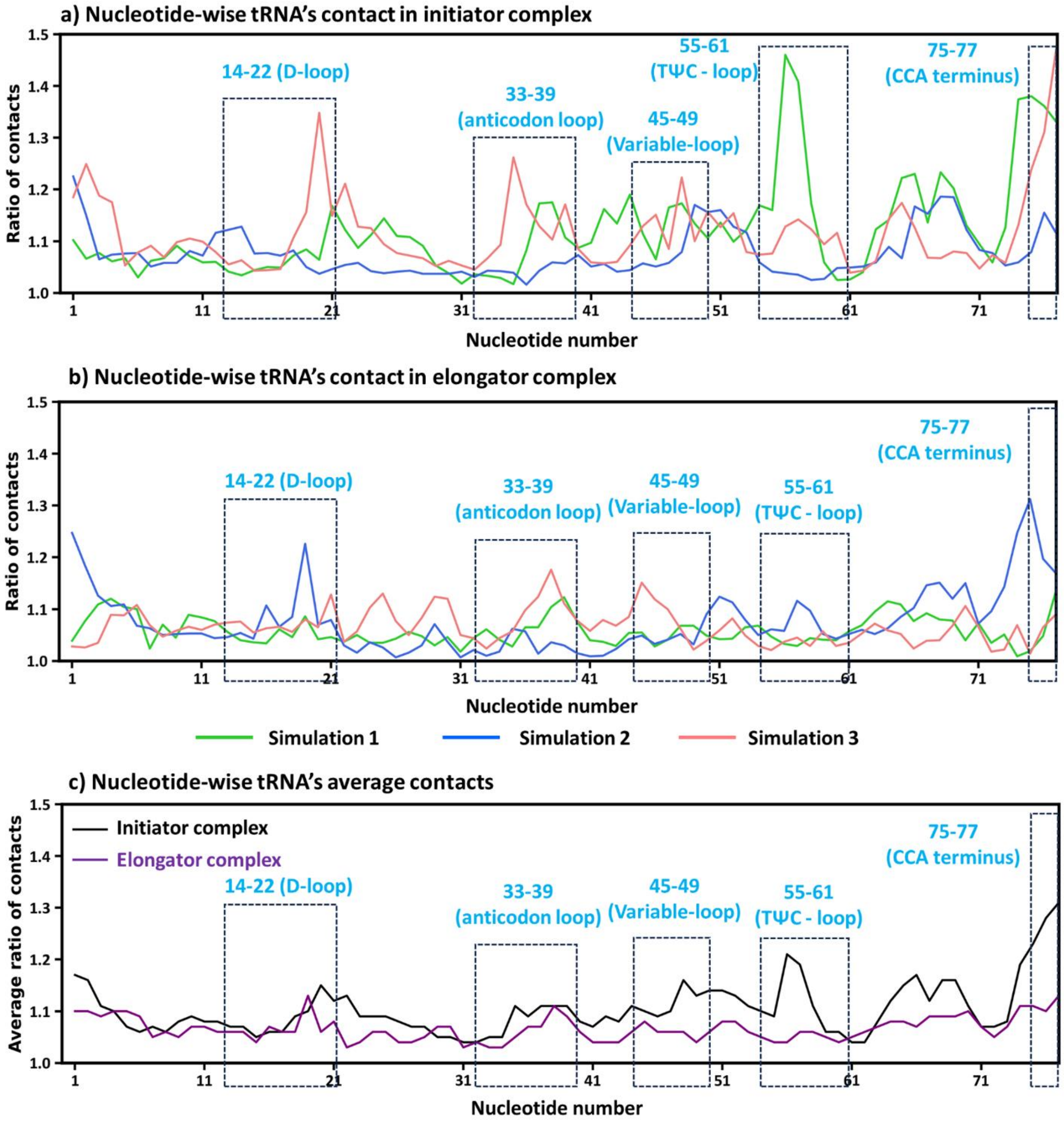


**Figure S13. Nucleotide-wise contact analysis of tRNA in (a)** initiator complex and **(b)** elongator complex. **(c)** Average contact ratio comparison reveals higher interaction dynamics in the initiator complex, especially at functionally important tRNA motifs. Here "contacts" means count of the number of different atomic contacts formed by each tRNA nucleotide (x-axis) with atoms of other nucleotides during simulations. "Ratio of contacts" is the total number of contacts divided by their mean value.

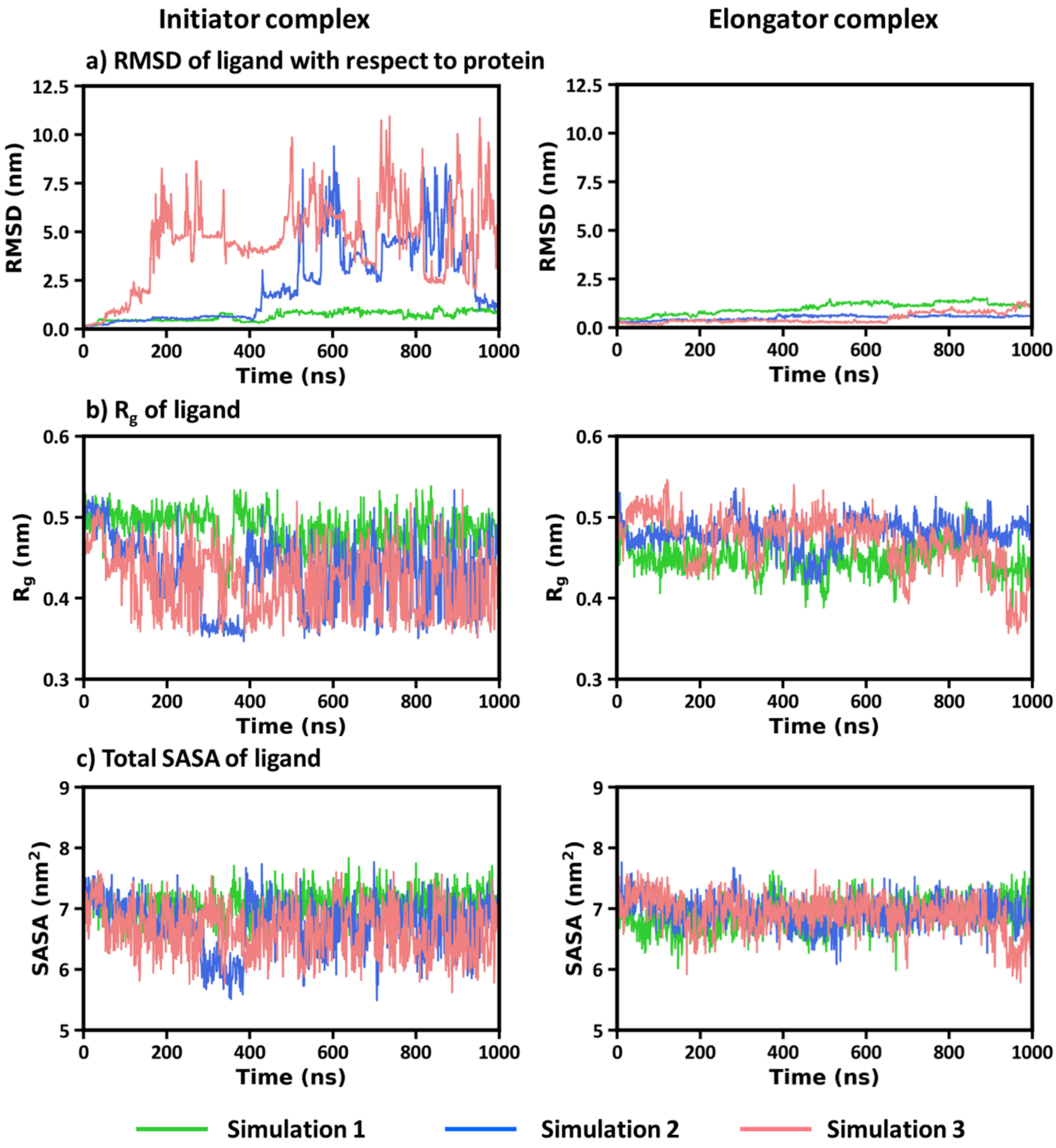


**Figure S14. Structural properties of the ligand (Met-AMP) in the initiator and elongator complexes. (a)** RMSD, **(b)** $R_g$, and **(c)** SASA of ligand.

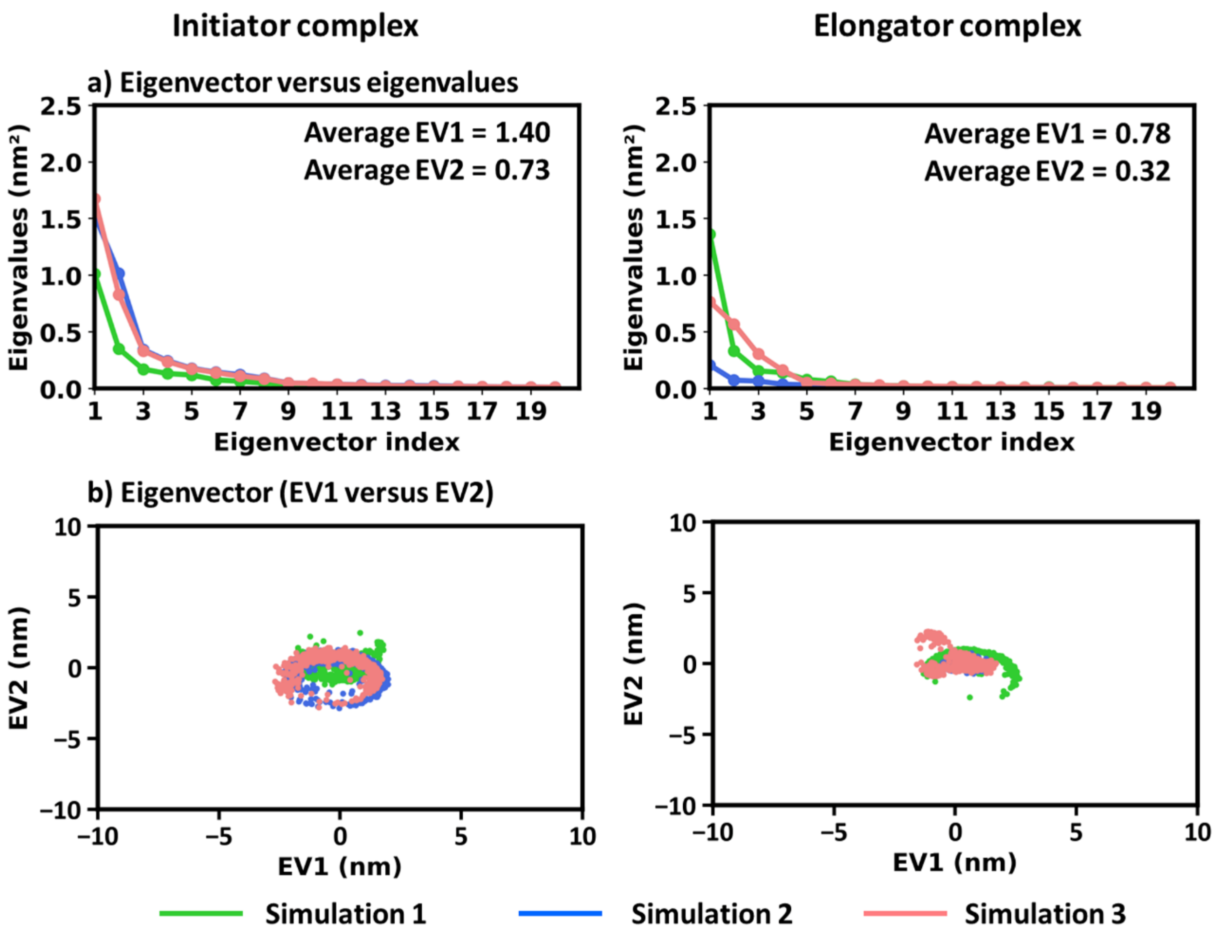


**Figure S15. PCA of the ligand (Met-AMP) in initiator and elongator complexes**. (a) Eigenvector versus eigenvalue plots. **(b)** 2D projection of eigenvectors (EV1 versus EV2).

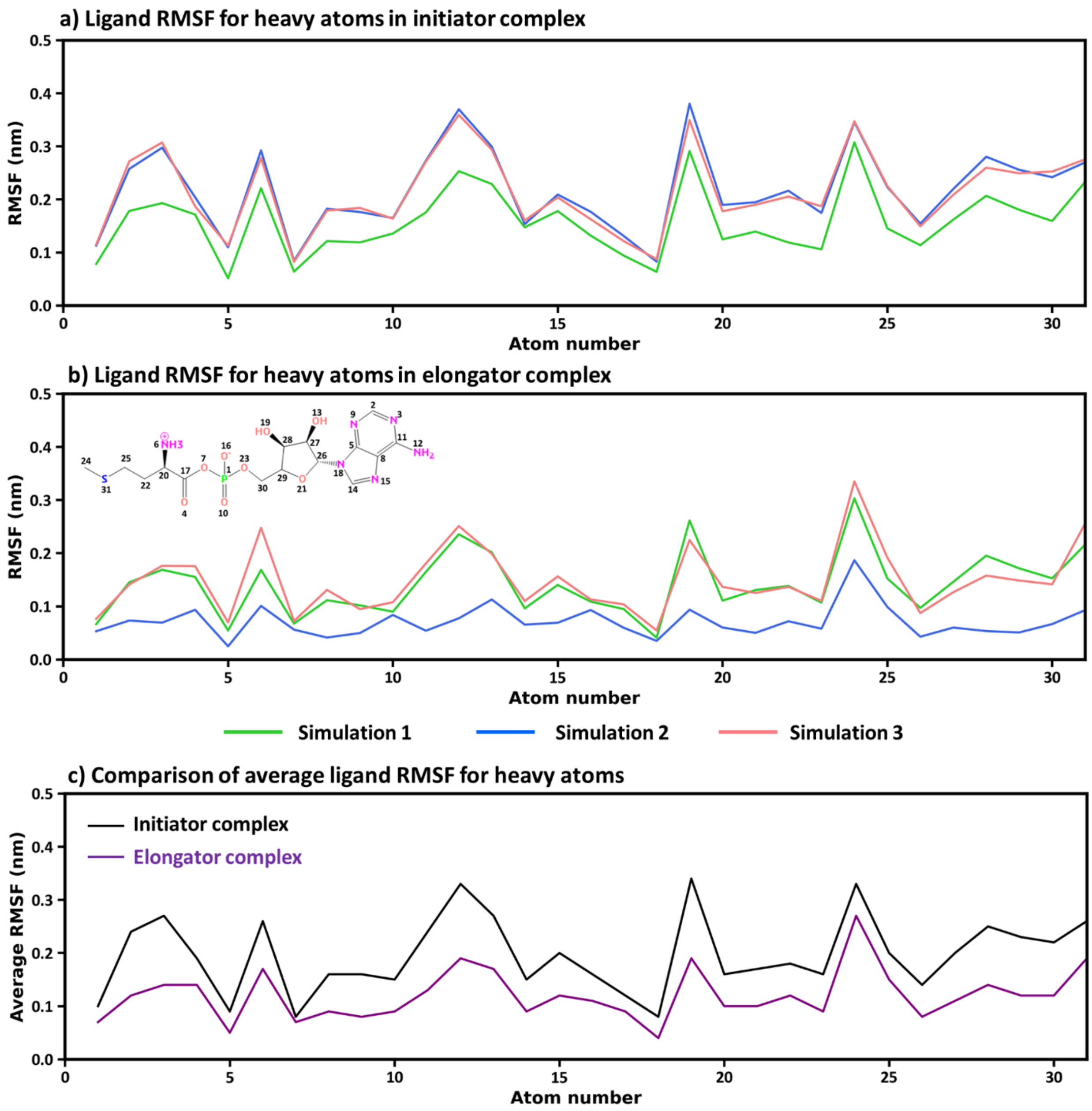


**Figure S16. RMSF analysis of heavy atoms in the ligand molecule. (a)** and **(b)** depict the heavy-atom RMSF of ligand in the initiator and elongator complexes, respectively. **(c)** Comparison of average RMSF values confirms reduced ligand mobility in the elongator complex.

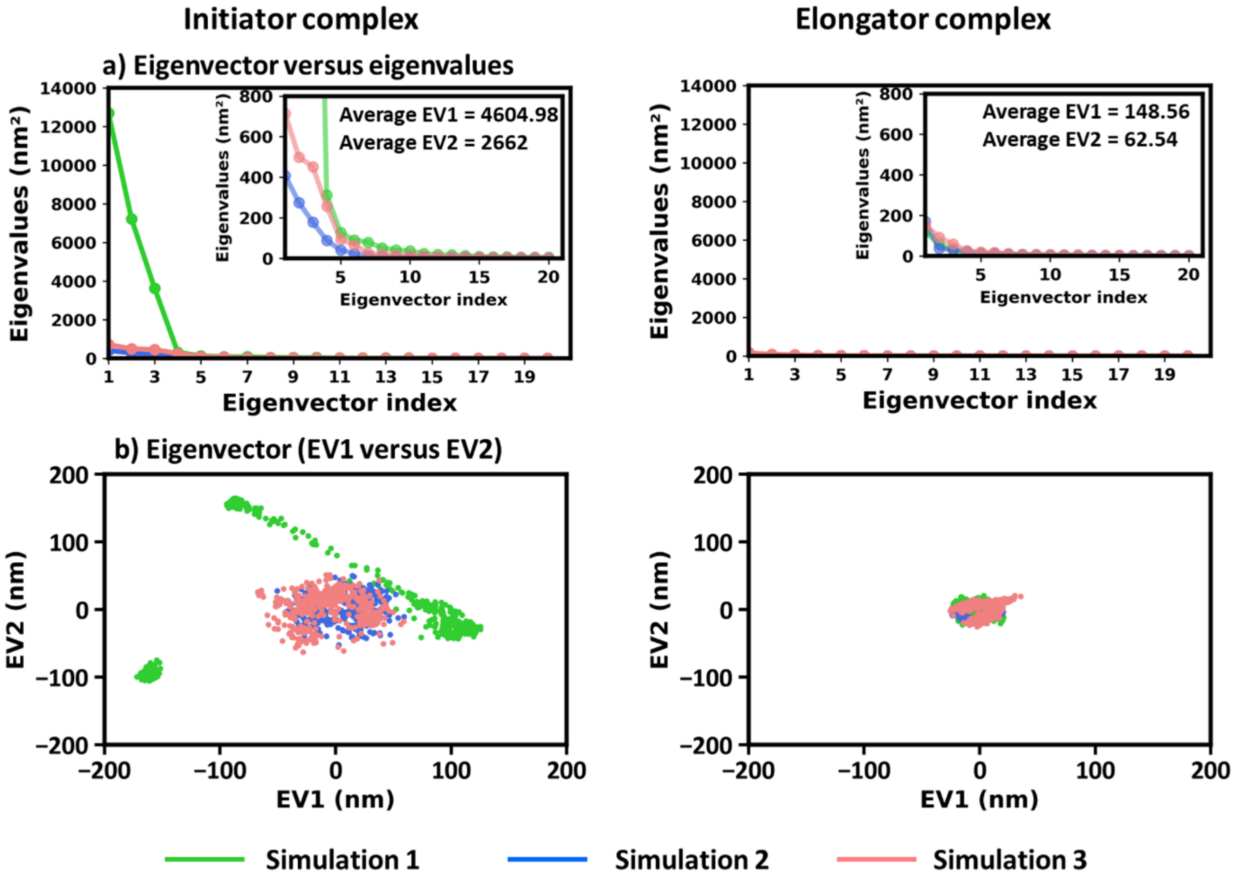


**Figure S17. PCA of complex (containing protein, ligand and tRNA) in initiator and elongator complexes. (a)** Eigenvector versus eigenvalue plots. **(b)** 2D projection of eigenvectors (EV1 versus EV2).

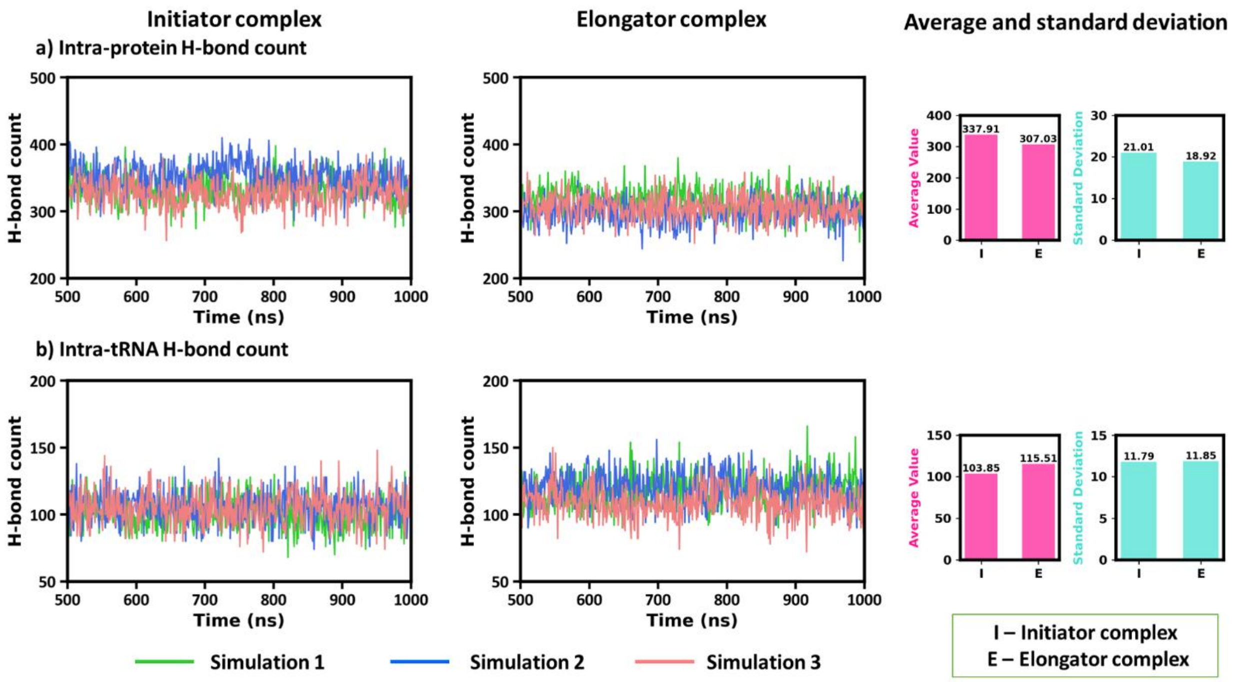


**Figure S18. Intramolecular hydrogen bonding of protein and tRNA.** Figure shows the H-bond count over time within **(a)** protein and **(b)** tRNA respectively, in initiator and elongator complexes. Average and standard deviation plots reveal higher intra-protein H-bonds in the initiator, while intra-tRNA H-bonding is slightly greater in the elongator complex.

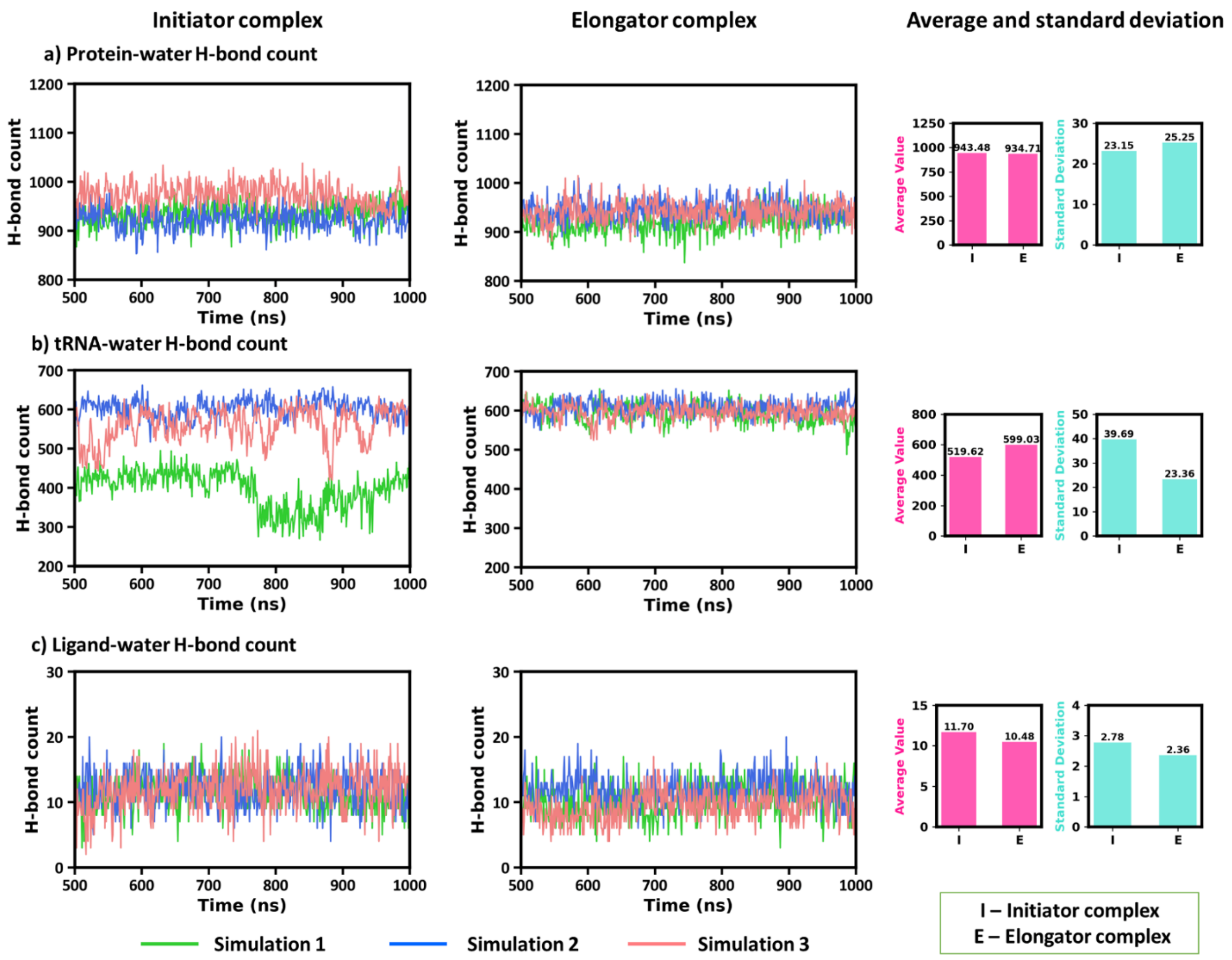


**Figure S19. Hydrogen bonds with water in initiator (I) and elongator (E) complexes.** H-bond counts for **(a)** protein-water, **(b)** tRNA-water, and **(c)** ligand-water interactions. Bar plots summarizing the average and standard deviation of H-bond counts.

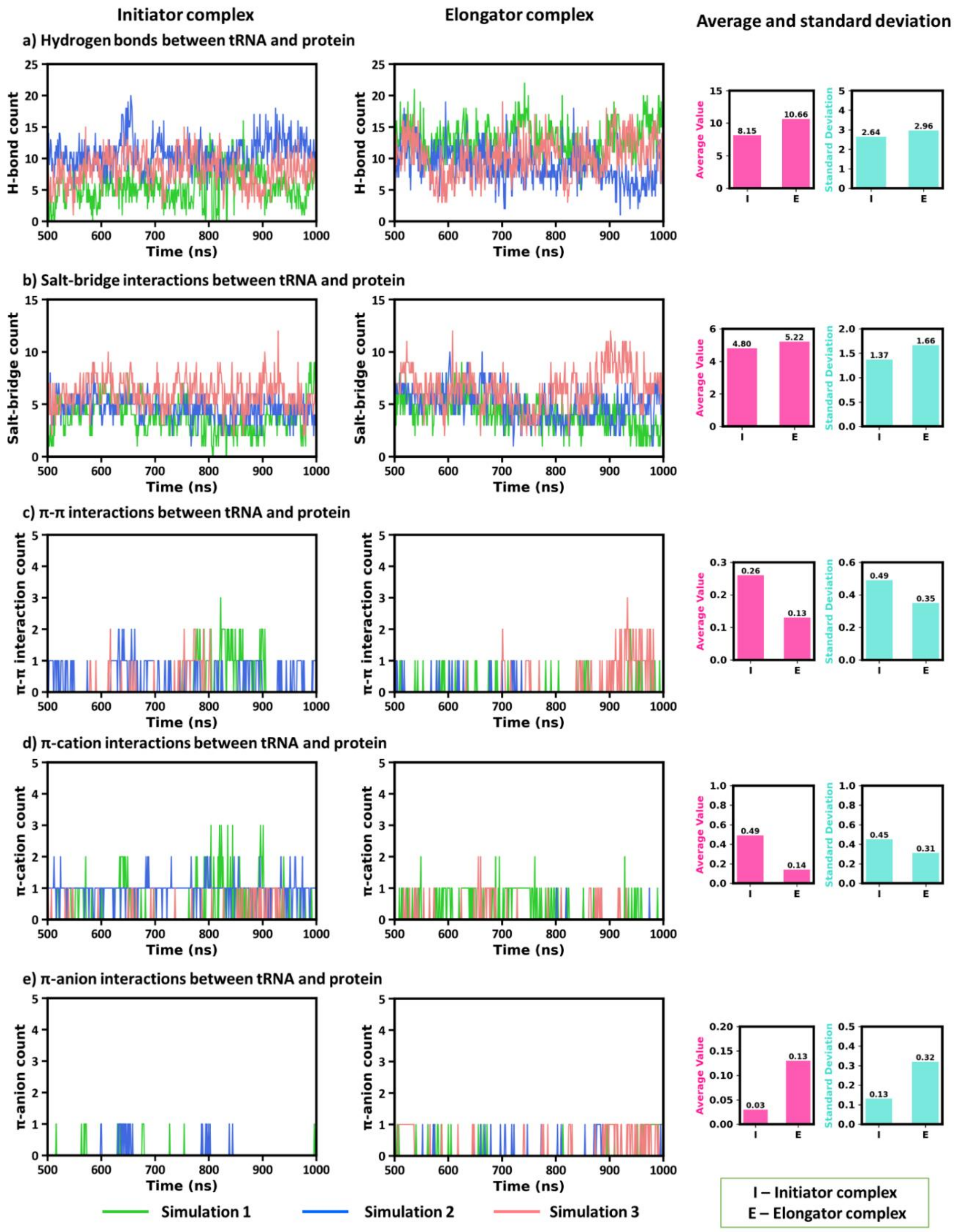


**Figure S20. Intermolecular interactions between tRNA and protein in initiator and elongator complexes. (a)** Hydrogen bonds. **(b)** Salt-bridge interactions. **(c-e)** Occurrence of π-π, π-cation, and π-anion interactions over time.

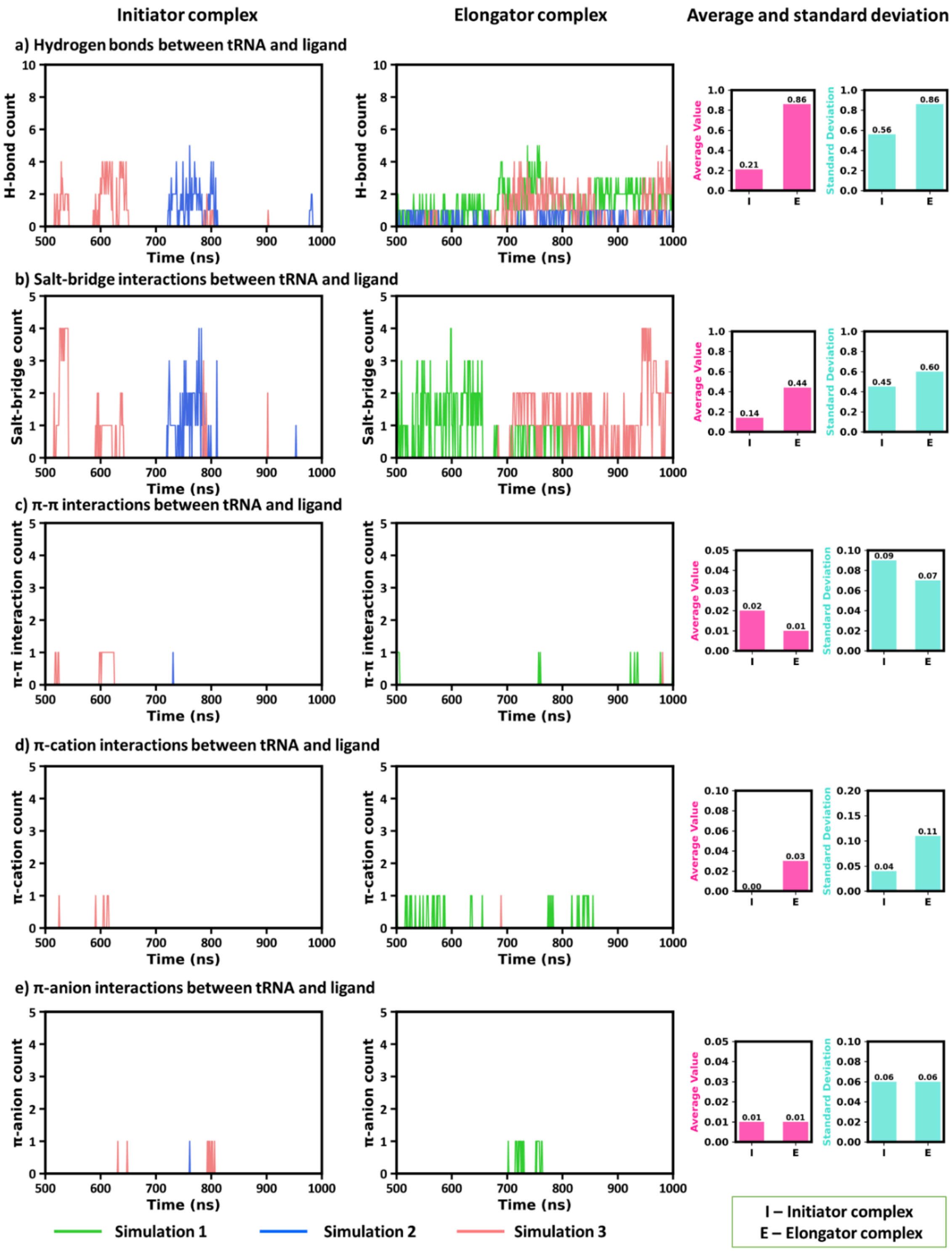


**Figure S21. Intermolecular interactions between tRNA and ligand in initiator (I) and elongator (E) complexes. (a)** Hydrogen bonds. **(b)** Salt-bridge interactions. **(c-e)** Occurrence of π-π, π-cation, and π-anion interactions over time.

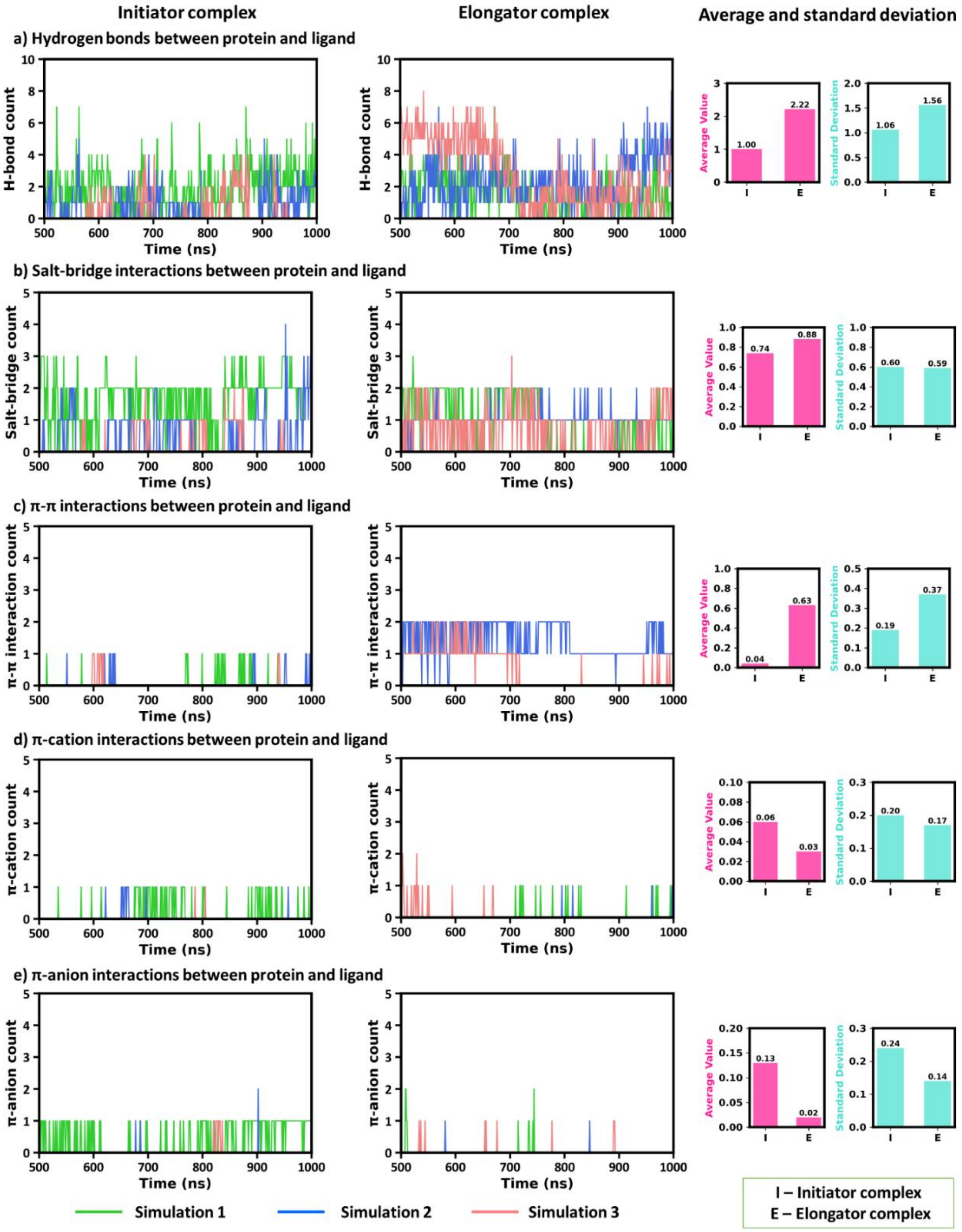


**Figure S22. Intermolecular interactions between protein and ligand in initiator (I) and elongator (E) complexes. (a)** Hydrogen bonds. **(b)** Salt-bridge interactions. **(c-e)** Occurrence of π-π, π-cation, and π-anion interactions over time.

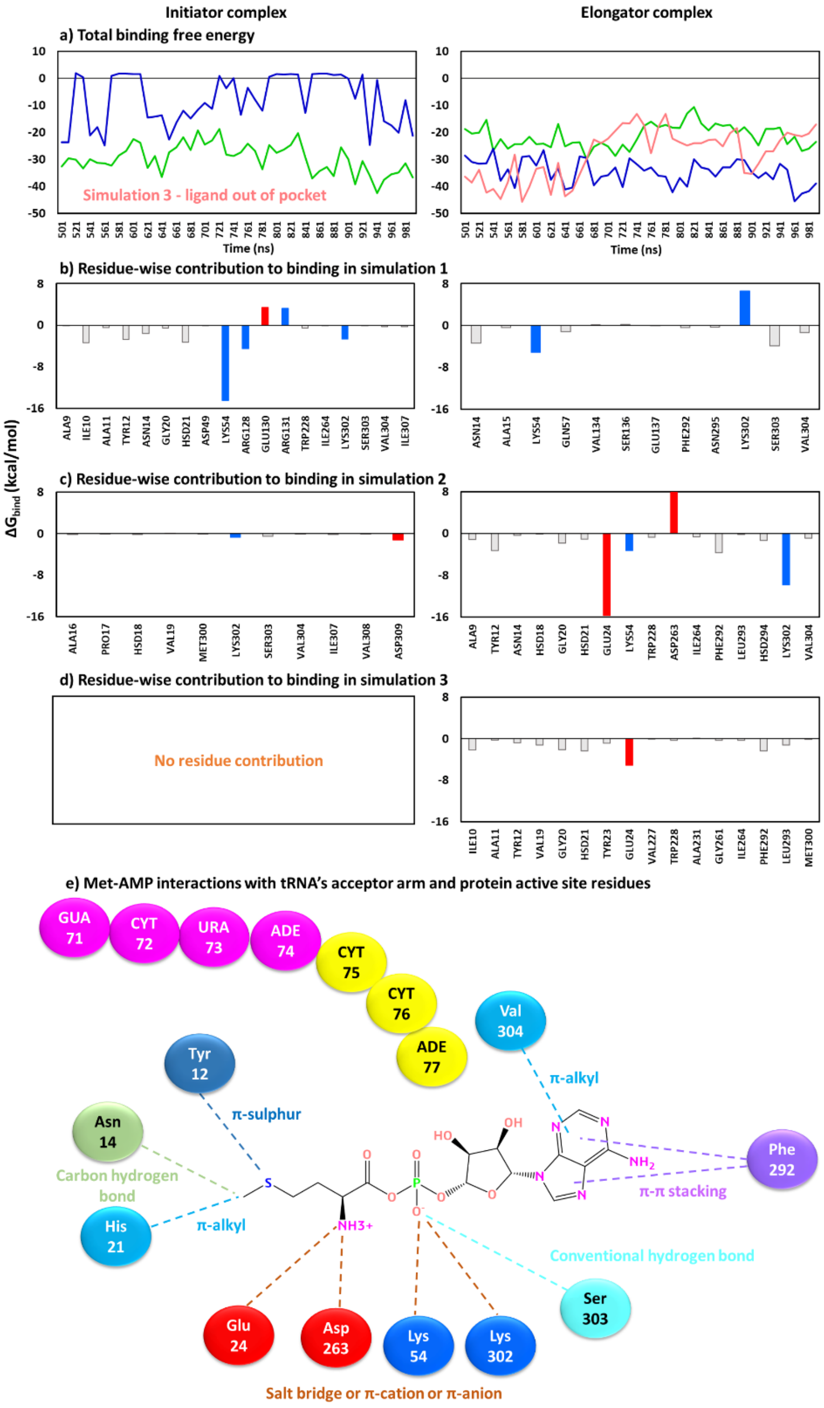


**Figure S23. Binding energy for protein-ligand binding in initiator and elongator complexes. (a)** Total binding free energy ($\Delta G_{bind}$) over time for three simulations. **(b-d)** Residue-wise contribution to binding energy over three simulations, highlighting residues involved in ligand binding (positively charged residue - blue, negatively charged - red and neutral - light gray). **(e)** Important interactions of ligand (Met-AMP) observed during simulations of both complexes.

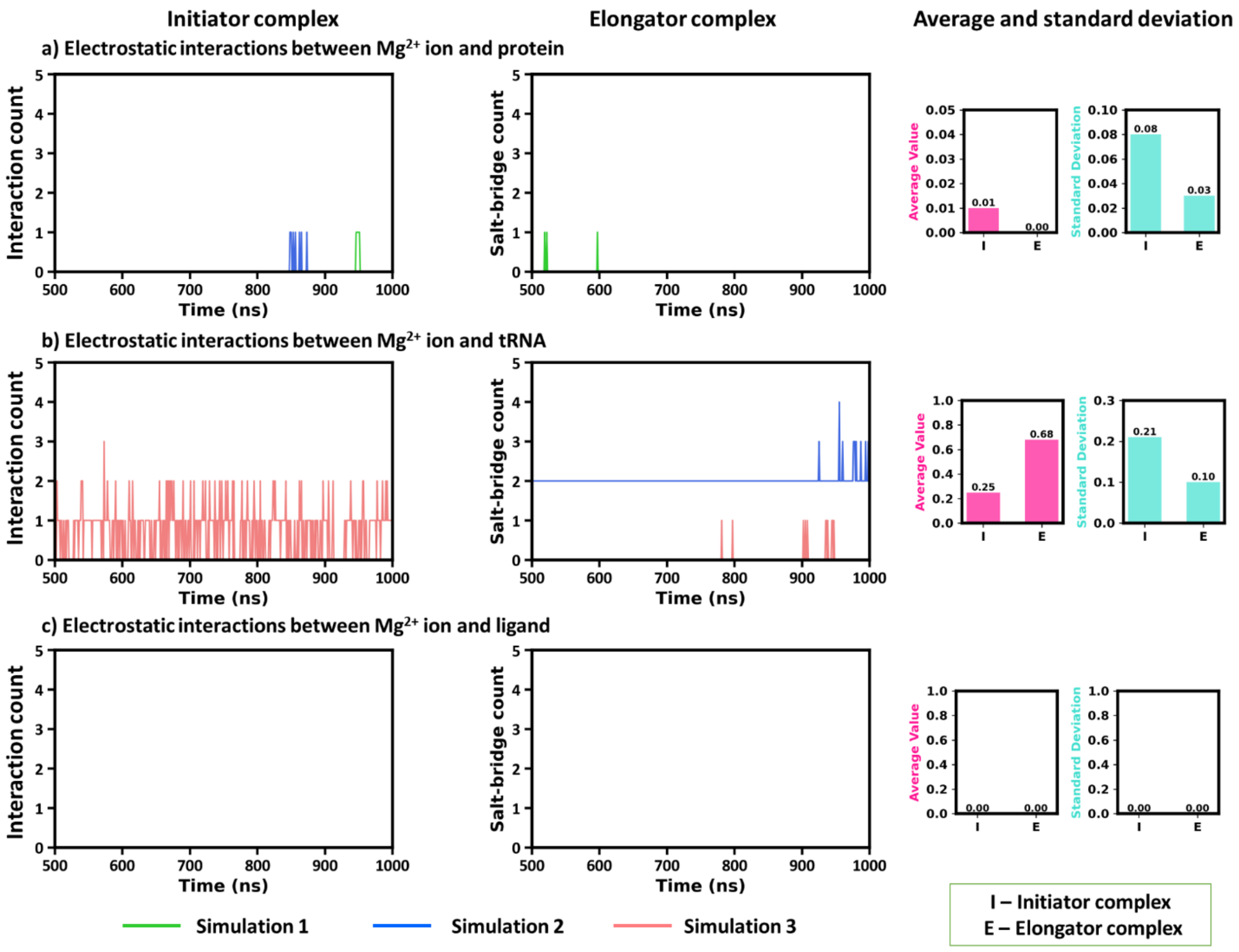


**Figure S24. Electrostatic interactions with $Mg^{2+}$ ion in initiator (I) and elongator (E) complexes.** Ionic interactions formation between $Mg^{2+}$ and **(a)** protein, **(b)** tRNA, and **(c)** ligand, respectively. Bar plots representing average interaction counts and standard deviations.

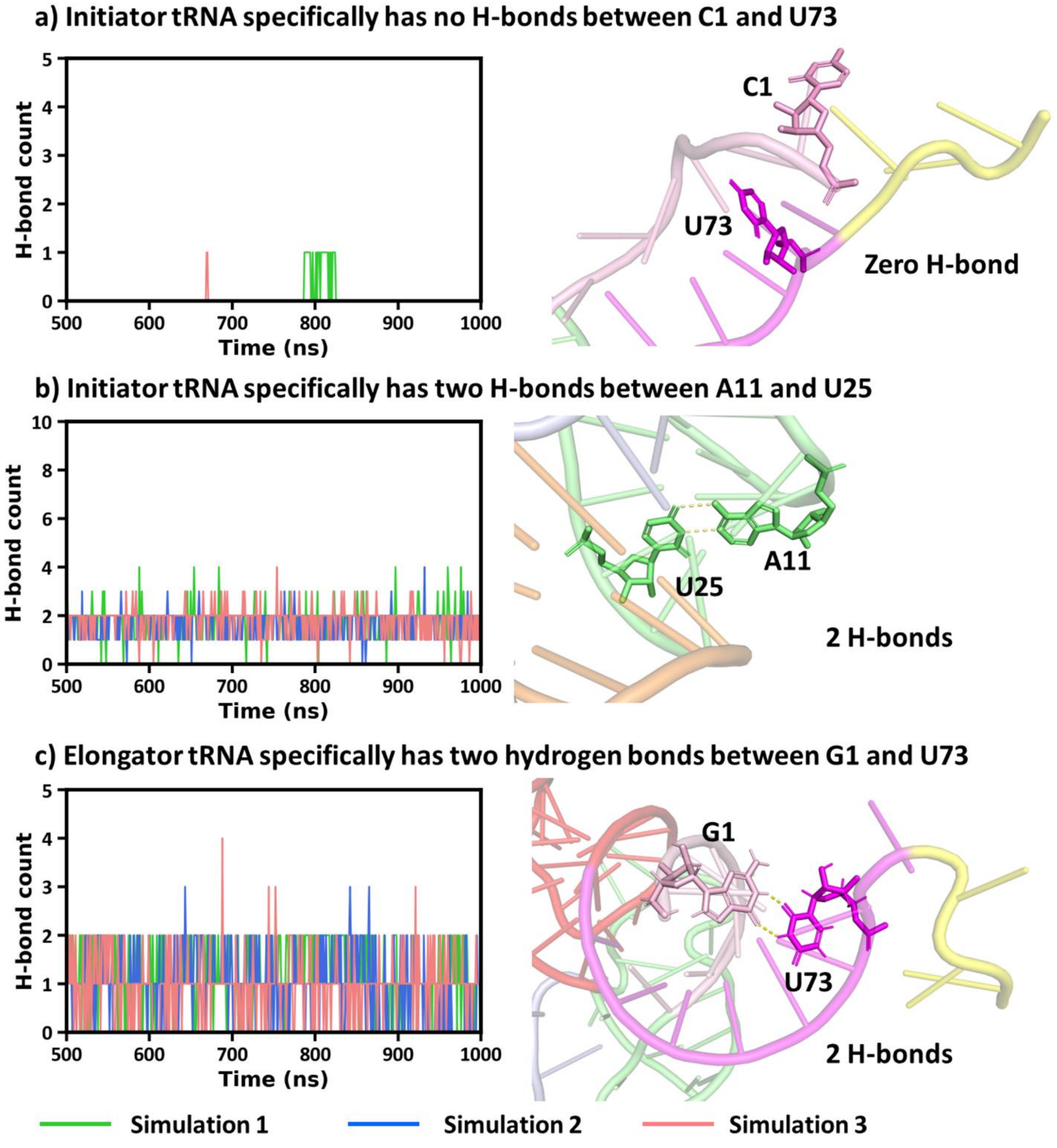


**Figure S25. Base-specific hydrogen bonding patterns in initiator and elongator tRNAs. (a)** Initiator tRNA lacks hydrogen bonds between C1 and U73, indicating an open end. **(b)** A stable A11-U25 base pair is observed in initiator tRNA with consistent hydrogen bonding. **(c)** Elongator tRNA shows a G1-U73 pair stabilized by two hydrogen bonds, reflecting canonical base pairing at the acceptor stem.

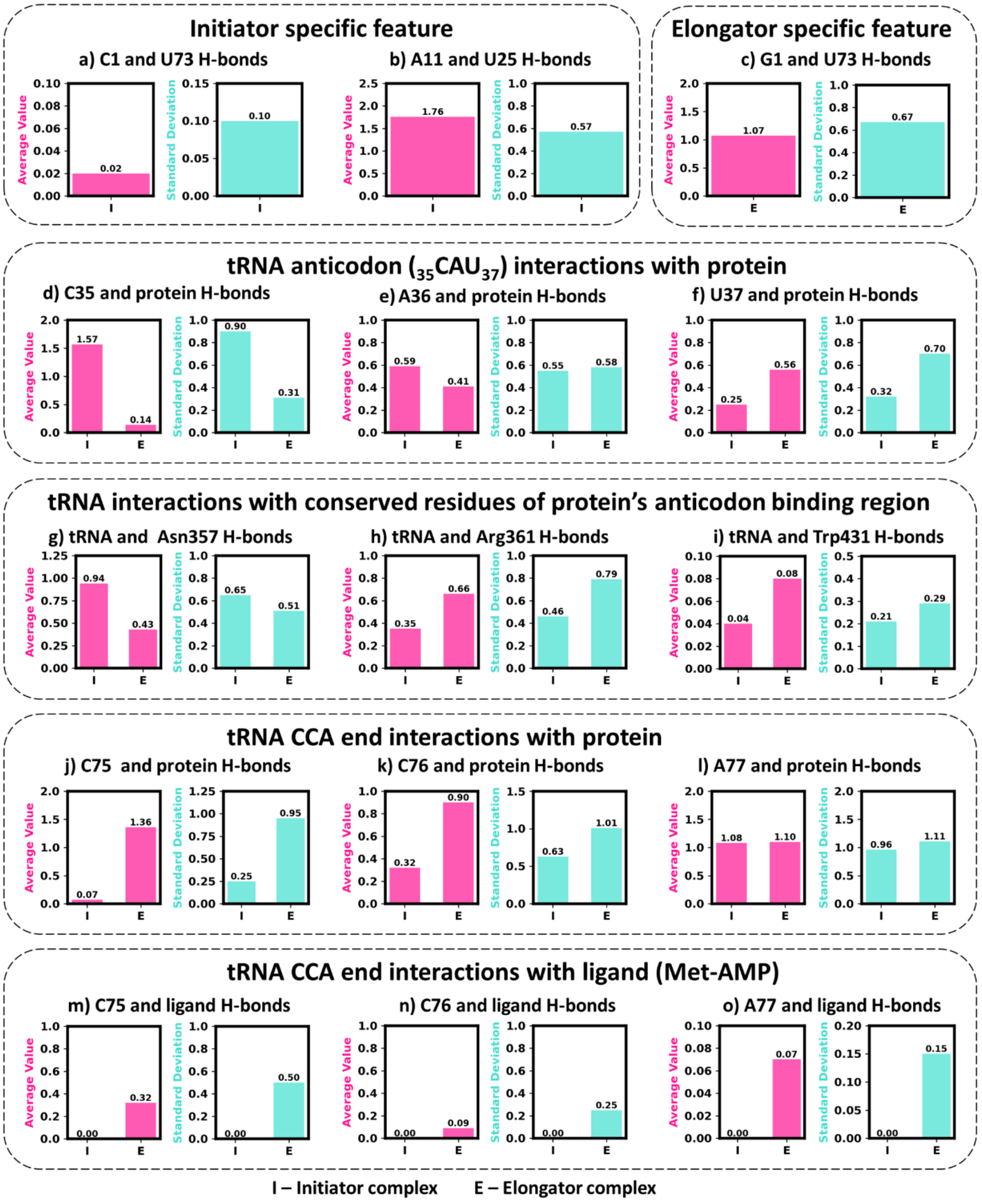


**Figure S26. Average and standard value comparison of H-bond counts for tRNA centric interactions. (a) - (b)** Characteristic interactions of initiator tRNA, **(c)** characteristic interactions of elongator tRNA, **(d) - (f)** anticodon (CAU) interactions with protein residues, **(g) - (i)** tRNAs interactions with conserved protein residues (Asn357, Arg361, Trp431), **(j) - (l)** CCA end of tRNA interactions with protein residues and **(m) - (o)** CCA end of tRNA interactions with ligand.

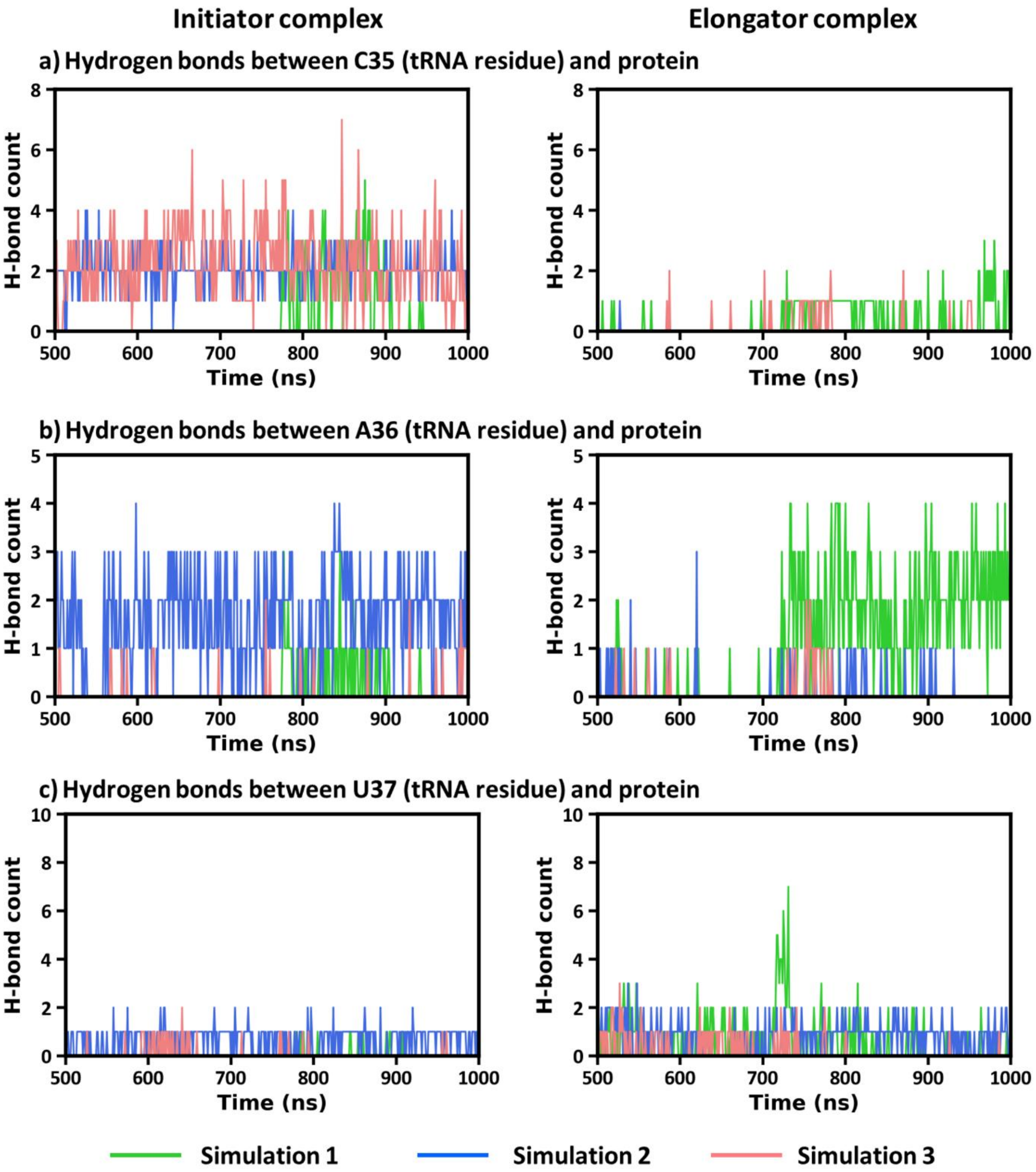


**Figure S27. Hydrogen bonding interactions formed between CAU (35-37) anticodon and protein residues during the simulations.** Interaction of **(a)** C35 nucleotide, **(b)** A36 nucleotide and **(c)** U37 nucleotide with protein.

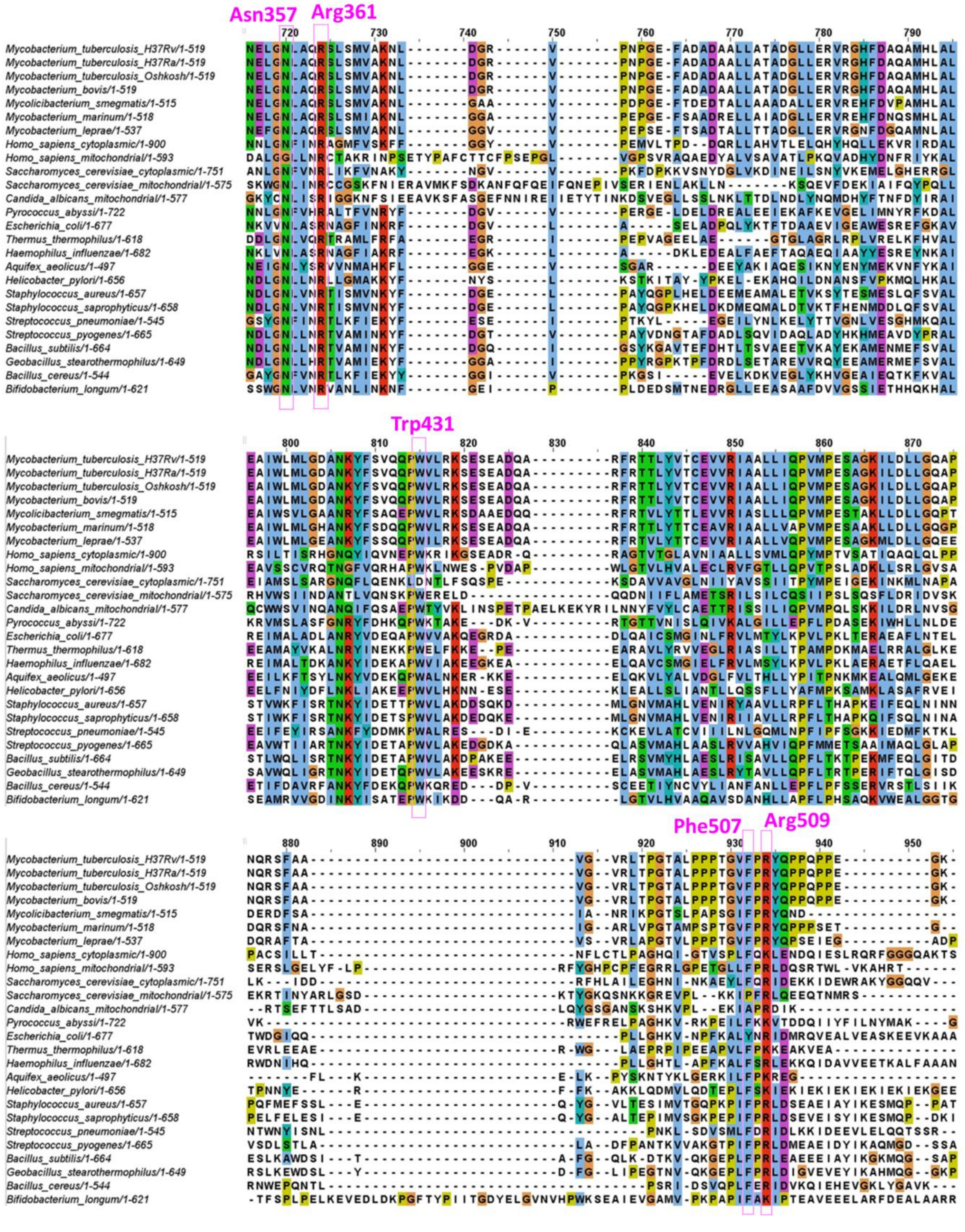


**Figure S28. Conserved residues in anticodon domain of MetRS (shown in purple) from 26 different species.**

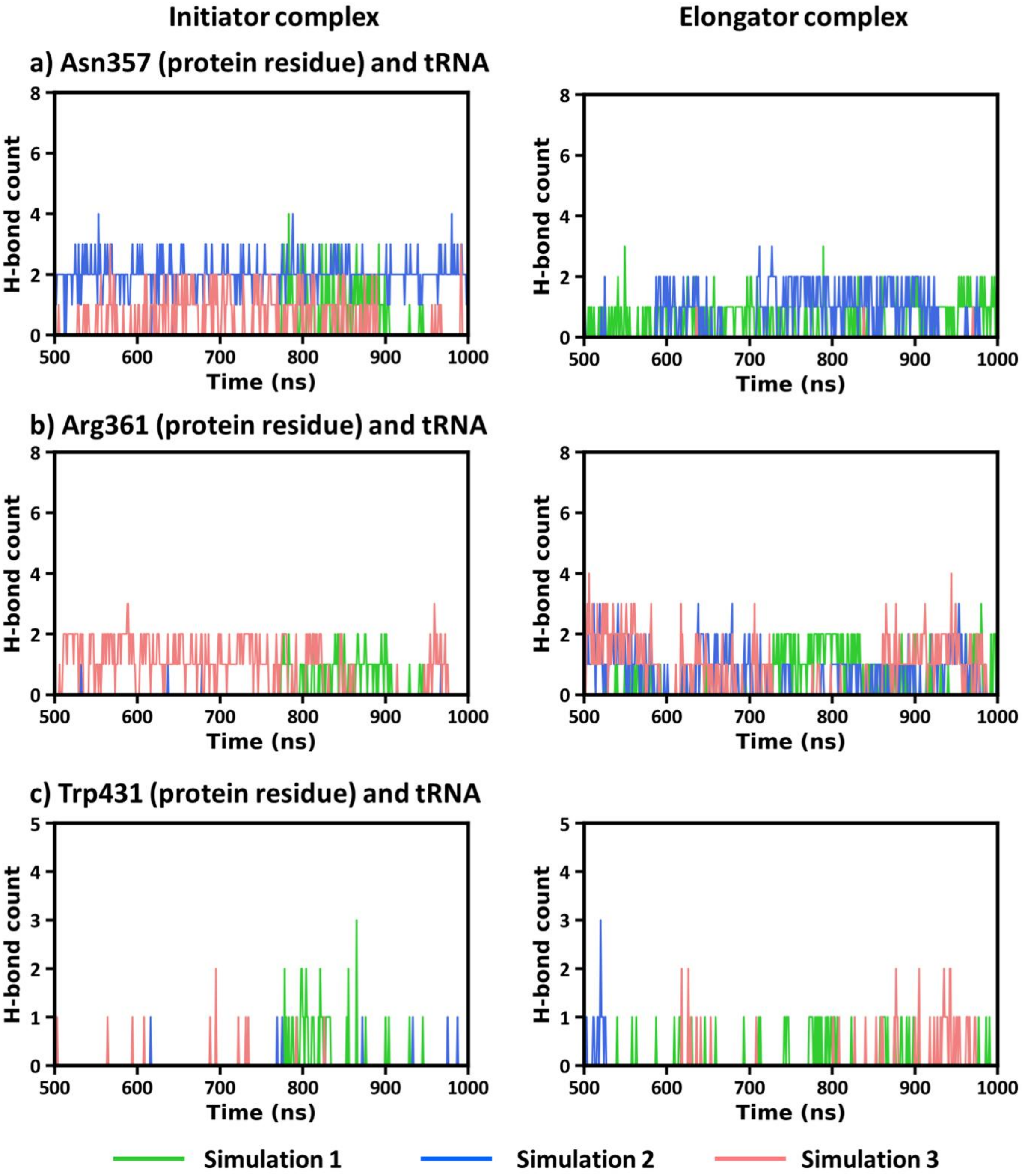


**Figure S29. Hydrogen bonding between three conserved protein residues of anticodon domain and tRNA in initiator and elongator complexes.** Hydrogen bond formed by **(a)** Asn357, **(b)** Arg361 and **(c)** Trp431 with tRNA.

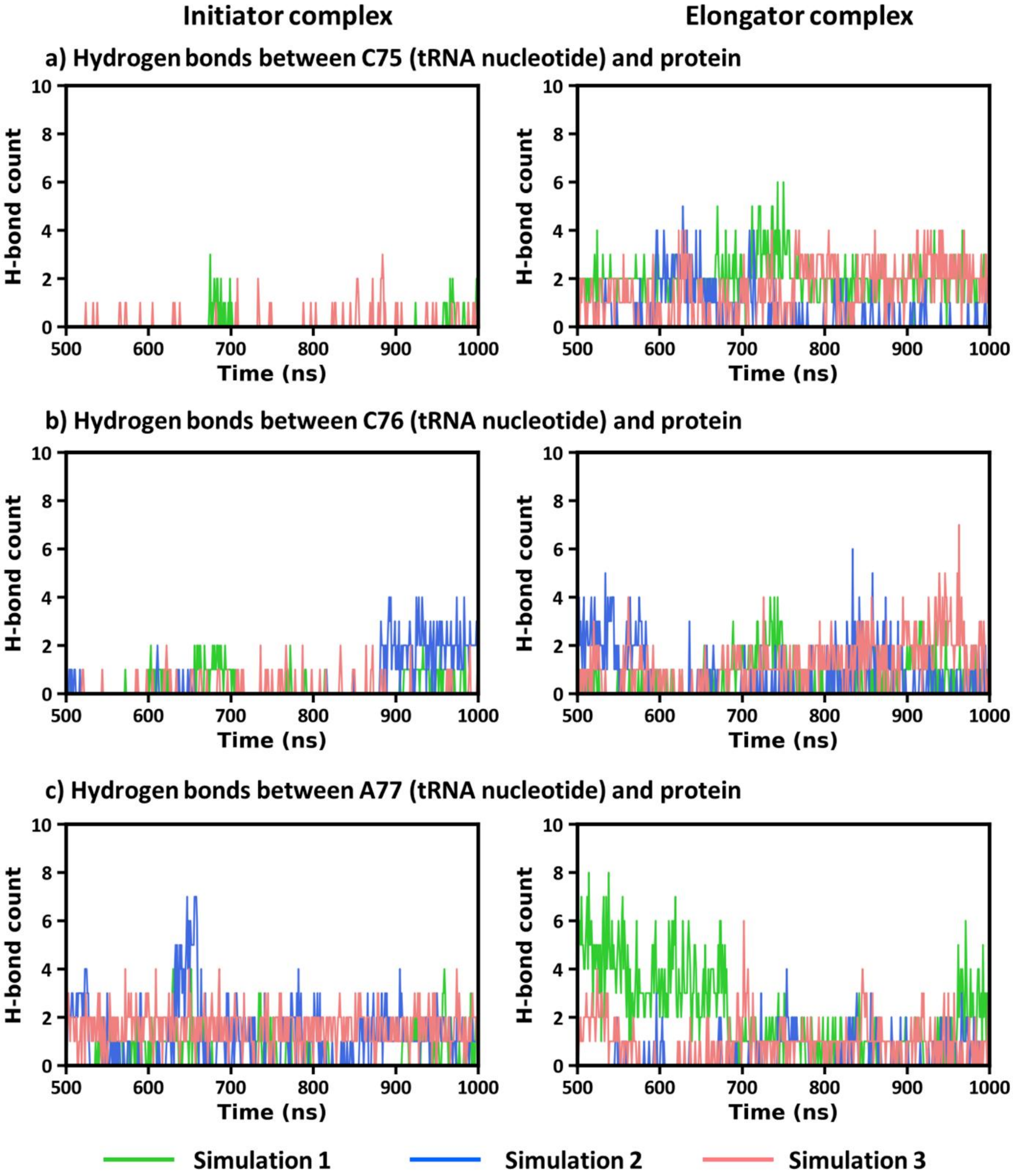


**Figure S30. Hydrogen bond interactions formed by CCA (75-77) end of tRNA with protein residues.** Interaction of **(a)** C75 nucleotide, **(b)** C76 nucleotide and **(c)** A77 nucleotide with protein.

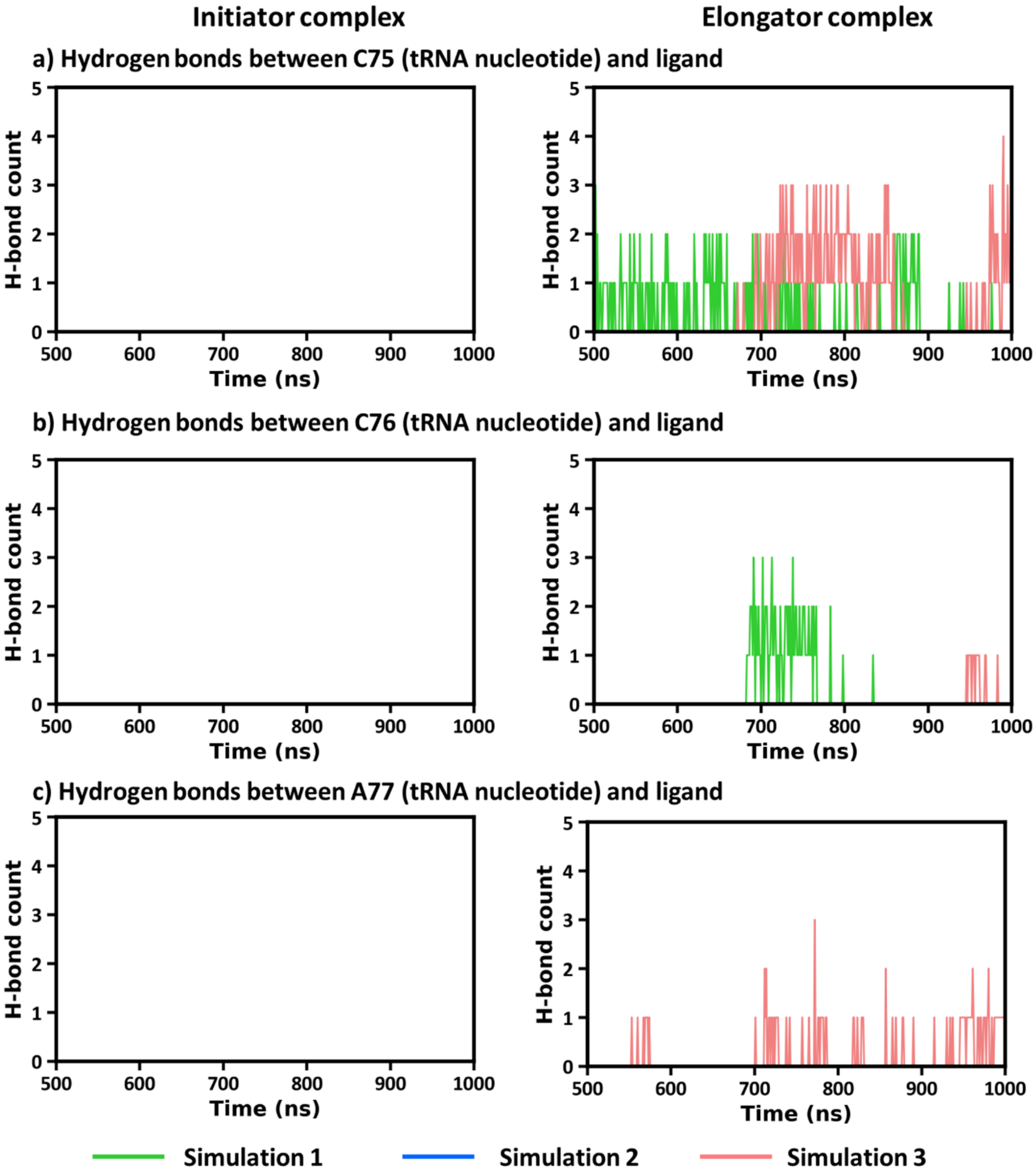


**Figure S31**. **Hydrogen bonding interactions formed by CCA end (75-77) with ligand (Met-AMP).** Interaction of **(a)** C75 nucleotide, **(b)** C76 nucleotide and **(c)** A77 nucleotide with ligand.

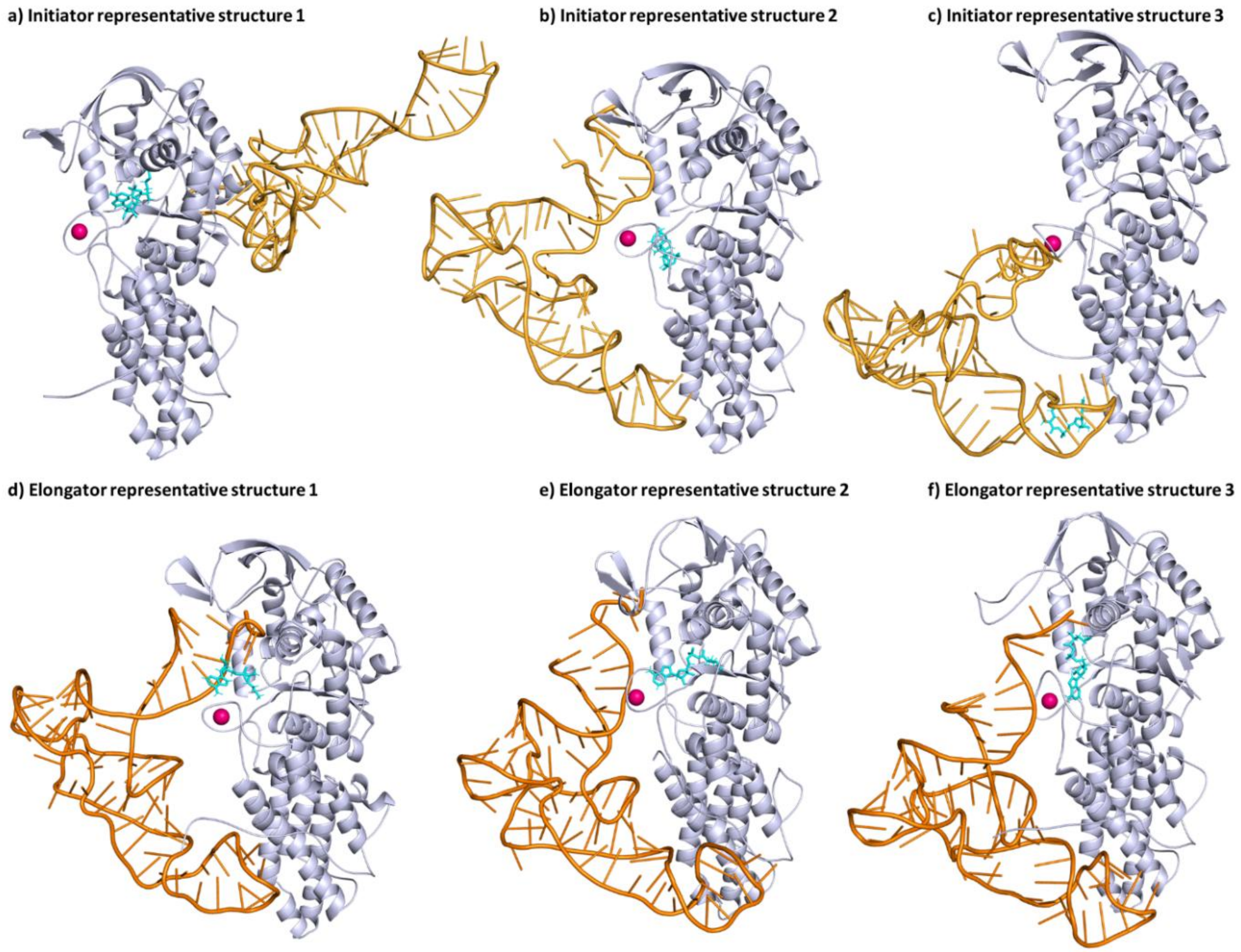


**Figure S32. Representative structures from each simulation run of initiator and elongator tRNA. (a-c)** initiator tRNA complex. **(d-f)** elongator tRNA complex.

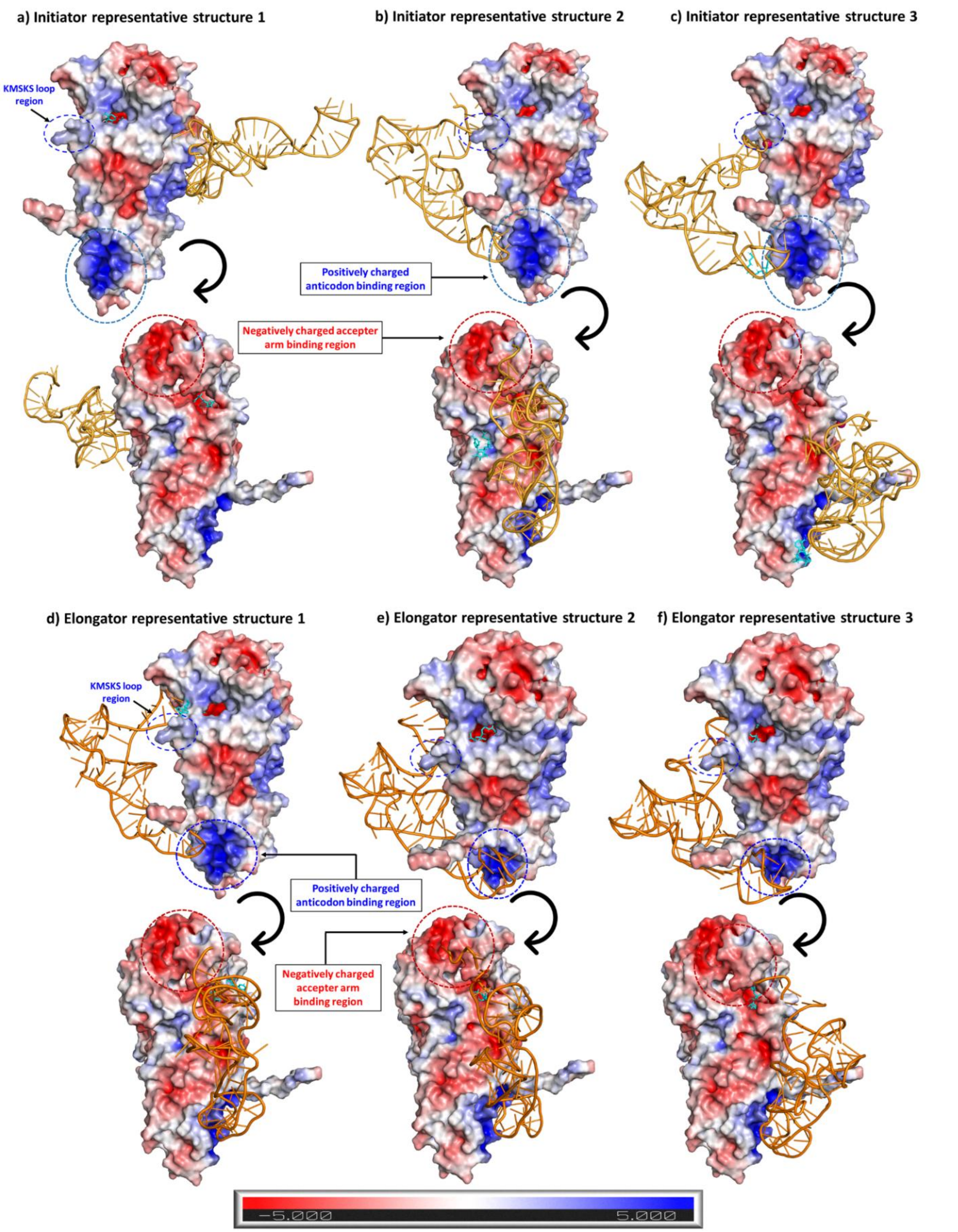


**Figure S33. Electrostatic potential surface of representative structures. (a-c)** initiator tRNA complex. **(d-f)** elongator tRNA complex.

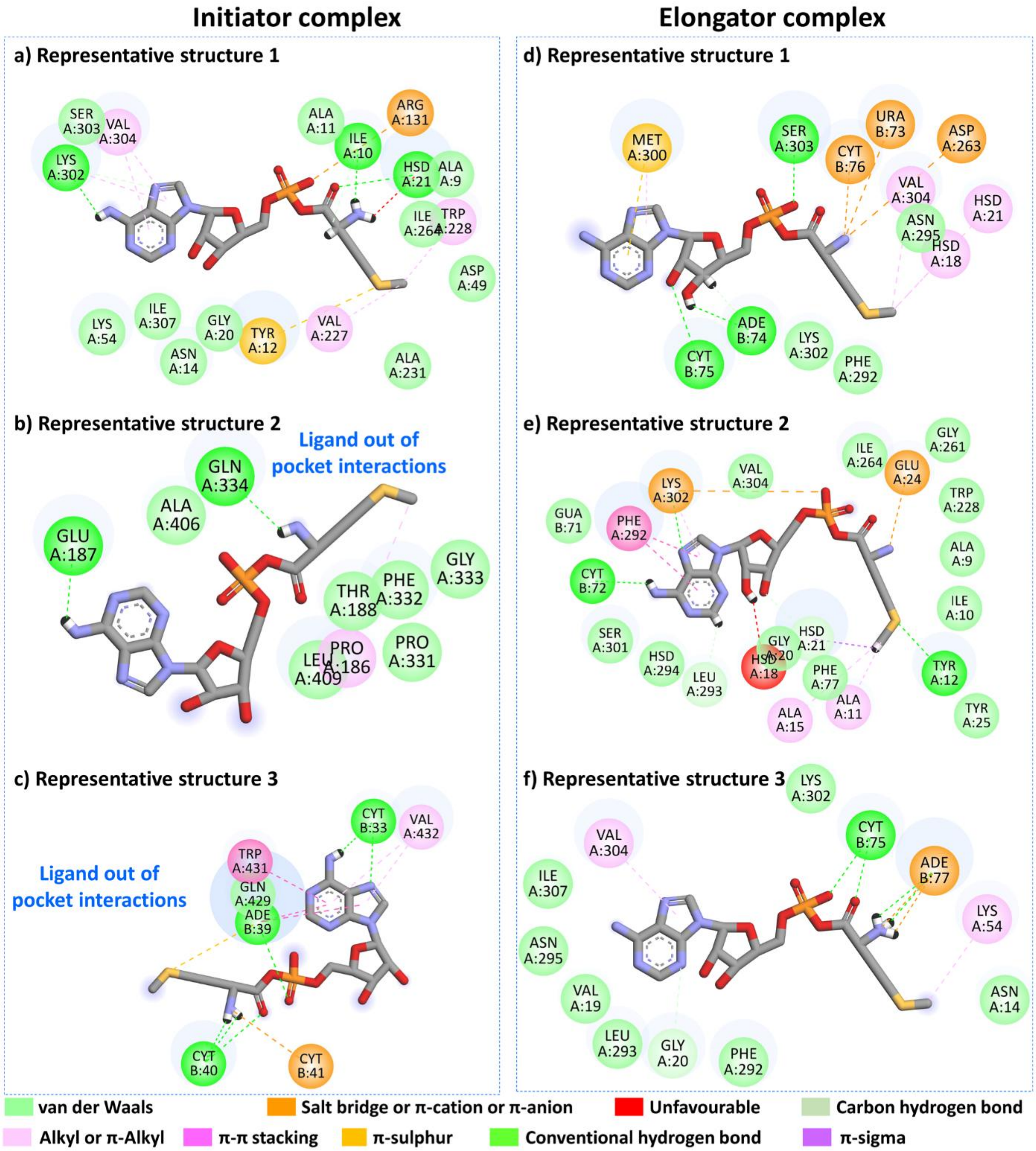


**Figure S34. Interactions observed between protein residues, ligand and tRNA nucleotides in representative structures. (a-c)** initiator and **(d-f)** elongator complex**.**

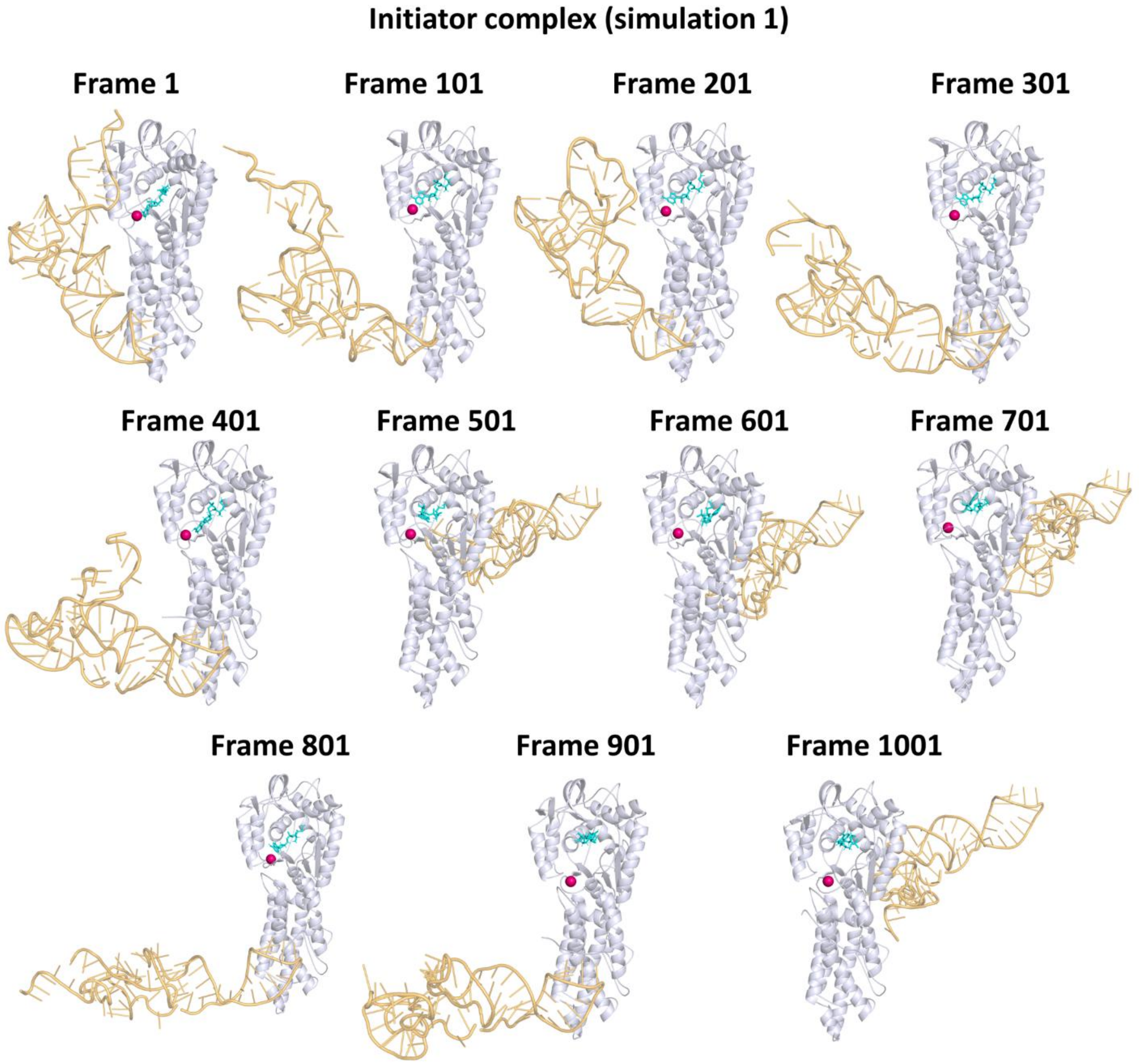


**Figure S35. Trajectory snapshots from simulation 1 of the initiator complex.** Frame 1 corresponds to 0 ns, frame 101 to 100 ns, and at the end frame 1001 to 1000 ns.

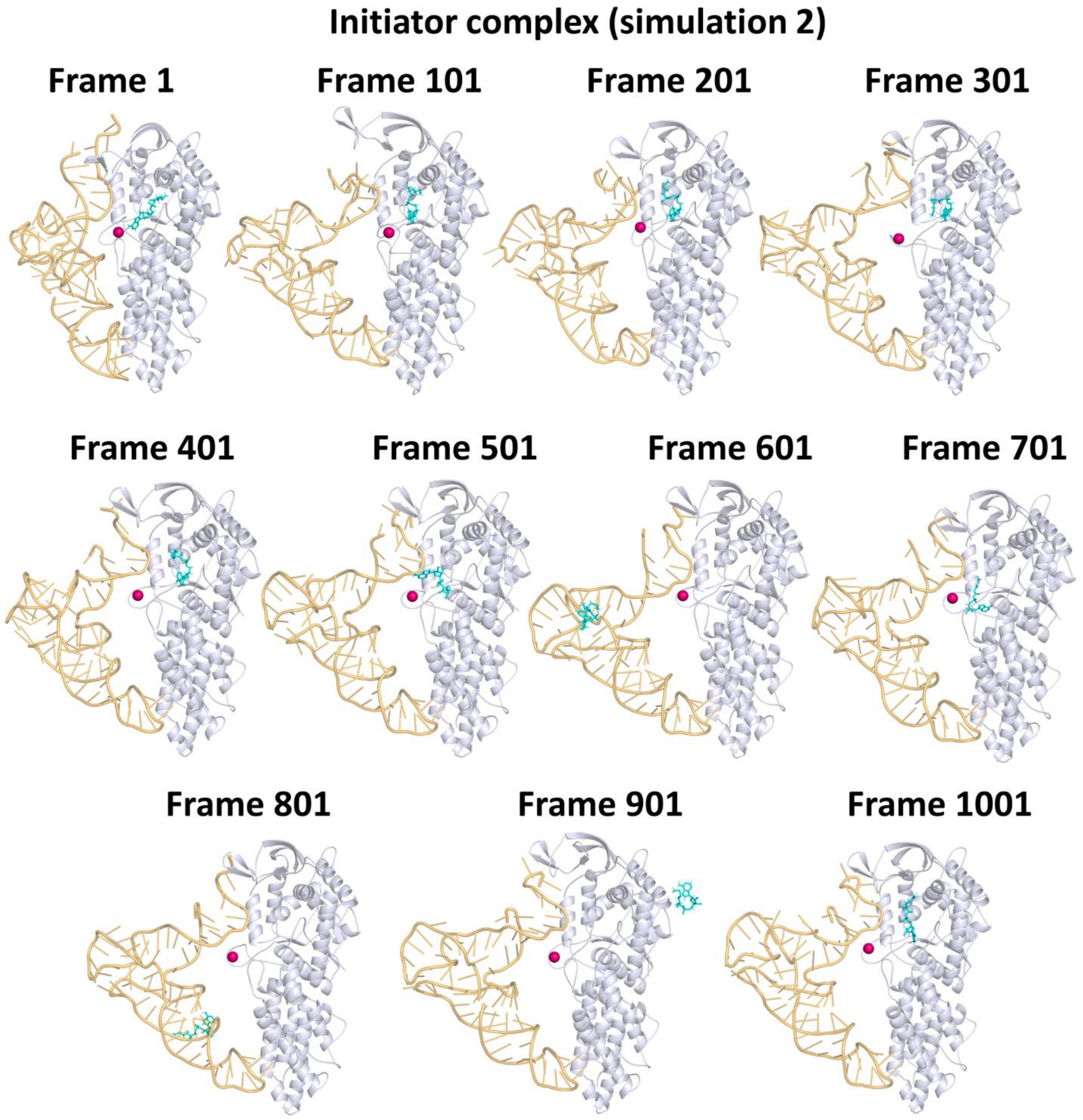


**Figure S36. Trajectory snapshots from simulation 2 of the initiator complex.** Frame 1 corresponds to 0 ns, frame 101 to 100 ns, and at the end frame 1001 to 1000 ns.

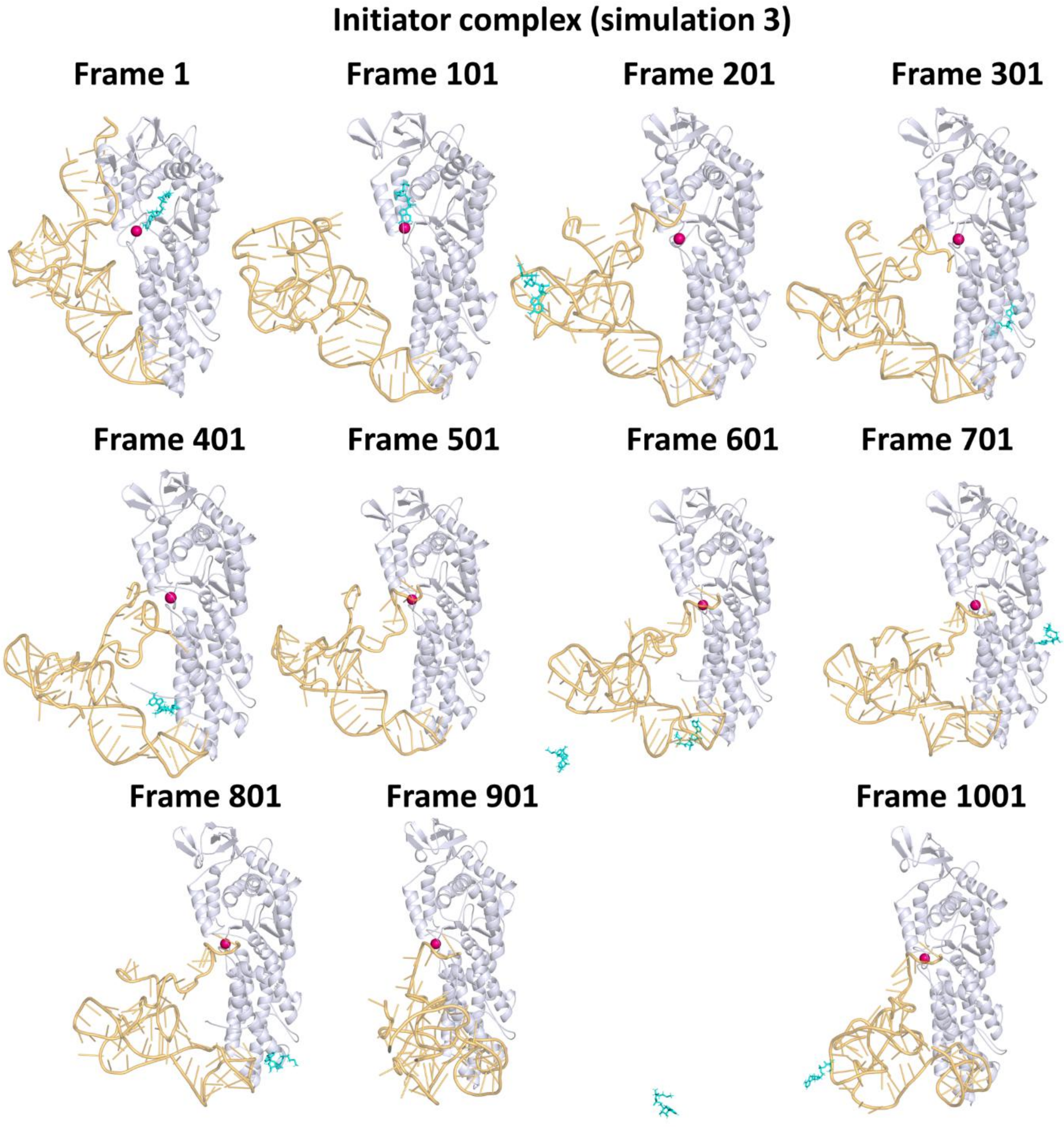


**Figure S37. Trajectory snapshots from simulation 3 of the initiator complex.** Frame 1 corresponds to 0 ns, frame 101 to 100 ns, and at the end frame 1001 to 1000 ns.

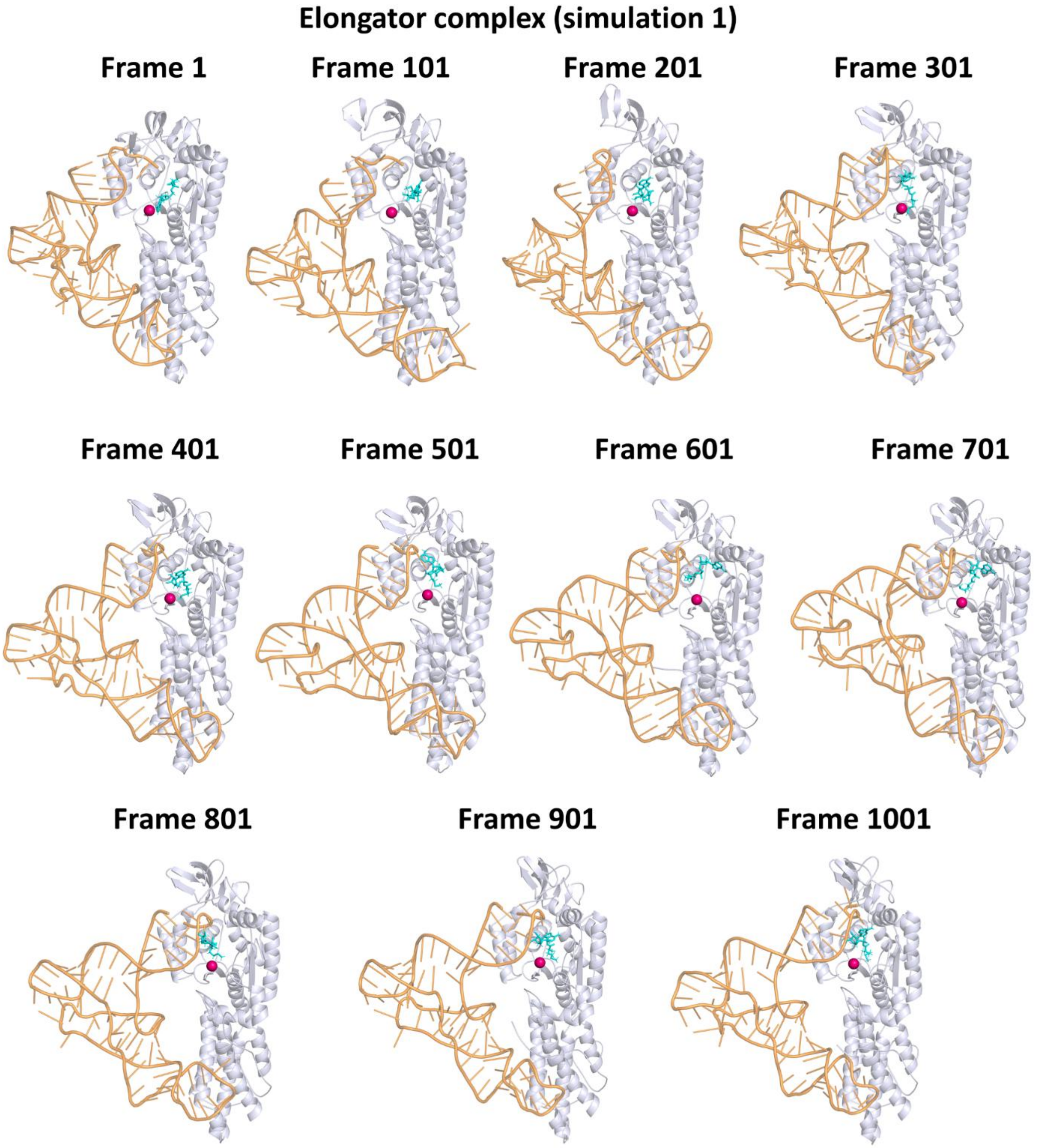


**Figure S38. Trajectory snapshots from simulation 1 of the elongator complex.** Frame 1 corresponds to 0 ns, frame 101 to 100 ns, and at the end frame 1001 to 1000 ns.

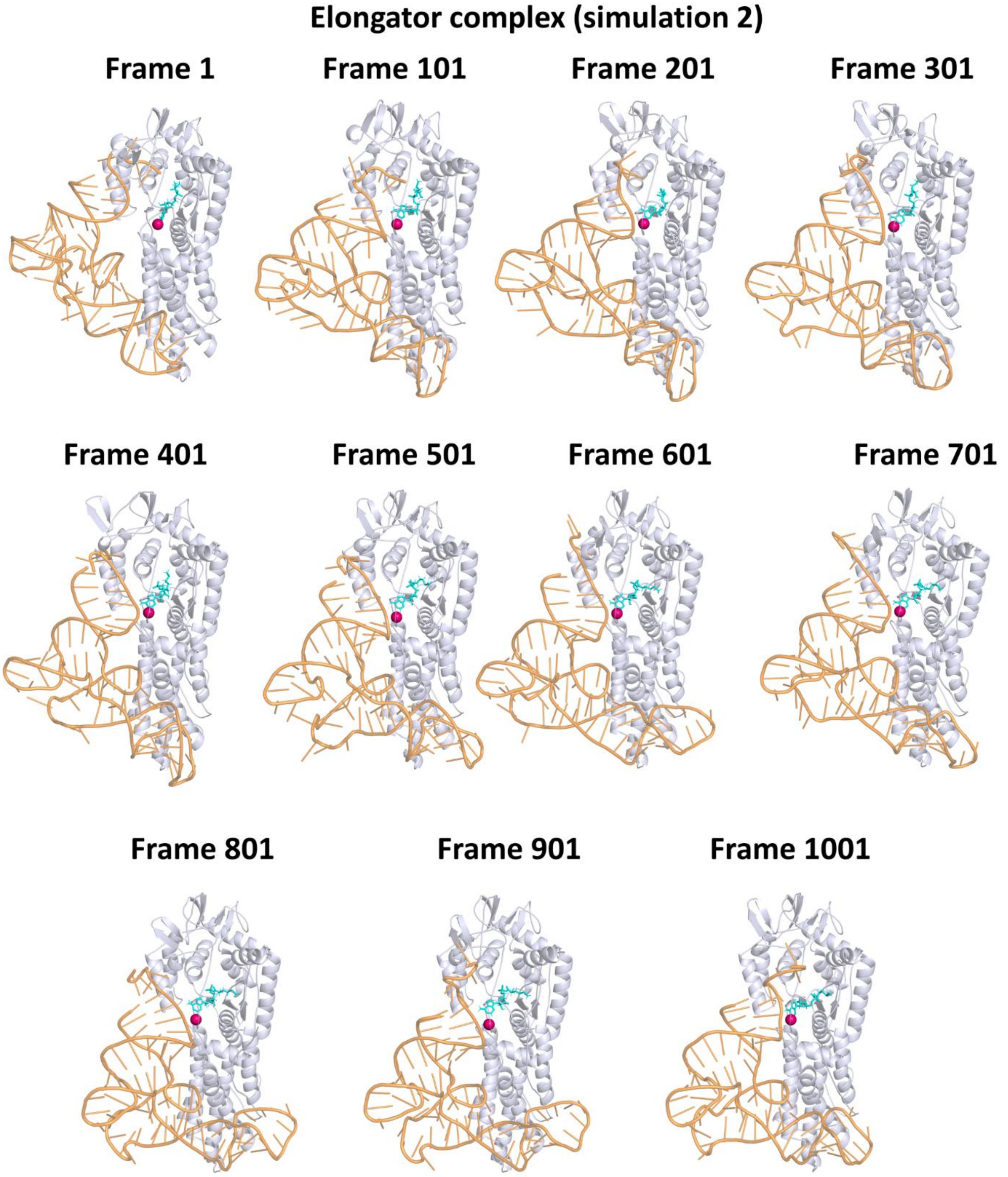


**Figure S39. Trajectory snapshots from simulation 2 of the elongator complex.** Frame 1 corresponds to 0 ns, frame 101 to 100 ns, and at the end frame 1001 to 1000 ns.

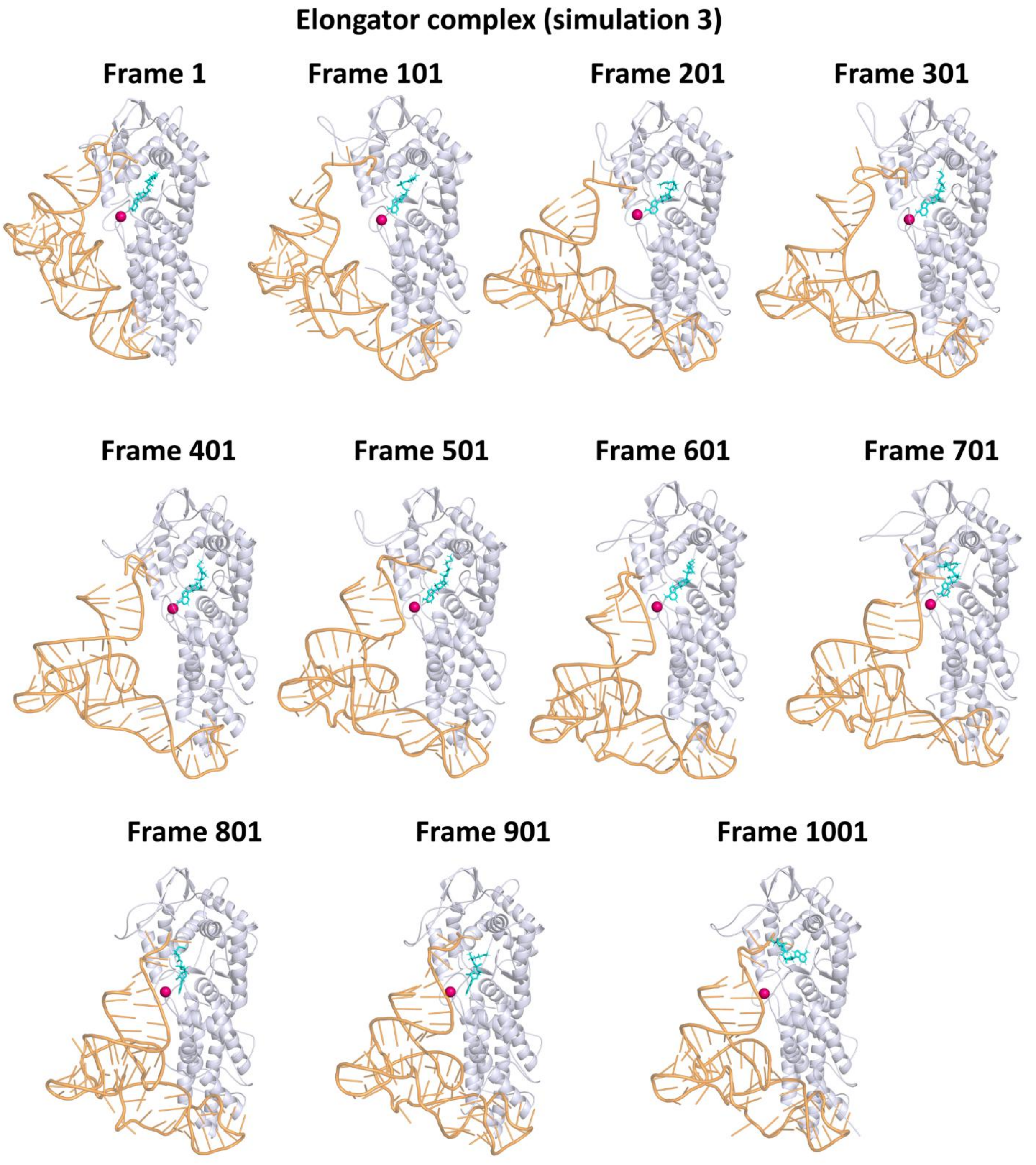


**Figure S40. Trajectory snapshots from simulation 3 of the elongator complex.** Frame 1 corresponds to 0 ns, frame 101 to 100 ns, and at the end frame 1001 to 1000 ns.
